# Transcriptional landscape of direct reprogramming toward the hematopoietic lineage

**DOI:** 10.1101/2024.08.26.609589

**Authors:** Jillian Cwycyshyn, Cooper Stansbury, Sarah Golts, Hyunsu Lee, Joshua Pickard, Walter Meixner, Indika Rajapakse, Lindsey A. Muir

## Abstract

Direct reprogramming of human fibroblasts into hematopoietic stem cells (HSCs) offers a promising strategy for generating autologous cells to treat blood and immune disorders. Current protocols are limited by low efficiency and insufficient tools for evaluating reprogramming outcomes. Although functional assays are the standard for confirming cell identity, they require fully reprogrammed cells, limiting their utility during protocol development. To address this, we assembled a single-cell transcriptomic reference atlas of hematopoietic reprogramming and tested an algorithmically-predicted transcription factor recipe for HSC induction. Long-read single-cell RNA sequencing of CD34^+^ reprogrammed cells revealed progressive loss of fibroblast identity alongside induction of early hematopoietic and endothelial programs, with reference-atlas benchmarking placing reprogrammed cells in an intermediate transcriptomic state between fibroblasts, endothelial cells, and HSCs. Isoform-level analysis further revealed transcriptional remodeling not captured by gene-level analyses. This experimental-computational framework offers a generalizable strategy for characterizing partially reprogrammed states and guiding optimization of reprogramming protocols.

## INTRODUCTION

The generation of hematopoietic stem cells (HSCs) through cell reprogramming would enable novel treatments for hematopoietic disorders. Current reprogramming strategies fall into two categories: direct reprogramming between somatic cell types, and differentiation of induced pluripotent stem cells (iPSCs) into cell types of interest. While the latter approach has achieved cells that have some functional properties of native HSCs^1^, direct reprogramming offers notable advantages for clinical translation. An important consideration for clinical applications is the use of autologous initial cells, which can circumvent the immunological incompatibility and elevated risk of mortality associated with allogeneic transplantation^2^. Since direct reprogramming does not require a pluripotent intermediate, has fewer target cell states to achieve and validate, and has a lower risk of tumorigenesis, this approach may be more efficient for generating autologous cells from a patient sample^3–7^. However, direct fibroblast-to-HSC reprogramming is limited by low conversion rates and has not yet produced cells that have durable engraftment potential^8–15^.

Assays that test cell function are the gold-standard for evaluating whether reprogramming cells have sufficiently adopted the target cell phenotype. While functional assays aid validation, they require cells to be fully reprogrammed, and therefore have limited utility until after an efficient direct reprogramming protocol is established. Better methods for evaluating reprogramming cells prior to this point would help identify successful and efficient paths of reprogramming. For example, cells that nearly achieve a target state might need only minimal intervention to fully reprogram, and methods that identify these cells would enable new protocols to be developed more quickly.

Single-cell transcriptomics has vastly improved the characterization of phenotype transitions such as differentiation in the hematopoietic lineage in mice^16–18^ and humans^19–21^. Similar approaches have been used to examine heterogeneity^1^ and identify intermediate or off-target states in reprogramming^22^. However, the extent to which reprogrammed cells and their gene expression dynamics compare to native cell types remains challenging to quantify. Comprehensive single-cell atlases can provide a powerful framework for placing incompletely understood or engineered cell types into the context of well-characterized phenotypes^23–25^, but this approach has not yet been used for characterizing cell states in direct reprogramming to HSCs.

Here, we construct and evaluate the utility of a single-cell transcriptomic reference atlas spanning hematopoietic, endothelial, stromal, and engineered cell types for evaluating direct fibroblast-to-HSC reprogramming of human cells. Our approach includes testing a novel combination of transcription factors (TF) —GATA2, GFI1B, FOS, REL, and STAT5A—that were predicted by an algorithmic framework^26^. We profile the resulting CD34^+^ cells using long-read single-cell RNA sequencing (scRNA-seq) and characterize several reprogramming intermediates within the context of our reference atlas. Lastly, we quantify gene- and isoform-level diversity in initial fibroblasts, reprogrammed cells, and native HSCs from bone marrow. Through this integrated experimental-computational approach, we provide a generalizable strategy that can be used to guide and optimize protocols for direct reprogramming.

## RESULTS

### Construction of an atlas for hematopoietic cell reprogramming

Standard analysis of scRNA-seq data relies on cluster annotation, in which transcriptionally similar cells are grouped and assigned cell type identities based on marker gene expression for expected cell types^27,28^ This approach works well for native cell types with well-characterized transcriptional profiles, but is less effective in reprogramming contexts where induced cells may occupy poorly defined intermediate states that do not clearly match existing cell types^29^. Here, we sought to establish a reference framework that aids assessment of cell reprogramming outcomes, serving as a ground truth for assessing similarity, divergence, and partial identity across reprogrammed cell states.

We constructed a single-cell reference atlas drawn from 270 publicly available datasets covering over one million human hematopoietic, immune, and stromal cells (Figure S1, Data S1). To mitigate technical variability, data were uniformly preprocessed and harmonized using single-cell variational inference (scVI) and conditioned on standardized cell type annotations provided by the source datasets^30^. Source cell type annotations were consolidated into major cell compartments (e.g., fibroblast, endothelial, HSCs), which serve as the primary units of analysis throughout this study. To visualize gene expression patterns in the reference atlas, we generated a low-dimensional *t*-distributed stochastic neighbor embedding (t-SNE) from the harmonized latent space (Figures 1A-B)^31^.

**Figure 1:**
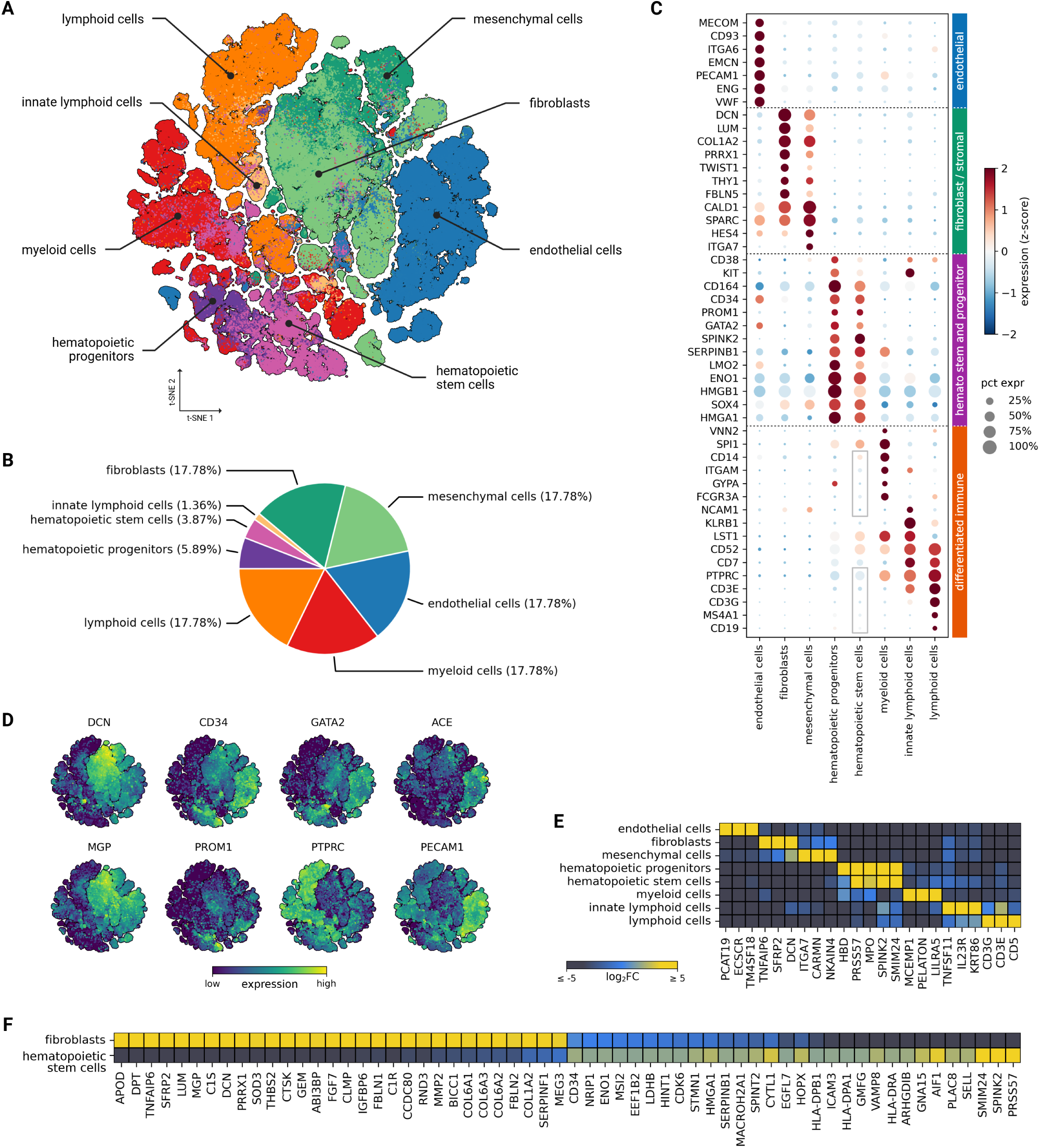
A reference atlas for human hematopoietic cell reprogramming. **(A)** t-SNE embedding of cell types relevant to fibroblast-to-HSC reprogramming (1.12 million cells). **(B)** Proportions of major cell compartments in (A). **(C)** Average expression (z-score) of marker genes for endothelial, fibroblast/stromal, hematopoietic stem and progenitor, and differentiated immune cells in the major cell compartments in (A). Dot size represents the percentage of cells expressing a given gene. Gray boxes indicate genes encoding standard lineage-negative markers for HSCs. **(D)** Expression of fibroblast, hematopoietic, and endothelial marker genes projected onto the t-SNE coordinates from (A). **(E)** Top three DEGs per major cell compartment (detected in ≥ 15% of cells per group). Color reflects *log*_2_ fold changes relative to the other cell compartments. **(F)** Top 30 DEGs detected in ≥ 35% of fibroblasts and hematopoietic stem cells. See also Figures S1-S7, Tables S1-S2, Data S1.

To validate the integrated reference atlas, we evaluated lineage-specific gene expression and differentially expressed genes (DEGs) across the major cell compartments. Lineage-specific gene expression was consistent with annotations for fibroblasts and mesenchymal cells (*DCN*, *COL1A2*, *CALD1*), endothelial cells (*ACE*, *PECAM1*, *ENG*), hematopoietic stem and progenitor cells (*CD34*, *GATA2*, *CD164*), myeloid cells (*SPI1*, *CD14*, *ITGAM*), and lymphoid cells (*CD7*, *CD3*, *CD52*) (Figures 1C-D and Figure S2), while lineage-specific marker transcripts were low or absent in HSCs (Figure 1C, gray boxes). Likewise, DEGs between the major cell compartments were consistent with established marker sets and TFs (Figures 1EF, Figure S3, Tables S1 and S2)^28,32^. Hematopoietic markers (*MYB*, *LMO2*, *HOXA9*, *SPINK2*) and TFs (*HMGA1*, *MSI2*) were enriched in the hematopoietic compartment (Figure 1C, Figures S3, S4, S5, S6), fibroblasts and mesenchymal cells expressed extracellular matrix genes (*LUM*, *FBLN1*, *COL6A1*) and mesenchymal regulators (*PRRX1*, *TWIST1*) (Figures 1C, F and Figure S8), and the endothelial compartment expressed endothelial markers (*VWF*, *EMCN*, *ECSCR*) and TFs (*ETS1*, *EPAS1*, *SOX17*) and was enriched for transcripts related to angiogenesis, immune signaling, and vascular transport (Figures 1C, E and Figures S3 and S7).

Together, these data establish a comprehensive transcriptomic reference atlas for evaluating the outcomes of fibroblast-to-HSC direct reprogramming with respect to native cell types.A data-guided TF recipe produces CD34^+^ cells

In prior work, we developed a data-guided control (DGC) algorithm to predict candidate combinations of TFs for direct reprogramming^26^. Here, we applied DGC to generate candidate TFs for fibroblast-to-HSC reprogramming, with the goal of producing a dataset for evaluation within our reference atlas. TFs previously reported to induce an HSC-like phenotype in human fibroblasts (3TF: GATA2, GFI1B, and FOS)^10^ ranked among the top DGC candidates. Two additional top-ranking TFs, STAT5A and REL, were selected for inclusion in a 5TF reprogramming protocol.

Human neonatal fibroblasts were transduced with inducible Tet-ON lentiviral vectors encoding the 5TF recipe (Figure 2A). On day 0, doxycycline (Dox) was added for 48 hours to induce TF expression. On day 3, transcripts from all five TFs were detected by qRT-PCR relative to untransduced Dox-treated controls (Figure 2B, Table S3). On day 7, approximately 3% of 5TF-induced cells expressed the hematopoietic marker CD34, representing a significant increase compared to transduced cells not treated with Dox (Figure 2C). Immunostaining confirmed CD34 expression in 5TF-induced cells (Figure 2D). A similar frequency of CD34^+^ cells was observed across independent experiments and variations of these TF combinations (Figure S10). Expression of the hematopoietic surface marker CD143 was also detected, though at lower levels. Other commonly used hematopoietic markers, including CD49f and CD9, were expressed in fibroblasts and therefore were not used to evaluate the reprogramming population.

**Figure 2:**
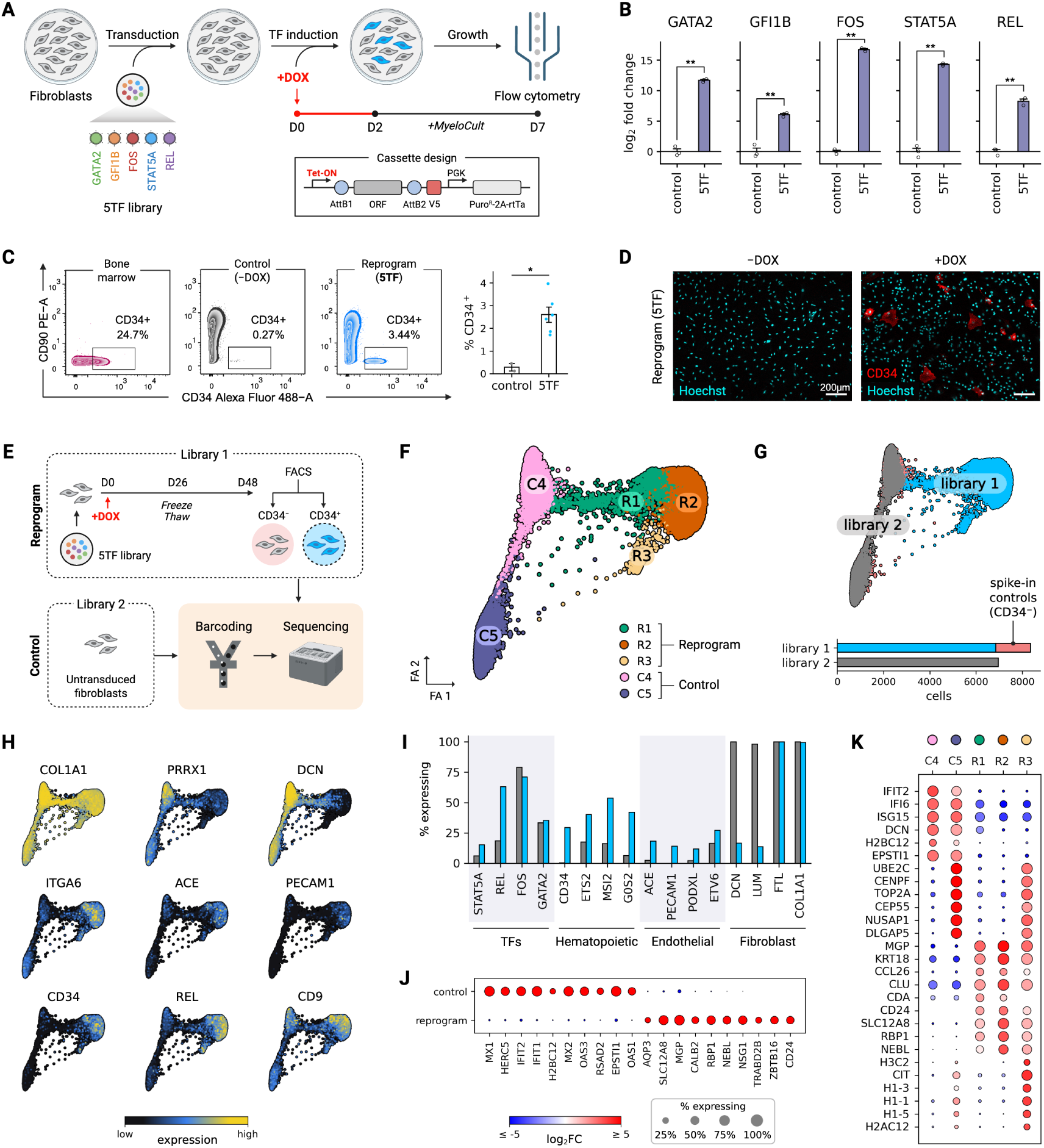
Direct reprogramming of fibroblasts into CD34^+^ cells. **(A)** Overview of the reprogramming workflow and design of the inducible TF expression cassette. **(B)** 5TF expression as measured by qRT-PCR in control and 5TF-induced fibroblasts. Bars show mean ± SEM of −ΔΔCt, normalized to *LDLR* and expressed relative to control (*n* = 3 replicates). **(C)** Left: Flow cytometry plots for bone marrow, uninduced fibroblast controls, and 5TF-induced cells showing CD34. Right: CD34^+^ cells as a percentage of live cells. Bars show mean ± SEM (*n* = 6 replicates for 5TF, *n* = 2 replicates for control). **(D)** Immunofluorescent staining of CD34 in 5TF cells with or without Dox. Scale bar = 200*µm*. **(E)** Overview of scRNA-seq library preparation. Library 1 (reprogram): CD34^+^ 5TF-induced cells with CD34spike-in controls. Library 2 (control): untransduced fibroblasts. **(F)** Force-directed graph of single cells colored by Leiden cluster. **(G)** Graph from (F) colored by library. Cell counts shown below (Library 1: 6, 849 CD34^+^ 5TF-induced cells, 1, 508 CD34spike-in controls; Library 2: 6, 960 cells). **(H)** Normalized expression of select fibroblast, endothelial, and hematopoietic genes on the graph from (F). **(I)** Fraction of cells with nonzero expression for key TFs, hematopoietic, endothelial, and fibroblast genes in cells from both libraries. **(J)** Top protein-coding DEGs between control (C4/C5) and reprogrammed (R1/R2/R3) cells. **(K)** Top protein-coding DEGs per cluster. In (J) and (K), dot color represents log_2_ fold change, and dot size reflects the percentage of cells expressing a gene in a given group. * *p <* 0.05, ** *p <* 0.01 by Welch’s *t*-test (B) or by Mann-Whitney U test (C). See also Figures S10, S11, and S12, and Table S3.

We next performed scRNA-seq on 5TF-induced cells to generate a dataset for evaluation within our reference atlas. Prior to sequencing, CD34^+^ cells were sorted on day 48 to enrich for cells exhibiting durable phenotypic changes (Figure S10D). We refer to this dataset as Library 1. To represent the initial cell state, we generated a second scRNA-seq dataset from untransduced fibroblasts (Library 2). After quality control, cells from both libraries were pooled, harmonized, and clustered, yielding 15,317 cells with three clusters of reprogrammed cells (R1-R3) and two fibroblast control clusters (C4, C5) (Figures 2F-G). As an internal control, spike-in CD34sorted cells from day 48 that were included in Library 1 clustered with fibroblasts from Library 2, confirming their non-reprogrammed identity and demonstrating successful harmonization across libraries (Figure S11). Only fibroblast control cells were used as the baseline population in downstream analyses.

To assess transcriptional identity in reprogramming cells, we examined fibroblast and hematopoietic gene expression across both libraries (Figures 2H-I). Fibroblast-associated genes, such as *PRRX1*, *DCN*, and *LUM*, were broadly reduced in reprogrammed cells, although residual fibroblast identity persisted in a subset of cells (Figures S11E and S12C). This was most apparent in cluster R1, which retained elevated expression of multiple fibroblast-associated genes. In contrast, R2 and R3 exhibited a more pronounced loss of fibroblast markers, though expression of structural genes such as *COL1A1* and *S100A4* remained elevated, indicating incomplete silencing of the fibroblast program. Reprogrammed cells also showed partial induction of hematopoietic genes, including increased expression of *CD34*, *ITGA6*, and *CD9*, as well as a greater proportion of cells expressing genes such as *ETS2*, *MSI2*, and *G0S2* (Figure S12A). Reprogrammed cells simultaneously upregulated endothelial-associated genes, including *ACE*, *PECAM1*, and *PODXL*, consistent with the endothelial-like intermediate previously reported during 3TF reprogramming (Figure S12B)^10^.

Next, we performed differential expression analysis to identify genes globally altered by reprogramming (Figure 2J). Among the most highly upregulated genes were *CD24* and *ZBTB16*, a TF implicated in balancing HSC self-renewal and differentiation^33^. Additional upregulated genes were associated with membrane transport (*AQP3*, *SLC12A8*), cytoskeletal remodeling (*NEBL*), and metabolic regulation (*NSG1*, *CALB2*), suggesting a more plastic, reprogramming-permissive state. In contrast, interferon-stimulated genes, including *MX1*, *MX2*, and *IFIT1*, were reduced, potentially reflecting suppression of a steady-state innate immune program during reprogramming^34^. At the cluster level, interferon-responsive genes and fibroblast markers were enriched in C4/C5 and broadly downregulated across R1-R3 (Figure 2K). Clusters R1 and R2 upregulated remodeling-associated genes (*MGP*, *KRT18*, *CLU*), with R2 exhibiting the strongest induction of progenitor-associated genes (*CD24*, *RBP1*). In contrast, R3 displayed a distinct proliferative and chromatin-remodeling signature characterized by elevated expression of cell cycle genes and histone transcripts (*TOP2A*, *CIT*, *H1-3*).

Together, these results demonstrate that the 5TF recipe converts CD34fibroblasts into transcriptionally distinct CD34^+^ cells. The resulting population co-expressed fibroblast, endothelial, and hematopoietic markers, consistent with a heterogeneous and partially reprogrammed transcriptional state.

### Velocity pseudotime reveals emergence of early hematopoietic signature

The transcriptional heterogeneity observed among 5TF-induced cells suggests that reprogramming produces a spectrum of partially reprogrammed states. We next asked whether these states arise sequentially from one another or instead represent distinct reprogramming outcomes. To better understand the transcriptional progression during reprogramming, we applied RNA velocity pseudotime, which leverages the relative abundance of spliced and unspliced transcripts to estimate directional gene expression changes and position cells along a transcriptional continuum^35,36^. We ordered cells according to their predicted direction of transcriptional change along a reprogramming trajectory rooted in the initial fibroblast population. To reduce the influence of cell cycle phase on this ordering, we computed velocity pseudotime using root cells selected from both G1/Sand G2/M-phase fibroblasts and averaged the resulting values to generate the final velocity pseudotime trajectory (Figures 3B-C; Figure S13 and Figures S14B, D; Table S4). Control fibroblasts were positioned at early pseudotime and reprogrammed cells at progressively later states (Figures 3C). Diffusion pseudotime analysis^37^ produced a similar ordering (Spearman’s *ρ* = 0.90, Figures S14), corroborating the inferred trajectory.

**Figure 3:**
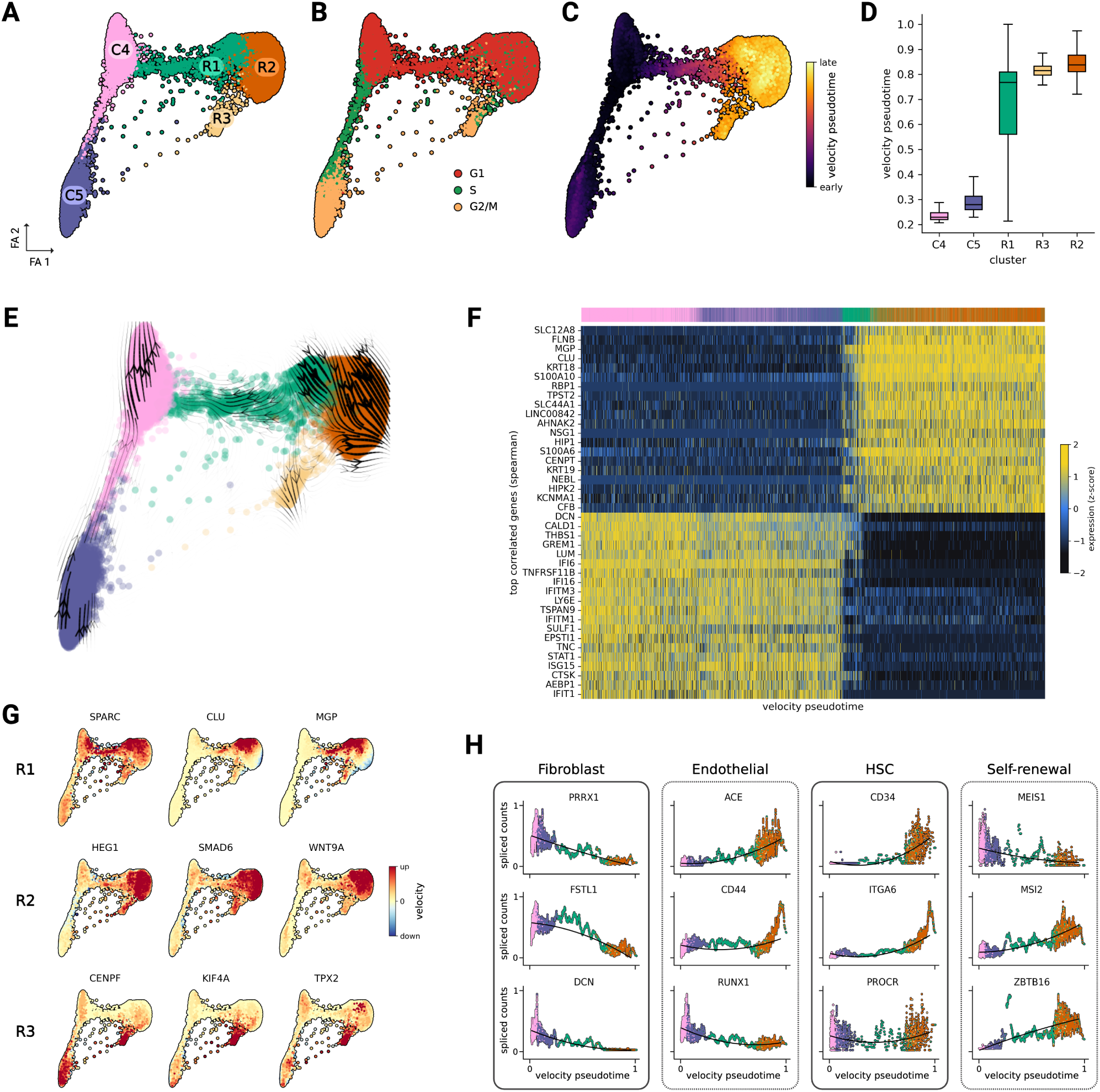
Pseudotime progression and gene expression dynamics from RNA velocity. **(A)** Force-directed graph layout colored by Leiden cluster assignment. **(B)** Same layout colored by cell cycle phase (G1, S, G2/M) used to select root cells for pseudotemporal ordering. **(C)** Cells colored by velocity pseudotime. **(D)** Per-cluster distributions of pseudotemporal ordering (Data are represented as median and interquartile range (IQR), with whiskers extending to 1.5 × IQR). **(E)** Low-dimensional projection of RNA velocity estimates. **(F)** Z-scored expression of the top positively and top negatively pseudotime-correlated genes (Spearman’s *ρ* with Benjamini-Hochberg FDR correction; adjusted *p <* 0.05). Cells (columns) are ordered by velocity pseudotime and annotated by cluster (color bar, top). **(G)** Top velocity genes for reprogrammed cell clusters. **(H)** Spliced (steady-state) gene expression shown over velocity pseudotime for select fibroblast, endothelial, and HSC genes colored by cluster. See also Figures S13-S20 and Table S4.

Cluster R2 was positioned farthest along pseudotime, with R3 closely overlapping, whereas R1 spanned an intermediate range between fibroblast clusters and the later R2/R3 states (Figures 3C-D). Projection of RNA velocity vectors onto the force-directed embedding showed that velocity fields were most coherent within the reprogramming clusters, as supported by velocity confidence scores (Figure S16C). Vectors suggested that subsets of R1 cells transition toward R2 and then continue away from the initial fibroblast state toward a shared future transcriptional state. Subsets of R3 cells were also oriented in this direction, although most R3 vectors followed a distinct trajectory, likely reflecting cell cycle-associated dynamics given the cycling nature of this cluster. We did not detect strong velocity flow from control fibroblasts directly into reprogrammed cells, suggesting that reprogrammed cells have diverged from the initial fibroblast state and occupy a distinct transcriptional landscape, despite retaining residual fibroblast-like features. We note that RNA velocity estimates local transcriptional directionality rather than definitive lineage relationships. We thus interpret the projected vector field as contextual support for the inferred trajectory rather than stand alone evidence of cell state progression.

To examine transcriptional programs associated with reprogramming, we correlated gene expression with velocity pseudotime (Spearman’s *ρ*, adjusted *p <* 0.05), enabling the identification of genes that are induced or lost as cells progress further away from the initial fibroblast state (Figure 3F). Early pseudotime genes included fibroblast markers (*DCN*, *LUM*, *TNC*) and interferon-stimulated genes (*STAT1*, *IFI6*, *ISG15*), consistent with diffusion pseudotime (Figure S14G) and findings from differential expression analysis. Late pseudotime genes included matrix remodeling factors (*MGP*, *CLU*, *FLNB*), cytoskeletal regulators (*KRT18*, *NEBL*, *S100A10*), and transport-related genes (*SLC12A8*, *RBP1*, *KCNMA1*). Positively correlated TFs (enriched in late pseudotime) included early hematopoietic regulators (*ZBTB16*, *ETS2*, *REL*) and developmental factors (*FOXN3*, *SOX13*, *SMAD6*), while negatively correlated TFs were enriched for innate immune responses (*IRF1*, *STAT2*, *SP100*) and fibroblast identity (*PRRX1*, *MKX*) (Figure S15), suggesting coordinated suppression of the fibroblast program alongside induction of early progenitor-like transcriptional regulators.

Cluster-specific velocity genes revealed distinct transcriptional drivers across reprogrammed populations (Figure 3G). R1 was characterized by expression of matrix remodeling genes (*SPARC*, *CLU*, *MGP*), consistent with a transitional identity. R3 exhibited strong velocity signals for cell cycle genes (*CENPF*, *KIF4A*, *TPX2*), representing a transcriptionally distinct, proliferative state. R2 was enriched for endothelial (*HEG1*) and hematopoietic specification genes (*SMAD6*, *WNT9A*). Gene expression dynamics along the pseudotemporal ordering further supported the acquisition of endothelial and hematopoietic features at late pseudotime (Figure 3H). Fibroblast markers decreased over pseudotime while endothelial markers (*ACE*, *CD44*) and HSC markers (*CD34*, *ITGA6*) increased, peaking in R2. Genes involved in HSC maintenance and self-renewal (*MSI2*, *ZBTB16*) also increased over pseudotime. Notably, canonical endothelial and hematopoietic regulators (*RUNX1*, *GATA2*, *SMAD3*, *PECAM1*) and long-term HSC genes (*PROCR*, *MEIS1*, *IL3RA*, *EPAS1*) showed minimal change in steady-state expression but displayed positive velocity signals at late pseudotime, suggesting transcriptional priming of the hematopoietic program (Figures S16D-E).

To better define intermediate stages prior to day 48, we collected a scRNA-seq time-series dataset at days 0, 7, and 21 (D0, D7, D21) of reprogramming (Figure S17). Given that time-series cells were not enriched for a hematopoietic marker, we sought to focus the analysis on cells that showed the most evidence of reprogramming. Clustering within the time-series dataset revealed distinct subpopulations at D7 and D21 (Figure S19D, H). Some clusters retained strong fibroblast-associated gene expression, while others showed substantial loss of the fibroblast program, consistent with the heterogeneity observed among day 48 cells (Figure S19F, J). We then integrated the time-series data with Libraries 1 and 2 and applied scANVI to classify cells as fibroblast-like or day 48-like (Figure S18)^38^. Cells classified as fibroblast-like predominantly belonged to a single cluster within each time point, corresponding to the clusters with the strongest retention of fibroblast gene expression. Clusters with the highest proportion of cells classified as day 48-like included D7-2 and D21-1. Consistent with this classification, cluster D7-2 was enriched for genes associated with transport, metabolism, and cytoskeletal regulation (e.g., *SLC7A11*, *ENO1*, *LPAR1*), resembling features upregulated in day 48 reprogrammed cells (Figure S19G) Cluster D21-1 showed induction of developmental regulators, including *TOX* and *MEIS2* (Figure S19K) We next asked whether this early heterogeneity was concordant with the inferred velocity pseudotime trajectory. To position time-series cells along this axis, we assigned pseudotime values to D0, D7, and D21 cells using a *k*-nearest neighbor projection approach (Figure S20). Projected pseudotime increased across experimental time points from D0 to day 48, supporting progressive transcriptional divergence away from the fibroblast state. However, D7 clusters showed broad pseudotime distributions, with the majority of cells overlapping with fibroblast controls and subsets of cells in D7-2 and D7-3 that overlapped with day 48 cells. Compared to D7, clusters D21-1 and D21-3 showed narrower distributions that were shifted closer to the day 48 states. The overlap of assigned pseudotime values across experimental days suggests that reprogramming does not proceed as a synchronized sequence but instead involved early emergence of day 48-like transcriptional states, followed by an increased frequency of such cells over time.

These findings demonstrate that 5TF reprogramming is associated with coordinated suppression of the fibroblast identity and progressive induction of early hematopoietic and endothelial transcriptional programs. Identification of intermediate states in time-series data suggests rapid induction of reprogramming in subsets of cells.

### Benchmarking against native and engineered cell types

Our analyses thus far demonstrate that reprogrammed cells diverged from fibroblasts and partially acquired hematopoietic features, but do not evaluate the degree to which reprogrammed cells resemble true HSCs, progenitors, or other cell types. To contextualize the reprogrammed cells, we leveraged our reference atlas (fibroblasts, mesenchymal cells, endothelial cells, HSCs, hematopoietic progenitors) and obtained publicly available data from engineered hematopoietic cell types^1,10^

To aid integration across the Oxford Nanopore and Illumina platforms, we generated a third longread scRNA-seq dataset (Library 3) of CD34^+^ hematopoietic stem and progenitor cells (HSPCs) from human bone marrow (Figure 4A). Library 3 cell types were annotated based on expression of canonical hematopoietic marker genes, identifying HSCs and lineage-restricted progenitors (Figure S21), consistent with published data^39^. We first integrated Libraries 1, 2, and 3 using scVI to learn a harmonized latent space while controlling for technical bias and cell cycle effects (Figure 4B)^30^. A uniform manifold approximation and projection (UMAP) embedding of this latent space showed graded expression of *CD34* and other hematopoietic genes across libraries (Figure S22A)^40^, with the highest expression in bone marrow cells. Differential expression analysis revealed substantial downregulation of fibroblast programs in R1-R3, although some extracellular matrix and mesenchymal gene expression persisted in subsets of reprogrammed cells (Figure 4C, Figure S22B). Several hematopoietic genes were upregulated in R1-R3 but remained lower than in native HSCs, supporting incomplete acquisition of the target state (Figure 4C, Figure S22A).

**Figure 4:**
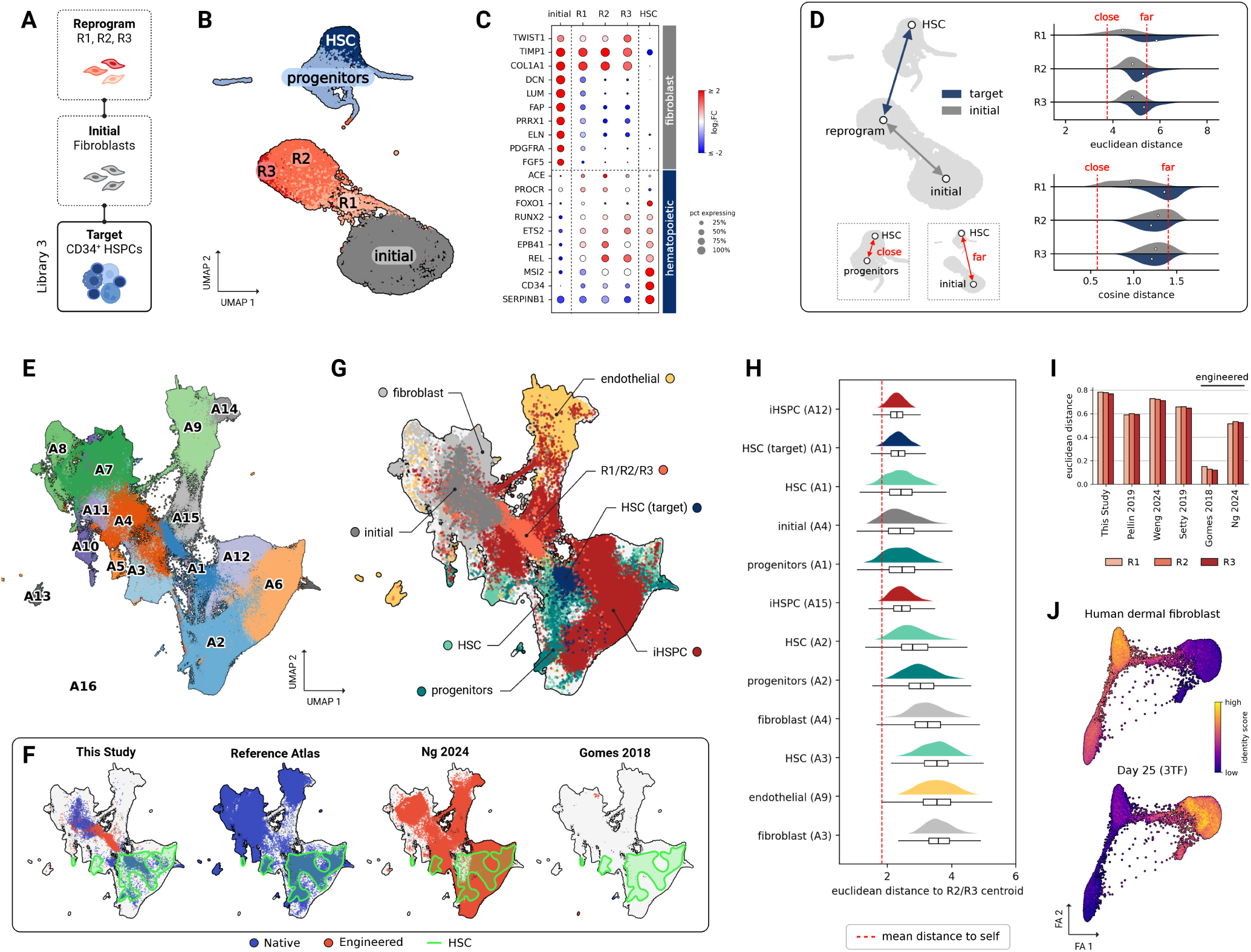
Benchmarking 5TF-induced direct reprogramming. **(A)** Long-read scRNA-seq datasets used for benchmarking, including data from human CD34^+^ bone marrow cells (Library 3: 5, 608 cells). **(B)** scVI-corrected UMAP embedding of cells from (A). **(C)** Select marker genes. Dot color represents log_2_ fold change, and dot size reflects the percentage of cells expressing a gene in a given group. **(D)** Left: schematic of cell–cell distance comparisons; Right: distributions of mean pairwise Euclidean and cosine distances from R1-R3 to the target and initial cell types. White dots indicate the mean distance. Red dashed lines represent the mean distance between close (HSC to progenitors) and far (HSC to initial) cells. **(E)** UMAP of 420,108 cells from the reference atlas and selected sources based on scVI latent space, colored by Leiden cluster. **(F)** Source of cells in (E), colored by native or engineered cell types. The green outline represents HSCs from the native reference atlas. **(G)** Cell groupings of interest from (E). **(H)** Distributions of per-cell Euclidean distances to the centroid of R2/R3 per cluster-resolved cell annotation, sorted by mean distance (Box plots: data are represented as median and IQR, with whiskers extending to 1.5 × IQR). Red dashed line represents the mean distance of R2/R3 cells to the R2/R3 centroid. **(I)** Normalized euclidean distances from pseudobulked profiles of R1-R3 to native and engineered HSCs from a variety of hematopoietic scRNA-seq datasets. **(J)** Capybara^22^ identity scores for human dermal fibroblasts and Day 25 3TF-reprogrammed cells from Gomes *et al.*^10^ projected onto the force-directed graph layout from Fig. 2F. See also Figures S21-S26, and Tables S5 and S6.

To quantify transcriptional similarity, we computed per-cell Euclidean and cosine distances in the scVI latent space between R1-R3 and either initial fibroblasts or target HSCs (Figure 4D). For each reprogrammed cell, distances were calculated to all cells in the reference population and averaged. We defined reference thresholds for “close” (mean HSC-to-progenitor distance) and “far” (mean HSC-to-fibroblast distance) relationships. R1 showed the greatest similarity to initial fibroblasts across both metrics, consistent with limited reprogramming progress. R2 and R3 showed reduced similarity to fibroblasts and increased alignment with HSCs. Notably, Euclidean distance—reflecting absolute separation in the latent space—suggested that R2/R3 remained closer to the initial state, whereas cosine distance—which captures similarity in the direction of latent representations—indicated greater similarity to HSCs. This suggests that R2/R3 have acquired hematopoietic-like transcriptional programs while retaining some aspects of their original fibroblast identity.

We next integrated our three libraries with the external datasets (Figures 4E-G, Figure S23). Leiden clustering of scVI-harmonized data revealed 16 clusters (A1-A16) with substantial overlap between similar cell types across data sources (Figure 4E, Table S5). R2/R3 localized near engineered HSPCs (iHSPCs) while remaining partially proximal to both initial fibroblasts and subsets of native HSCs and hematopoietic progenitors (Figures 4F-G, Figure S24). We computed Euclidean distances between the R2/R3 centroid and individual cells within cluster-resolved cell types in the integrated atlas (Figures 4H-I) R2/R3 were most similar to subsets of iHSPCs and native HSCs, but were also close to initial fibroblasts and native hematopoietic progenitors. A similar analysis of six independent human hematopoietic scRNAseq datasets showed that reprogrammed cells were consistently closer to engineered hematopoietic and stromal populations than to native HSCs (Figure 4I, Figure S25, Table S6) and aligned most closely with day 25 3TF-reprogrammed cells^10^. Application of Capybara^22^, a tool for aiding classification of hybrid cell identities, further supported a high degree of similarity between R1-R3 and engineered cell types (Figure 4J, Figure S26).

Together, benchmarking analyses place reprogrammed cells in an intermediate state between initial cells, endothelial cells, and HSCs.

### Transcript-level diversity in direct reprogramming

While our reference atlas enabled a quantitative comparison of reprogrammed cells to a variety of native and engineered cell types, these analyses relied primarily on gene-level expression, which incompletely reflects molecular and phenotypic changes^41,42^. Commonly used functional assays for evaluating reprogrammed HSCs, such as transplantation or colony-forming assays, lack the resolution needed to quantitatively define intermediate cell states or inform iterative optimization of reprogramming protocols^43^ Protein-level measurements can capture key aspects of cell identity but remain limited in throughput and scalability^44^. Thus, existing approaches provide only a partial view of the molecular changes accompanying reprogramming. Alternatively spliced (AS) transcript isoforms represent an additional regulatory layer that is more directly linked to protein structure, regulation, and post-transcriptional control, and are incompletely captured by short-read sequencing^45,46^. Leveraging our long-read scRNA-seq data, we sought to examine transcript-level changes across initial, reprogrammed, and target cells, hypothesizing that isoform-level analyses could reveal features of cell state obscured at the gene level.

We first characterized global patterns of AS across Libraries 1 − 3. As expected for mammalian cells, exon skipping was the most common AS event observed in each condition (Figure 5A)^47^. To assess transcript-level diversity, we quantified the number of detected transcripts per gene and classified genes as not expressed (NE), expressed as a single isoform, or expressed as multiple isoforms. Between conditions, genes were scored as showing a gain in isoform diversity (NE to single or multiple, single to multiple) or a loss in diversity (multiple to single or NE, or single to NE). A visual representation of this analysis is shown in Figure 5B. Among reprogrammed populations, cluster R2 showed the largest gain in isoform diversity relative to initial fibroblasts, and the greatest overlap with hematopoietic progenitors and HSCs. In contrast, R3 exhibited the greatest loss of isoform diversity and the least overlap with both initial and target states, consistent with more divergent or heterogeneous transcript usage.

**Figure 5:**
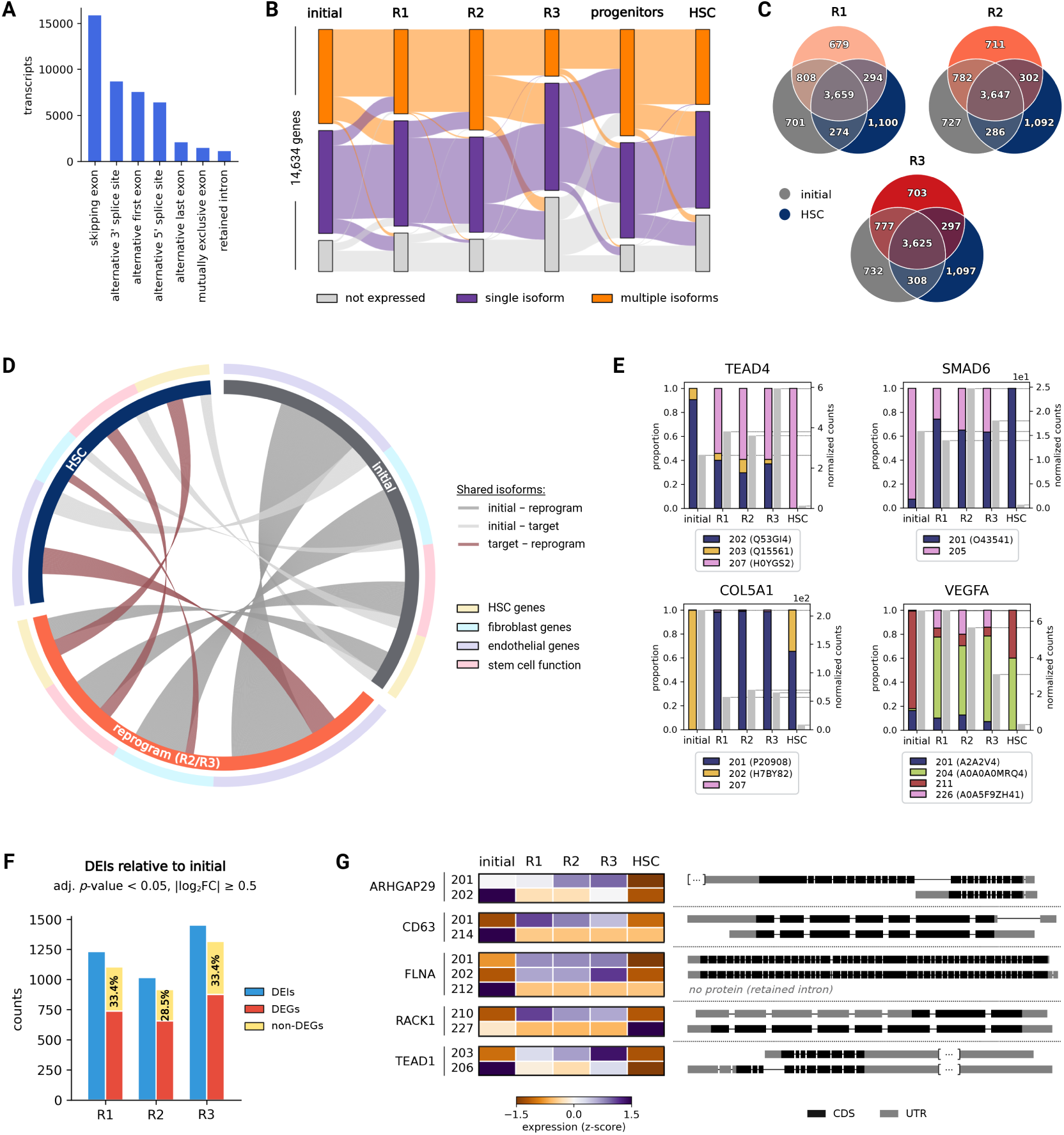
Isoform diversity in fibroblast to HSC reprogramming. **(A)** Count of pooled transcripts by splicing event. **(B)** Changes in isoform usage for protein-coding genes across reprogramming stages. Transcripts with less than 10 counts per group were excluded. **(C)** Overlap of shared major isoforms between reprogrammed clusters and initial or target cell types. **(D)** Shared major isoforms across cell groupings for curated gene sets from GO Biological Process^49,50^ and marker gene databases^28,32,51^. Links between transcripts signify shared major isoforms between cell groups. Unlinked transcripts represent unique major isoforms. **(E)** Isoform switching events. Bar plots show the proportion of total gene expression for each detected transcript, with gray bars signifying total normalized counts per gene (counts per million, CPM). Transcripts are labeled with corresponding UniProt accessions. **(F)** Left bars: Number of DEIs for each reprogrammed cluster relative to initial cells. Right bars: Proportion of corresponding genes that are or are not differentially expressed at the gene-level. **(G)** Left: Z-scored expression (CPM) for representative transcripts where the gene is not differentially expressed at the gene-level. Transcripts were filtered for those accounting for > 10% of total counts per gene. Right: Exon structures for corresponding transcripts with coding sequences (CDS) and untranslated regions (UTR) annotated according to the GRCh38 reference genome (10x Genomics, 2024-A).

To further assess transcript-level differences, we identified protein-coding genes whose expression was dominated by a single transcript, designating this transcript as the ‘major isoform’. HSCs exhibited the largest number of unique major isoforms, consistent with a distinct transcriptomic architecture (Figure 5C). Among reprogrammed clusters, R2 shared the greatest number of major isoforms with HSCs, whereas R3 showed the least overlap with the initial fibroblast state. R2/R3 displayed the highest numbers of unique major isoforms among reprogrammed populations, reflecting progressive transcript-level remodeling associated with later reprogramming states. Similar patterns were observed when examining major isoform overlap across curated gene sets, including fibroblast-, endothelial-, HSC-, and stem cell-associated genes (Figure 5D). For all gene sets, reprogrammed cells shared the greatest proportion of major isoforms with initial fibroblasts, while exhibiting increased overlap with HSCs relative to initial cells. Across gene sets, all cell populations retained moderate to high fractions of unique major isoforms, highlighting the prevalence of transcript-level differences across initial, reprogrammed, and target cell states.

We next identified genes undergoing isoform switches (IS) across cell types (Figure 5E), defined as changes in the dominant protein-coding transcript. Several genes exhibited consistent IS between reprogrammed cells and the initial fibroblast state, with R1-R3 preferentially expressing transcript isoforms that encode distinct protein products. For example, initial fibroblasts predominantly expressed the Ensembl canonical isoform *TEAD4-202* (UniProt: Q53GI4), whereas R1-R3 primarily expressed *TEAD4-207* (UniProt: H0YGS2), which lacks annotated binding domains present in the canonical transcript. This example illustrates how IS can introduce qualitative changes in predicted protein structure that are not captured by gene-level expression analyses.

To quantify transcript-level regulation independent of gene-level differential expression, we identified differentially expressed isoforms (DEIs) in clusters R1-R3 relative to initial fibroblasts and assessed whether the corresponding genes were differentially expressed at the gene level (Figure 5F). Across all reprogrammed clusters, approximately 30% of DEIs occurred in genes without detectable gene-level differential expression, suggesting that a substantial fraction of transcriptional remodeling would be missed by gene-centric analyses. Visualization of representative DEIs from genes lacking gene-level differential expression revealed pronounced changes in transcript usage across conditions (Figure 5G). Notably, several DEIs exhibited changes in both coding sequences and untranslated regions, which regulate mRNA stability, localization, and translation efficiency^48^.

Together, these findings demonstrate that isoform-level analyses reveal substantial transcriptional remodeling during reprogramming that is not apparent from gene-level expression alone, and underscore the value of long-read transcriptomics for resolving molecular features of reprogramming cell states.

## DISCUSSION

Our study highlights the challenge of developing new direct reprogramming protocols and evaluating the resulting cells. The outcome of partial reprogramming for a novel protocol contributes to inefficient discovery, since partially reprogrammed cells are not likely to pass classical functional testing despite having some desired properties. Thus new frameworks are needed to evaluate whether or not outcomes are worthy of further development. Here we integrate prediction of new recipes with an evaluation workflow; our prior DGC algorithm^26^ predicted a TF recipe for hematopoietic reprogramming while a reference atlas generated using 270 publicly available datasets aided the evaluation of reprogrammed cells. Our analysis showed acquisition of an early hematopoietic- or endothelial-like program and previously uncharacterized patterns in gene and isoform expression. Reprogrammed cells showed transcriptomic shifts that distinguished them from fibroblasts, but they also remained distinct from native HSCs.

Among the predicted TFs, GATA2, GFI1B, and FOS are key hematopoietic regulators that have been used in previous studies for direct reprogramming to HSCs^10,11,52^, supporting that the algorithm-predicted TFs are a practical starting point for recipe discovery. Of the novel predicted factors, STAT5A is a hematopoietic regulator that supports creation of HSCs from murine embryonic stem (ES) cells and has a role in HSC self-renewal, proliferation, and survival^53–55^. REL encodes a subunit of the NF*κ*B complex, which has an important role in the emergence of HSCs during embryonic development^56^ Proinflammatory signaling via NF-*κ*B is involved in HSC specification alongside Notch signaling in the hemogenic endothelium^57,58^. REL also regulates apoptosis by promoting expression of anti-apoptotic genes *BCL* and *BCL2L1*^59–61^. Together, STAT5A and REL are shown to regulate chromatin accessibility at shared shared target loci^62^, supporting a model in which they act as complementary regulators in the hematopoietic gene expression program. However, the context-dependent role of these TFs in reprogramming requires further elucidation.

Prior studies support a model in which fibroblast-to-HSC reprogramming proceeds through an endothelial precursor, exhibiting core aspects of developmental endothelial-to-hematopoietic transition (EHT)^12,63^ For example, murine fibroblast-to-HSC reprogramming studies report endothelial-like and hemogenic endothelial intermediates during early reprogramming stages, including expression of embryonic hemoglobin (*b-H1*), *Pecam1*, and *Vwf* ^64,65^. Human fibroblast-to-HSC studies likewise show expression of endothelial genes such as *PPARG*, *VWF*, and *FOXC2*^10,11,66^. Similarly, our reprogrammed cells expressed endothelial genes such as *ACE*, *PECAM1*, and *PODXL*. Consistent with this endothelial intermediate model, gene expression dynamics along the 5TF reprogramming trajectory suggested transcriptional priming of key endothelial and hematopoietic regulators, including *RUNX1* and *GATA2*. Overall, for the 5TF recipe, our data support a model in which some cells rapidly acquire a reprogrammed phenotype by day 7 after induction, progress to a more stable and well-defined population by day 21, and show a lineage relationship among clusters at day 48. The dynamics and drivers of these transitions remain unknown.

Notably, within our integrated reference atlas, 5TF cells showed substantial overlap with subsets of iPSC-derived iHSPCs from Ng *et al.*, a system in which multilineage bone marrow engraftment has been demonstrated in immunodeficient mice^1^. The transcriptional proximity of our cells to an engineered hematopoietic system with demonstrated in vivo potential supports that 5TF cells are progressing along a trajectory compatible with HSC specification. While the Ng *et al.* cells generate a broad spectrum of hematopoietic cell types, our 5TF cells appear to follow a more constrained, direct trajectory from fibroblasts toward HSCs, forming a dense population positioned between endothelial cells, HSCs, and fibroblasts. Given that iPSC-based approaches recapitulate aspects of developmental EHT, our findings raise the possibility that direct reprogramming may similarly engage key features of the developmental program but through a more defined or accelerated route. Future studies will be required to determine whether direct fibroblast-to-HSC reprogramming follows a path similar to iPSC-based hematopoiesis, and whether this potentially accelerated path can likewise produce functionally equivalent HSCs.

finding evidence of isoform switching in genes critical to hippo signaling if these are functional, would imply altered hippo signaling (this has been noted before in cell reprogramming)

find link to some4thing making it related to HSCs

Long-read single-cell profiling revealed isoform-level changes during direct reprogramming that are missed at the gene level. Across our reprogrammed cells, hundreds of transcripts showed differential isoform usage without corresponding changes in total gene expression. These isoform-level differences may have important functional consequences: distinct transcripts from the same gene can encode proteins with different domain architectures or contain alternative untranslated regions that influence post-transcriptional regulation^47,48^. Such mechanisms are particularly relevant in reprogramming, where subtle changes in TF activity or signaling pathways can alter how cells interpret cues and respond to fate-determining signals. Indeed, prior work has shown that reprogramming to pluripotency involves distinct, sequential phases of splicing regulation^67–70^ and that isoform switching in TFs can influence reprogramming outcomes^71,72^. In this study, we identified isoform switching in several genes associated with the Hippo signaling pathway, such as the TEAD-family TFs *TEAD1* and *TEAD4*. Hippo signaling has established roles in hematopoietic development^73,74^ and has been implicated in reprogramming^75^. If these transcript isoforms produce functionally distinct protein products, they could alter Hippo pathway activity without changes in overall gene expression, providing a potential mechanism through which alternative splicing contributes to cell fate transitions. Importantly, the functional impact of individual isoforms cannot be inferred from transcriptomic data alone, and targeted experimental validation will be required to determine how specific isoforms contribute to reprogramming efficiency or the stability of a specific cell state.

Despite acquisition of CD34^+^ surface expression and marked transcriptional changes, the reprogrammed cells retained expression of key fibroblast markers such as *COL1A1* and *FTL*^32^. Complete reprogramming may require overcoming barriers in cell state transitions, including both local stability and maintenance of the new identity^76–78^. Quantifying intermediate or hybrid cell states remains a key challenge^22,79^. Computational tools such as Capybara^22^ enable partial assignment of transcriptomic profiles to single cells, supporting estimation of hybrid identities; however, the accuracy of this approach is limited by the availability of suitable reference data. Our approach parallels other recent atlasing strategies for identifying rare, diseased, or engineered cell states and evaluating new engineering protocols^24,80,81^. Future studies may assess the applicability of emerging single-cell foundation models^23,25,82,83^, though their capacity to represent engineered and intermediate cell types remains to be determined.

Contextualization of reprogrammed cell states provides critical feedback for refining reprogramming protocols. Systematic characterization of similarities and differences between reprogrammed and native cell types can reveal barriers to efficient conversion and identify opportunities to overcome them. Beyond the present system, our approach is broadly applicable to other contexts where engineered or perturbed cells occupy intermediate states, including regenerative medicine, disease modeling, and synthetic biology In summary, our approach offers a generalizable framework for evaluating unknown cell identities and contributes to the effort to improve direct reprogramming methods for generating autologous cells.

### Limitations of the study

While the time points evaluated by scRNA-seq contained a minimum of 5,000 cells each, there were a limited number of time points for defining reprogramming progression, and each sample was a single replicate. In addition, the time-series data showed a lower depth of sequencing than the other datasets, which in particular could hinder the capture and analysis of transcripts expressed at low levels. The methods used do not resolve whether cell clusters at a given time point retain a lineage relationship or whether they form independent trajectories. In addition, our study does not address potentially novel transcriptional states in intermediate cells that differ substantially from known cell states^22,29,79^. While the data highlight genes that undergo isoform switching, further study is required to validate these differences at the protein level. The current reprogramming population, given its partially-reprogrammed phenotype, was not classically tested for HSC features, for example via colony-forming unit assay. This study was not powered to consider sex-based differences in the chosen cell types. Male BJ fibroblasts were chosen to maintain experiment continuity and alignment with past datasets. The active lot for ATCC human CD34^+^ bone marrow cells was from a male donor, which was not expected to impact outcomes in its use as a biological positive control for hematopoietic markers in flow cytometry. For human CD34^+^ bone marrow from STEMCELL Technologies, the male donor was chosen based on cell quantity, donor health, and body mass index.

## RESOURCE AVAILABILITY

### Lead contact

Requests for further information and resources should be directed to and will be fulfilled by the lead contact, Lindsey A. Muir.

### Materials availability

This study did not generate new unique reagents.

### Data and code availability

- Single-cell RNA sequencing data have been deposited to the Gene Expression Omnibus (GEO) repository (GEO: GSE283658) and is publicly available.
- This paper analyzes existing, publicly available data. All accession numbers for curated scRNA-seq datasets can be found in Table S6. Dataset and standardized ontology identifiers for cells in the reference atlas can be found in Data S1.
- All original code is publicly available at https://github.com/jrcwycy/Hematokytos as of the date of publication.
- Any additional information required to reanalyze the data reported in this paper is available from the lead contact upon request.

## ACKNOWLEDGMENTS

We thank Max Wicha at the University of Michigan for valuable discussions conversations and ideas on how to present results. We thank the University of Michigan Advanced Genomics Core for assistance with 10x Genomics single cell barcoding and the Flow Cytometry Core for assistance with cell sorting We acknowledge BioRender, which was used to assemble figures and for generating the graphical abstract. This work was supported by the National Science Foundation (NSF) award number 2035827 (IR) subcontracted from iReprogram, Inc., the Air Force Office of Scientific Research (AFOSR) award number FA9550-22-1-0215 (IR), the Defense Advanced Research Projects Agency (DARPA) award number HR00112490472 (IR and LM), NIGMS GM150581 (JC), iReprogram, Inc, and ONT. Computational resources and services were provided by Advanced Research Computing at the University of Michigan, Ann Arbor (RRID: SCR_027337). We thank Lakmal Jayasinghe and the research and support teams at ONT for helpful discussions and technical assistance.

## AUTHOR CONTRIBUTIONS

Conceptualization, LM and IR; methodology, LM, IR, JC, WM, CMS, and JP; investigation, WM, HL, LM, JC, CMS, and SG; writing – original draft, JC and CMS; writing – review and editing, JC, CMS, SG, HL, JP, WM, LM, and IR; funding acquisition, LM and IR; resources, LM and IR; software, JC, CMS, and SG; visualization, JC, CMS, SG, and LM; supervision, LM and IR.

## DECLARATION OF GENERATIVE AI AND AI-ASSISTED TECH NOLOGIES

During the preparation of this work, the authors used ChatGPT (OpenAI) and Gemini (Google) for copyediting. Gemini was also used for source code generation and documentation in early versions of the manuscript. The authors reviewed and edited the content as needed and take full responsibility for the content of the published article.

## SUPPLEMENTAL INFORMATION INDEX

Document S1. Figures S1-S22, Tables S1-SX, and supplemental references in a PDF.

Data S1. Metadata for CELLxGENE datasets used to generate the reference atlas, related to Figure 1, in a .csv file.

## STAR★METHODS

### Experimental model and study participants detail

Human fibroblasts

Human neonatal fibroblasts were used as the initial cell type for reprogramming experiments. Cryopreserved BJ fibroblasts were purchased from ATCC (#CRL-2522, RRID: CVCL_3653). BJ cells were derived from a male donor and authenticated for experimental use by the commercial supplier. Cells were thawed and expanded in growth medium (GM): Dulbecco’s Modified Eagle Medium high glucose (DMEM; Gibco, #11965092) with 10% fetal bovine serum (FBS; Corning, #35-015-CV). Cells were cryopreserved at passage 2 or 4 and freshly thawed for each experiment. 1x TrypLE Express Enzyme (Gibco, #12604-021) was used for dissociation and cell passaging. Cells were cultured at 37°C with 5% CO_2_.

Human CD34^+^ HSPCs

Primary CD34^+^ hematopoietic stem and progenitor cells (HSPCs) from adult male human bone marrow were purchased from STEMCELL Technologies (#70002, Lot 2410421008, Donor ID: CE0010639), containing 1 × 10^6^ cells. CD34^+^ HSPCs were thawed according to the 10x Genomics cell thawing protocol (#CG000447, Rev B, February 2024). The primary cell donor was chosen based on quantity of cells, donor health, and body mass index, and sex was not considered in this selection. This study was not powered to assess sex-based differences.

As additional staining controls in flow cytometry experiments, male human primary bone marrow CD34^+^ cells were purchased from ATCC (#PCS-800-012, Lot 80516705) and cultured in complete MyeloCult H5100 (STEMCELL Technologies, #05150) for 12-16 hours prior to analysis. All cells were cultured at 37°C with 5% CO_2_. Lot number was determined by the commercial source, and choice of sex was not available.

### Method details

#### Reference atlas construction

Single cell transcriptomic data were retrieved from the CELLxGENE Census (2024-07-01) using the cellxgene-census Python API. We selected 1,125,041 primary cells (52,164 genes) from non-diseased human samples representing major hematopoietic, myeloid, lymphoid, innate lymphoid, fibroblast, mesenchymal, and endothelial cell types. Standardized source data metadata, such as cell type and tissue annotations, were stored along with gene expression data for downstream processing and analysis using Scanpy^84^ (version 1.11.4) and/or rapids-singlecell^85^ (version 0.12.3). Consistent quality control was applied across all cells. First, data were filtered for cells with at least 200 UMI reads and genes with a minimum of 50 reads. Expression was normalized to 10,000 counts per cell, followed by log-transformation using log1p.

We performed batch harmonization and biologically meaningful latent space extraction in two steps. First, we used scVI^30^ implemented in the scvi-tools (version 1.3.3) Python library^86^. We developed a scVI model with two hidden layers and a 24-dimensional latent space, a negative binomial likelihood based on individual dataset set identifiers and cell type annotations. Training was performed on GPU (NVIDIA Tesla V100 or A100) for up to 100 epochs with early stopping (patience = 5), using a batch size of 1000 and monitoring validation loss. Next, we initialized a semi-supervised scANVI from the pretrained scVI model to improve cell type influence on the learned latent space^38^. The scANVI model was trained for up to 20 epochs using GPU acceleration, with early stopping (patience = 5) based on validation loss. Training used a batch size of 1,000 and 100 samples per label. Training was performed on the same GPU hardware while monitoring validation loss. For visualization, we developed a t-SNE embedding based on the scANVI model latent space. To identify DEGs, we used scanpy.tl.rank_genes_groups using the Wilcoxon rank-sum test (two-sided) over log-normalized expression. DEGs were identified for major cell compartments and cell types, excluding cell types with fewer than 500 cells.

#### TF recipe selection

TFs for fibroblast-to-HSC reprogramming were selected using the previously constructed “data-guided control” (DGC) algorithm^26^. The DGC algorithm provided a ranked list of TFs that partially overlapped with a prior published TF recipe (GATA2, GFI1B, FOS), and STAT5A and REL were selected since they were among the top five TFs by rank. Briefly, in this framework, TFs are identified by minimizing the distance between the initial and target cell states according to the following linear difference equation:

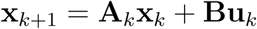

where:

- **x***_k_ ∈* R*^N^* : gene expression vector at time *k*
- **A***_k_ ∈* R*^N×N^* : transition matrix representing time-varying changes in gene expression
- **B** ∈ R*^N×M^* : binary input matrix capturing interactions between genes (*N*) and TFs (*M*)
- **u***_k_ ∈* R*^M^* : input vector indicating which TF(s) are introduced at time *k*

TFs are ranked through the following optimization:

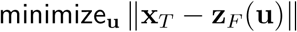

The objective is to minimize the Euclidean norm, ‖ · ‖, of the difference between the target state **x***_T_*and the final state **z***_F_*(**u**), where **z***_F_*depends on the initial condition **x_0_** and the input signal **u**.

#### Lentiviral vector design

Human GATA2, GFI1B, FOS, STAT5A, and REL plasmids were procured from the DNASU repository, subcloned into pLIX-403 Tet ON lentiviral backbones (RRID:Addgene_41395), and verified by Sanger sequencing. For TF combinations labeled FH3 (GATA2, GFI1B, FOS) and FH6 (GATA2, GFI1B, FOS, STAT5A, REL), lentiviral vectors were prepared individually for each plasmid, then pooled for transductions (Table S7). For recipe FH4a (GATA2, GFI1B, FOS, STAT5A, REL), plasmids were pooled in equimolar amounts followed by lentiviral packaging. All lentiviral vectors were prepared by the University of Michigan BRCF Vector Core (RRID:SCR_026696). AllPrep DNA/RNA/Protein Mini Kits (Qiagen, #80004) were used to process cells for testing genome integration and protein and gene expression.

#### Lentiviral transduction

BJ fibroblasts were plated at a density of 21, 000 − 26, 000 cells/cm^2^ in either six-well plates or 10 cm dishes. The next day, cells were transduced with one of three TF recipes in Table S7. Transductions were performed in standard GM containing 8 *µ*g/mL polybrene (Millipore Sigma, #TR-1003-G). GM supplemented with 1% penicillin/streptomycin (P/S; Gibco, #15140-122) was exchanged 24 hours posttransduction. Recipes FH3 and FH6 were used for flow cytometry experiments, while FH4a was used for transductions in all other experiments.

#### Reprogramming induction

48 hours after transduction, TF expression was induced via addition of 1 *µ*g/mL doxycycline hyclate (Dox; Millipore Sigma, #D9891). We consider this Day 0 of reprogramming. On Day 2, cells were plated in six-well plates coated with 0.1% porcine gelatin (Millipore, #SF008) in complete MyeloCult H5100 (STEMCELL Technologies, #05150) + 10e-6 M hydrocortisone (Millipore Sigma, #H0888) + 1% P/S. Media was exchanged every other day for the duration of experiments. Reprogramming cultures were grown for 7 − 48 days, with cell passaging occurring every 6 days. Cells used for control and reprogramming cultures were used at initial passage number between 2 and 4, and were at 10 or fewer passages for endpoints. All cells were cultured at 37*^◦^*C with 5% CO_2_.

#### qRT-PCR validation

Total RNA was collected from independent replicates (*n* = 3) of fibroblasts transduced with FH4a (Table S7) using the RNeasy Kit (Qiagen, #74104) after 3 days of 1 *µ*g/mL Dox treament in complete Myelocult media. The yield and purity of RNA samples were assessed by Nanodrop and Qubit (Thermo Fisher Scientific). Using 1 *µ*g of purified RNA per sample, first-strand cDNA synthesis was performed using SuperScript IV Reverse Transcriptase (Invitrogen, #18090010) with random hexamers (Invitrogen, #N8080127). Quantitative real-time PCR (qRT-PCR) was conducted using SYBR Green qPCR Master Mix (Thermo Fisher Scientific, #A66732) for all samples in triplicate in a QuantStudio 5 Real-Time PCR System (Applied Biosystems). Relative quantification was determined using the ΔΔCt method and normalized to the endogenous control LDLR. Primer efficiency was verified for each exogenous TF prior to running qRT-PCR. See Table S8 for a list of primers. Quantile-quantile plots were used to assess data normality, and group differences were evaluated using an unpaired *t*-test with Welch’s correction (two-sided). Statistical details are provided in Table S3.

#### Flow cytometry

Antibodies used for flow and/or cell sorting included anti-human CD90 PE (1:250, BioLegend # 328109, RRID: AB_893442), CD34 Alexa Fluor 488 (1:250, BioLegend # 343517, RRID: AB_1937204) or CD34 PE (1:250, BioLegend # 343605, RRID:AB_1732033), and CD143/ACE (1:250, BioLegend # 344207, RRID: AB_2783230). LIVE/DEAD^®^ Fixable Yellow Dead Cell Stain Kit (0.8 *µ*L per test, Invitrogen, #L34968) and DAPI were used as viability markers. Cells were incubated with primary antibodies or LIVE/DEAD stain for 30 minutes at 4*^◦^*C in Hank’s balanced salt solution or phosphate buffered saline (PBS; Gibco, #10010-023). UltraComp eBeads (Invitrogen, #01-2222) and ArC Amine Reactive Beads (Invitrogen, #A10628) were used for single color compensation controls. After gating for singlets and live cells, fluorescence-minus-one (FMO) controls were used to aid gating for reprogrammed cells. As additional staining controls, human primary bone marrow CD34^+^ cells were purchased from ATCC (#PCS-800-012TM, Lot 80516705) and cultured in complete MyeloCult H5100 (STEMCELL Technologies, #05150) for 12-16 hours prior to analysis, and Veri-Cells CD34 PBMCs (BioLegend, #426901, Lot B375717) were reconstituted following the manufacturer’s protocol. Flow cytometry data were collected on a Bio-Rad ZE5 Cell Analyzer and cells were sorted using a MoFlo Astrios or a Bigfoot Spectral Cell Sorter. FlowJo (Becton Dickinson; version 10.10.0) was used for flow cytometry analyses. For % CD34^+^ comparisons between independent replicates of 5TF-induced cells (*n* = 6) and fibroblast controls (*n* = 2), statistical significance was determined using a one-sided Mann-Whitney U test (mean ± standard deviations for controls: = 0.285 ± 0.233, and 5TF-induced cells: = 2.603 ± 0.835; Mean difference = 2.318; *p* = 0.0357).

#### Immunofluorescence

On day 26 of reprogramming, cells were fixed with 4% paraformaldehyde (PFA; Alfa Aesar, #30525-89-4) for 10 minutes, washed with cold PBS, and permeabilized for 10 minutes in PBS containing either 0.25% Triton X-100, 100 µM digitonin, or 0.5% saponin. After washing with PBS, blocking was performed using 1% bovine serum albumin (BSA; Sigma, #A7906) and 22.52 mg/mL glycine (Sigma, #G8898) in PBS with 0.1% Tween 20 for 30 minutes. Cells were then incubated overnight at 4°C in primary antibody solution containing anti-human CD34 PE (1:200, BioLegend, Clone 581, #343505, Lot B373336) and 1% BSA/PBST. Cells were washed with PBS and counterstained with 2 µM Hoechst 33342 (Cayman Chemical, #15547) for 10 minutes. Imaging was performed using a Zeiss LSM 710 confocal microscope.

#### CD34^+^ HSPC sorting

DAPI was used as a cell viability marker. Veri-Cells CD34 PBMCs (BioLegend) were reconstituted following the manufacturer’s instructions and served as staining controls. Approximately 3 × 10^5^ Veri-Cells were used for full staining and FMO controls. UltraComp eBeads (Invitrogen, #01-2222) were used for single-color compensation. Approximately 1 × 10^6^ bone marrow CD34^+^ HSPCs (STEMCELL Technologies) were prepared for sorting. Cells were first incubated with Fc receptor block (BioLegend, #422301) for 10 minutes on ice to minimize nonspecific antibody binding. Subsequently, cells were incubated with conjugated antibodies for 30 minutes on ice in the dark, washed with 2 mL of cell staining buffer (BioLegend, #422301), pelleted, and resuspended in fresh buffer (BioLegend, #420201). Antibodies included APC anti-human CD34 (Clone 581, BioLegend #343509, RRID: AB_1877154), PE anti-human CD38 (Clone HIT2, BioLegend #303505, RRID: AB_314357), and FITC-conjugated human Lineage Cocktail (Clone UCHT1; HCD14; 3G8; HIB19; 2H7; HCD56, BioLegend #348801, RRID: AB_10612570), which labels T cells, monocytes/macrophages, NK cells, B cells, neutrophils, and eosinophils. Dead cells and FITC-positive cells were excluded during gating. CD34^+^/CD38^−^ and CD34^+^/CD38^+^ cell populations were sorted on a Cytek Aurora-1 instrument.

#### Cell hashing

For hashing of fibroblasts and bone marrow HSPCs, sorted cells were incubated with Fc receptor block (BioLegend, #422301) for 10 minutes on ice, followed by incubation with TotalSeq-B human hashtag antibodies (BioLegend) for 30 minutes at 4°C. Prior to use, antibodies were centrifuged at 14, 000 × g for 10 minutes at 4°C to remove protein aggregates. After labeling, cells were washed three times with 2 mL of cell staining buffer (BioLegend, #422301), with centrifugation at 200 × g for 5 minutes at 4°C after each wash.

#### Single-cell transcriptomics

A summary of the scRNA-seq datasets generated in this study is shown in Table S9. All datasets were generated using 10x Genomics single-cell barcoding followed by Oxford Nanopore Technologies (ONT) long-read sequencing, unless otherwise specified. Cells were partitioned into Gel Beads-in-Emulsion (GEM), where cDNA was synthesized from polyadenylated transcripts and labeled with cell-specific barcodes and unique molecular identifiers (UMIs). Full-length barcoded cDNA was pooled, purified with SILANE magnetic beads (10x Genomics, #PN-2000048), amplified by PCR, and prepared for ONT sequencing according to manufacturer protocols. Briefly, cDNA amplicons were biotin tagged by PCR using the primers shown in Table S10. Biotinylated cDNA was purified with Dynabeads M-280 Streptavidin beads (Invitrogen, #11205D), followed by secondary PCR amplification and Agencourt AMPure XP bead clean-up (Beckman Coulter, #A63881). Products were assessed for quality following ONT recommendations. Libraries were sequenced on ONT GridION or PromethION P2 platforms using the corresponding protocols shown in Table S10 and basecalled using Guppy (version 7.1.4) or Dorado (versions 7.2.13, 7.3.9, 7.6.7) with the High Accuracy models.

#### Reprogrammed cell library

Cryopreserved 5TF-induced cells (FH4a recipe; frozen on day 26) were expanded in complete Myelocult with Dox and harvested on day 48 of reprogramming. 20,000 FACSsorted CD34^+^ reprogrammed cells with 2,000 CD34spike-in cells were processed using the 10x Genomics Next GEM Single Cell 3’ Kit V3.1 (100% viability). Libraries were prepared for sequencing on the ONT GridION or P2 platform as described above.

#### Control fibroblast library

Untransduced BJ fibroblasts (passage 6) were processed using the 10x Genomics Next GEM Single Cell 3’ Kit V4 following the same workflow as Library 1. Prior to single-cell barcoding, cells were stained with 16 *µ*M Hoechst 33342 for 50 minutes and sorted by cell cycle phase (G1, S, G2/M) using a Bigfoot Spectral Cell Sorter. Each population was labeled with a TotalSeq-B hashtag antibody (BioLegend, # 394631, #394633, and #394635, respectively), followed by pooling and single cell barcoding (100% viability). Transcript and hashtag libraries were prepared as described above and sequenced on the ONT P2 platform.

#### Bone marrow HSPC library

CD34^+^/CD38^−^ and CD34^+^/CD38^+^ HSPCs were labeled with TotalSeqB hashtags (BioLegend, # 394631, # 394633, respectively) and pooled prior to 10x Genomics Next GEM Single Cell 3’ V4 barcoding (∼72% viability). Approximately 8,000 cells (6,400 CD34^+^/CD38^−^ and 1,600 CD34^+^/CD38^+^) were processed. Libraries were prepared and sequenced as described above on the ONT P2 platform.

#### Reprogramming time-series

5TF-induced cells were collected on days 0, 7, 14, and 21 of reprogramming, along with non-transduced Doxfibroblast controls. 1 × 10^5^ − 1 × 10^6^ cells were washed with PBS, dissociated using TrypLE Express (Gibco), and pelleted (350 × g, 10 min, 4°C). The Evercode Cell Fixation v3 kit (Parse Biosciences) was used to fix cells in each sample. Prior to fixation, DNase I (Thermo Scientific, #EN0523) and RiboLock RNase Inhibitor (Thermo Scientific, #EO0381) were added to cell suspensions to prevent clumping and remove RNA-degrading enzymes. Fixed cells were processed using the Evercode WT Mini v3 kit (Parse Biosciences). Briefly, up to 5,000 cells per time point underwent split-pool combinatorial barcoding across three rounds (reverse transcription, ligation 1, ligation 2 with UMI incorporation and biotinylation). After barcoding, cells were pooled and lysed. Biotinylated cDNA was isolated using streptavidin-coated magnetic beads, followed by template switching and amplification. cDNA amplicons were then prepared for ONT sequencing using the Ligation Sequencing Kit (SQK-LSK114) and sequenced on the ONT P2 platform.

#### scRNA-seq data processing

Raw sequencing reads from Libraries 1 − 3 were processed using the EPI2ME Labs wf-single-cell pipeline (github.com/epi2me-labs/wf-single-cell). Reads were first aligned to the GRCh38 reference genome (10x Genomics, 2024-A), and gene-level count matrices were generated using either 3’ V3 or 3’ V4 barcode sets. The following parameters were used: full_length=True, expected_cells=10000, barcode_min_quality=15, barcode_max_ed=2, and gene_assigns_minqv=30.

Reads from each library were subjected to quality control using Scanpy. To integrate Library 1 (reprogram) and Library 2 (control), low quality cells were filtered in a library-aware manner using median absolute deviation (MAD) thresholds. Cells exceeding five MADs from the library-specific median for log-transformed total counts or detected genes, or showing high mitochondrial content (> 5 MADs or > 20%), were filtered out. Across both libraries, genes with fewer than 10 total UMI counts were removed, and raw counts were normalized to 10,000 counts per cell and log-transformed using scanpy.pp.log1p

Highly variable genes (HVGs) were selected using the seurat_v3 method with library as the batch key, and principal component analysis (PCA) was performed using scanpy.pp.pca. Data was integrated using Harmony^87^ (version 0.0.10) with library as the batch key. A neighborhood graph was computed using the Harmony-corrected PCA space (n_neighbors=100, n_pcs=25). A force-directed graph was constructed using scanpy.tl.draw_graph with a ForceAtlas2^88^ layout, and leiden clustering was performed with resolution=0.3.

Cells from Library 3 (bone marrow) were processed separately. Cells with fewer than 1,000 detected genes or fewer than 500 total UMI counts were filtered out, and genes with fewer than 10 total UMI counts were removed. Unique UMI counts were normalized to 10,000 counts per cell and log-transformed using log1p. PCA was performed on HVGs, and a neighborhood graph was computed with 45 nearest neighbors. A UMAP embedding was constructed (min_dist=0.2), and Leiden clustering was performed with resolution=0.38. Clusters were annotated based on marker gene expression for expected cell types.

#### TotalSeq-B data processing

We developed a pipeline using Snakemake^89^ (version 7.32.0) to process BioLegend TotalSeq-B libraries sequenced on ONT platforms. Briefly, reads were demultiplexed based on hash sequences using nanoplexer (https://github.com/hanyue36/nanoplexer), with parameters -L 150 -m 3 -x 2 -o 3 -e 1. Following demultiplexing, barcode correction was performed using Flexiplex^90^, with parameters -e 1 -x ′′ -b-u ′′ -x ′′. Corrected barcode-hash sequence pairs were quantified and aggregated per cell. For Library 2, the hash-sequence with the most read-pairs per cell barcode was taken as the barcoded phase for each cell.

#### Parse data processing

To process raw reads from the reprogramming time-series library, the Parse Biosciences split-pipe scRNA-seq pipeline (version 1.7.3) with adaptations for long-read data was used. Briefly, reads were split into R1 (transcript) and R2 (barcode + UMI) read pairs based on Parse linker sequences. Cell barcodes were demultiplexed from R2 reads, and R1 reads were aligned to the GRCh38 reference genome (10x Genomics, 2024-A) using minimap2^91^ (version 2.31) with parameters –MD -a -u f -x splice. To distinguish true cell barcodes from background, a minimum transcript threshold was defined as the barcode transcript count corresponding to the steepest drop in the log-transformed barcode-rank plot. Barcodes above this threshold were classified as cells, and barcodes below this threshold were deemed as background and removed. Additionally, cells exceeding five MADs from the median for log-transformed total counts or detected genes, or showing high mitochondrial content (> 5 MADs or > 20%), were filtered out, resulting in *n* = 6, 174 total cells. Genes with fewer than 10 total UMI counts were removed Raw counts were normalized to 10,000 counts per cell and log-transformed in scanpy. For each time point, PCA was performed on HVGs and a neighborhood graph was computed with 10 nearest neighbors A UMAP embedding was constructed, and leiden clustering was performed with resolution=0.4.

#### Differential expression analysis

Differential expression analysis was performed with the scanpy.tl.rank_genes_groups over normalized expression (log-transformed) using the Wilcoxon rank-sum test (two-sided) with Benjamini-Hochberg correction. Unless stated otherwise, differential genes were identified in each cell type or cluster with the following criteria: expressed in > 30% of group; |log_2_FoldChange| > 0.5; adjusted *p*-value *<* 0.05.

#### RNA velocity inference

We developed a Snakemake workflow to preprocess BAM files for RNA velocity analysis with velocyto^35^ (version 0.17.17). Alignment files generated by the EPI2ME Labs wf-single-cell pipeline were used as input for velocyto. Pre-processed gene expression data was merged with spliced/unspliced counts from sample-specific velocyto output files. Cell barcodes were aligned, and barcodes that did not pass quality control in the gene expression data were excluded from downstream analysis. All RNA velocity analysis was carried out in scVelo^36^ (version 0.3.3). First, we computed a nearest-neighbor graph (n_neighbors=150) on the Harmony-corrected PCA space, after which moments were estimated with scv.pp.moments. Gene-specific kinetics were inferred using the dynamical model (scv.tl.recover_dynamics). Per-cell velocities were calculated with scv.tl.velocity, and confidence of velocities were calculated with scv.tl.velocity_confidence. A velocity graph was then constructed (scv.tl.velocity_graph), terminal states and latent times were inferred (scv.tl.terminal_ states, scv.tl.latent_time), and velocity-associated genes were ranked (scv.tl.rank_velocity_ genes). Stream and grid embeddings were visualized on a force-directed layout using scv.pl.velocity_ embedding_stream.

#### Pseudotemporal inference

To identify biologically-informed root cells for pseudotime inference, cell cycle phases were assigned to individual cells with scanpy.tl.score_genes_cell_cycle() using established S and G2/M gene signatures^92^. Predicted phases were compared to DNA content-based phase assignments obtained from the hashing barcodes that were added after cell sorting (Table S4). Representative root cells were selected from the control cell population. We chose root cells from both G1 and G2/M cell cycle phases based on minimum Euclidean distance from the centroid of the respective phase group in Harmony-corrected PCA space^87^. For each root, velocity psueodotime was computed using the scVelo function scv.pl.velocity_pseudotime, producing pseudotemporal orderings for G1 and G2/M, which were averaged for each cell. Aggregate pseudotemporal orderings serve as root-agnostic measure of transcriptional distance from the initial state. To analyze transcriptional dynamics along the inferred reprogramming trajectory, log-normalized gene expression was correlated with aggregated pseudotime. We computed Spearman correlation coefficients and associated p-values for each gene relative to pseudotime, followed by multiple testing correction using the Benjamaini-Hochberg method to control the false discovery rate^93^. Only genes with adjusted *p* < 0.05 were included for downstream analyses.

For diffusion-based pseudotime^37^, a diffusion map was constructed on the Harmony-corrected PCA space (rsc.tl.diffmap), and a neighborhood graph was constructed from the resulting diffusion components. For G1 and G2/M root cells, diffusion pseudotime orderings were inferred using scanpy.tl.dpt, which were averaged for each cell. Gene expression correlations with diffusion pseudotime were computed as described above.

#### Time-series integration

We evaluated multiple data integration strategies to generate a shared low-dimensional representation for pseudotime projection (Table S11). Processed count matrices from time-series cells, control fibroblasts, and day 48 reprogrammed cells were concatenated. Cell cycle phase scores were computed using scanpy tl.score_genes_cell_cycle(), and the top 3,000 HVGs were selected across datasets. PCA was performed to define an unintegrated baseline representation.

Integration was performed using Harmony^87^, scVI^30^, and scANVI^38^. For methods incorporating cell cycle correction, cell cycle phase scores were regressed out using scanpy.pp.regress_out (Harmony) or included as continuous covariates (scVI/scANVI). For scVI, we used a a zero-inflated negative binomial (ZINB) likelihood with per-gene-per-cell-dispersion, two hidden layers of 128 units, a dropout rate of 0.25, and a 24-dimensional latent representation. Models were trained for up to 1, 000 epochs with early stopping (patience=20) using GPU acceleration (NVIDIA Tesla V100 or A100). For semi-supervised integration, scANVI models were initialized from pretrained scVI models and trained using partial cell identity labels. Control fibroblasts from Library 2 (*n* = 6, 960 cells) and control/day 0 cells from the time-series dataset (*n* = 1, 341 cells) were labeled as Fibroblast, clusters R2 and R3 were labeled as D48_CD34pos_like, and all remaining cells were labeled as Unknown. scANVI models were trained for up to 50 epochs with early stopping (patience=20) and a learning rate of 1 × 10^4^. Prediction probabilities for Fibroblast and D48_CD34pos_like were extracted for downstream analyses.

To evaluate integration performance, quantitative benchmarking was performed using scib_metrics^9^ (version 0.5.9). Batch correction and biological conservation metrics were computed with benchmark Benchmarker() and compared across low-dimensional representations, with the unintegrated PCA serving as a baseline. Batch correction was assessed using “non-reprogrammed cells”, including control fibroblasts from Library 2 (*n* = 6, 960), control/day 0 cells from the time-series dataset (*n* = 1, 341), and spike-in CD34cells from Library 1 (*n* = 1, 508). As these cells represent a shared baseline state across datasets and sequencing technologies, residual variation is expected to be primarily technical rather than biological. Biological conservation was evaluated using time-series cells (days 0 − 21), which comprise defined and distinct experimental time points, ensuring that integration preserved meaningful temporal structure. Integration performance was further assessed by visualizing two-dimensional embeddings derived from each representation. Neighborhood graphs were constructed using 35 nearest neighbors, and force-directed graphs were generated using a ForceAtlas2^88^ layout.

#### Pseudotime projection

For simplicity, “query” refers to the reprogramming time-series cells and “reference” refers to the primary control and day 48 reprogrammed cells. Query cell pseudotime values were projected from the reference dataset using *k*-nearest-neighbor (kNN) transfer in each scVI/scANVI latent space. For each query cell, its *k* nearest reference cells were identified using Euclidean distance, and projected pseudotime was computed as an inverse-distance-weighted average of reference pseudotime values:

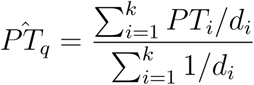

where *P*^^^*T_q_* is the projected pseudotime of query cell *q*, *PT_i_*is the pseudotime of the *i^th^* nearest reference cell, and *d_i_*is the Euclidean distance between them. To identify query cells with poor reference coverage, the mean distance to the *k* nearest reference neighbors (*d*) was compared to the distribution of internal kNN distances among reference cells. A coverage threshold was defined as:

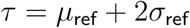

where *µ*_ref_ and *σ*_ref_ are the <u>m</u>ean and standard deviation of reference-cell internal kNN distances, respectively. Query cells with *d > τ* were excluded from downstream analyses. The association between experimental day and projected pseudotime was quantified using Spearman’s rank correlation. Correlation coefficients were computed for each latent space and value of *k*. For each experimental day, 95% confidence intervals (CIs) for the median projected pseudotime were estimated by nonparametric bootstrap resampling with replacement (1, 000 iterations). Bootstrapping was performed separately for each latent space and value of *k*, and CIs were defined by the resulting bootstrap median distribution.

Day 14 cells exhibited the lowest transcript and gene complexity among the time-series dataset (20 − 30% reduction compared to days 7 and 21; Figure S17C) and was predominantly classified as Fibroblast-like by scANVI while failing to integrate consistently with day 7 and day 21 cells across multiple integration strategies. Thus, day 14 cells were considered to be insufficiently resolved for reliable pseudotime projection and were excluded from correlation analyses.

#### Long-read scRNA-seq harmonization

We used scVI^30^ to harmonize data across the three primary long-read scRNA-seq libraries (Libraries 1 − 3). Processed UMI count matrices were concatenated and cell cycle scores were computed using scanpy.tl.score_genes_cell_cycle() with established gene sets^92^. Next, we defined a variational autoencoder with the batch identifier as a categorical covariate and S and G2/M scores as continuous covariates. We used a ZINB likelihood with per-gene–per-cell dispersion, two hidden layers of 128 units each, a dropout rate of 0.25, and a 24-dimensional latent representation. Training was performed on an NVIDIA Tesla V100 or A100 for up to 1,000 epochs with early stopping (patience=10). Next, we performed PCA on the scVI latent space, and computed a neighborhood graph using 35 neighbors Cluster connectivity was inferred using scanpy.tl.paga^96^, grouping cells by their annotated labels. The resulting Partition-based Graph Abstraction (PAGA) coordinates were used to initialize a two-dimensional UMAP embedding with parameters spread=0.55 and min_dist=0.2.

Pairwise Euclidean and cosine distances were used to quantify similarity between reprogrammed cells and initial or target populations in the 24-dimensional latent representation. For each reprogrammed cell, distances were calculated to all cells annotated as initial or HSC (cuml.metrics.pairwise_distances) and averaged. Per-cell average distances were grouped by cluster (R1-R3) and visualized. For visual interpretation of distances, “close” and “far” reference thresholds were calculated as the mean pairwise distance between HSCs and hematopoietic progenitors (“close”) and between HSCs and initial fibroblasts (“far”).

#### Integrated atlas benchmarking

We compiled scRNA-seq data from four sources: the native reference atlas, Ng 2024^1^, Gomes 2018^10^, and this study, yielding 983,455 cells and 105 annotated cell types. After excluding immune populations, the reference atlas dataset contributed 709,808 cells across 72 cell types. The Ng 2024 dataset provided 252,607 cells spanning 20 types, Gomes 2018 contributed 286 cells from five populations, and this study added 20,754 cells across 12 clusters. All datasets were filtered to remove unwanted cell types and non-coding genes, and genes with fewer than 10 total counts were excluded. Up to 250,000 cells were randomly sampled per dataset, retaining relevant metadata fields where available. After sampling, 521,040 cells and 17,607 genes remained. To ensure robust downstream integration, batch labels with fewer than 100 cells were excluded. Additional filtering removed cells with fewer than 500 total counts (17,783 cells) and genes with low or zero counts (234 genes). Cells with fewer than 1,000 detected genes were also removed. The final dataset (Figure S23) contained 420,108 cells and 17,373 genes, which were normalized to 10,000 total counts per cell and log-transformed. Cell-cycle phase scores were predicted for each cell as described above. HVGs were selected using the seurat_v3 method, retaining the top 3,000 features. Genes annotated as mitochondrial or ribosomal were excluded from HVG selection.

To harmonize single-cell profiles, a scVI model was trained using the data source as the batch key and including continuous covariates for cell cycle phase (S and G2M scores). The model architecture included two encoder/decoder layers and 128 hidden units each with a dropout rate of 0.15, and a 24-dimensional latent representation. Gene expression counts were modeled with a ZINB likelihood and per-gene-per-cell dispersion parameters. Training was performed on either an NVIDIA Tesla V100 or A100 for up to 500 epochs with early stopping (patience=30), a learning rate of 1 × 10*^−^*^3^, and a batch size of 500. Model performance was monitored by tracking training and validation loss, evidence lower bound (ELBO), reconstruction error, and Kullback–Leibler (KL) divergence. We performed PCA on the scVI latent space, computed a neighborhood graph (n_neighbors=15), and generated a UMAP embedding (min_dist=0.2). Leiden clustering was performed with resolution=0.45.

To compute the abstract connectivity graph of cell clusters, we applied PAGA^96^ using scanpy.tl paga, and visualization was generated with scanpy.pl.paga. Pairwise Euclidean distances were used to compare pseudobulk profiles of cell clusters in the latent space using cuml.metrics.pairwise_ distances, where pseudobulk profiles were obtained by taking the mean of the latent vector per cluster. To compare single-cell profiles, Euclidean distances were computed between each cell and the mean vector of R2/R3 cells in the latent space using linalg.norm from CuPy^97^ (version 13.5.1). Per-cell distances were grouped by cluster-resolved cell type annotations.

#### Curated dataset benchmarking

We benchmarked our scRNA-seq data on six publicly available scRNA-seq datasets (Table S6). Briefly, we obtained 359,848 single cells, and a broad spectrum of 71 annotated cell types from^1,10,19,21,39,98^ For each dataset, pseudobulk profiles were obtained by summing over raw UMI counts for each source-defined cell type label. Aggregated counts were normalized using scanpy.tl.normalize_total with target_sum=10^4^ and log-transformed using scanpy.tl.log1p. Cell-cycle phase scores were predicted for each pseudobulk profile as described above. We filtered genes annotated as mitochondrial, ribosomal, non-protein-coding, or cell-cycle related^92^, and only genes whose mean expression lay in the upper quartile in both the test dataset and the curated datasets were retained, yielding 15,216 shared genes across 82 pseudobulk profiles. Pseudobulk profiles were harmonized as described for integrated atlas benchmarking with a dropout rate of 0.25, batch size of 25, and a 90/10 train/validation split. PCA was then computed on the harmonized latent space. Pairwise Euclidean distances were used to compare pseudobulk profiles in the PCA representation of the latent space using (cuml.metrics.pairwise_distances).

#### Partial identity scoring

We used Capybara^22^ to estimate the fractional identity scores of single-cells based on aggregated pseudobulk profiles. Briefly, we used the intersection of gene sets between the input data and the reference pseudobulk profiles. Identity scores were computed using single.round.QP.analysis with 24 CPU cores, log-transformed and scaled both bulk and single-cell counts, and forced no equality constraints (force.eq=0). Resulting scores were stored for downstream analysis.

#### Isoform identification

Transcript-level UMI count matrices were generated with the EPI2ME Labs wf-single-cell pipeline (github.com/epi2me-labs/wf-single-cell) using default parameters. Sequencing reads were aligned to the GRCh38 reference genome (10x Genomics, 2024-A) prior to quantification.

Reads were assigned to transcripts following a strategy adapted from FLAMES^99^. Alignments mapping to intronic query transcripts were labeled as unknown (“--”). Reads with a single, unique alignment to the annotated transcriptome were directly attributed to that transcript. For ambiguously mapped reads, alignments were ordered by alignment score (AS); if the top hit exhibited a higher AS or greater query coverage than the second hit and covered at least 40% of the transcript, the read was assigned accordingly, otherwise it was marked unknown. When the top two alignments shared identical AS values, the alignment with the greater transcript coverage was selected provided its coverage exceeded 80%. After filtering out cell barcodes that failed gene-expression quality control, the final dataset is comprised of 20,754 cells across three libraries and 113,335 detected transcripts. For downstream analyses, transcripts with fewer than 30 total UMI counts were filtered out in Scanpy. Raw counts were normalized to 10,000 counts per cell and log-transformed. Protein identifiers were assigned to transcripts by mapping Ensembl transcript IDs to UniProt accessions from HUMAN_9606_idmapping_selected.tab.gz.

#### Splicing event annotation

AS events were annotated with SUPPA^100^ (version 2.4) using the GRCh38_2024-A gene annotation. Analyses were run in intron-over-intron (ioi) and intron-over-exon (ioe) modes with gene pooling enabled; the ioe run was restricted to the SE, SS, MX, RI, and FL event classes (Table S12).

### Quantification and statistical analysis

Statistical analyses were performed using GraphPad Prism (version 10.4.1) or SciPy^101^ (scipy.stats, version 1.16.1). All statistical tests, sample sizes, and replicate definitions are described in corresponding Methods sections or in the main text and figure legends. Statistical tests included two-sided Welch’s *t*-test, one-sided Mann-Whitney U test, Wilcoxon rank-sum test, and Spearman or Pearson correlation The Benjamini-Hochberg method was used for multiple testing correction. Unless stated otherwise, *p* values or FDR-adjusted *p* values *<* 0.05 were considered statistically significant and were indicated as follows: *p <* 0.05 (*), *p <* 0.01 (**). Unless otherwise specified, boxes show the median and IQR; the lower whisker indicates Q11.5 × IQR; the upper whisker indicates Q3 + 1.5 × IQR.

