## Supplementary material for "Transcriptional landscape of direct reprogramming toward the hematopoietic lineage": Document S1

### 1 Supplemental Material

#### 1.1 Supplementary Figures

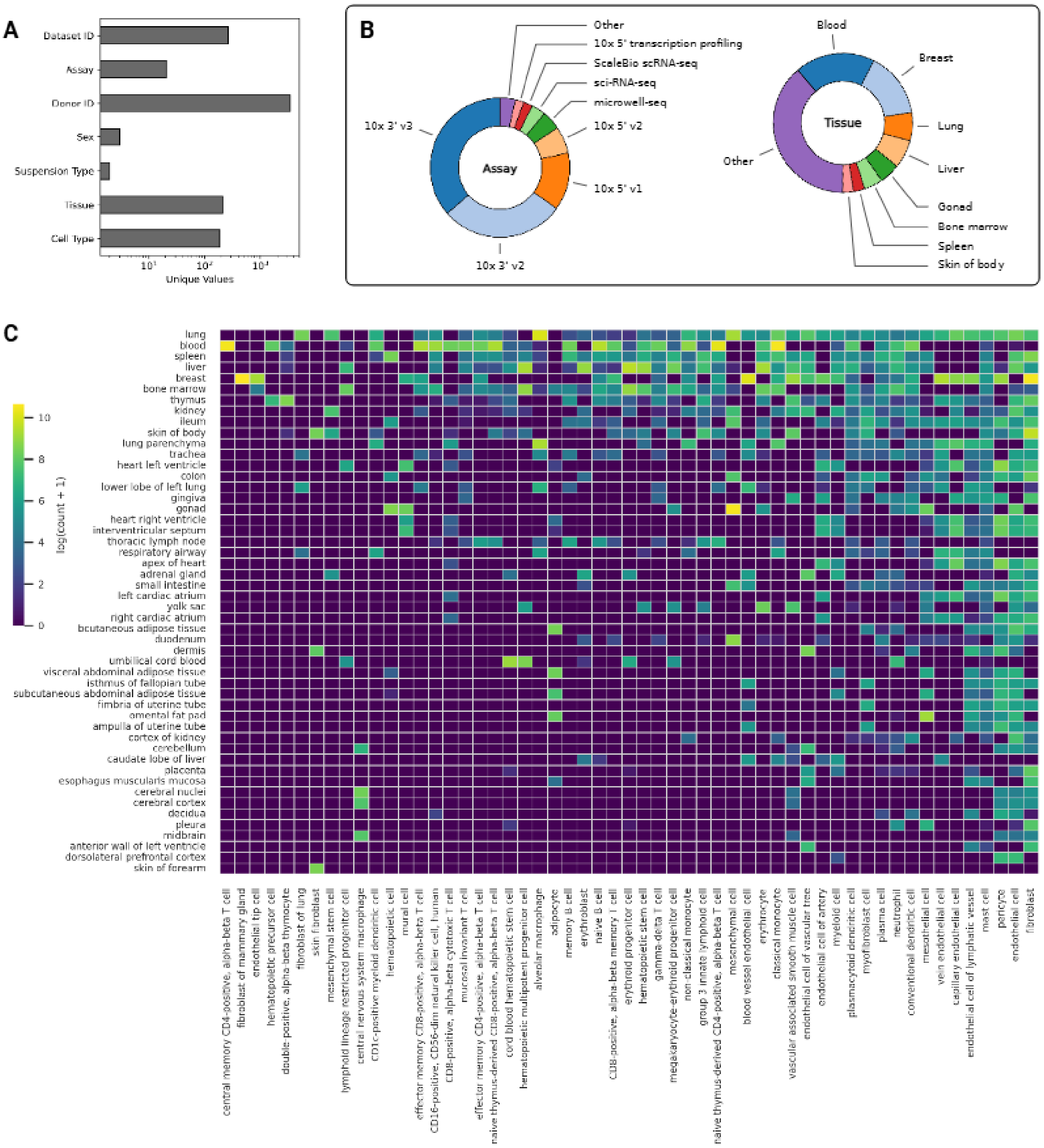


Figure S1: **Overview of tissue, cell type, and metadata composition in the reference atlas. (A)** Number of unique values per metadata field (log scale). **(B)** Distribution of tissue sources and assay modalities. **(D)** Heatmap showing log-transformed counts of cells ($\log(\text{count} + 1)$) for the most abundant tissues and cell types across the dataset. Tissues (rows) and cell types (columns) were selected to collectively represent 90% of the total counts in each category.

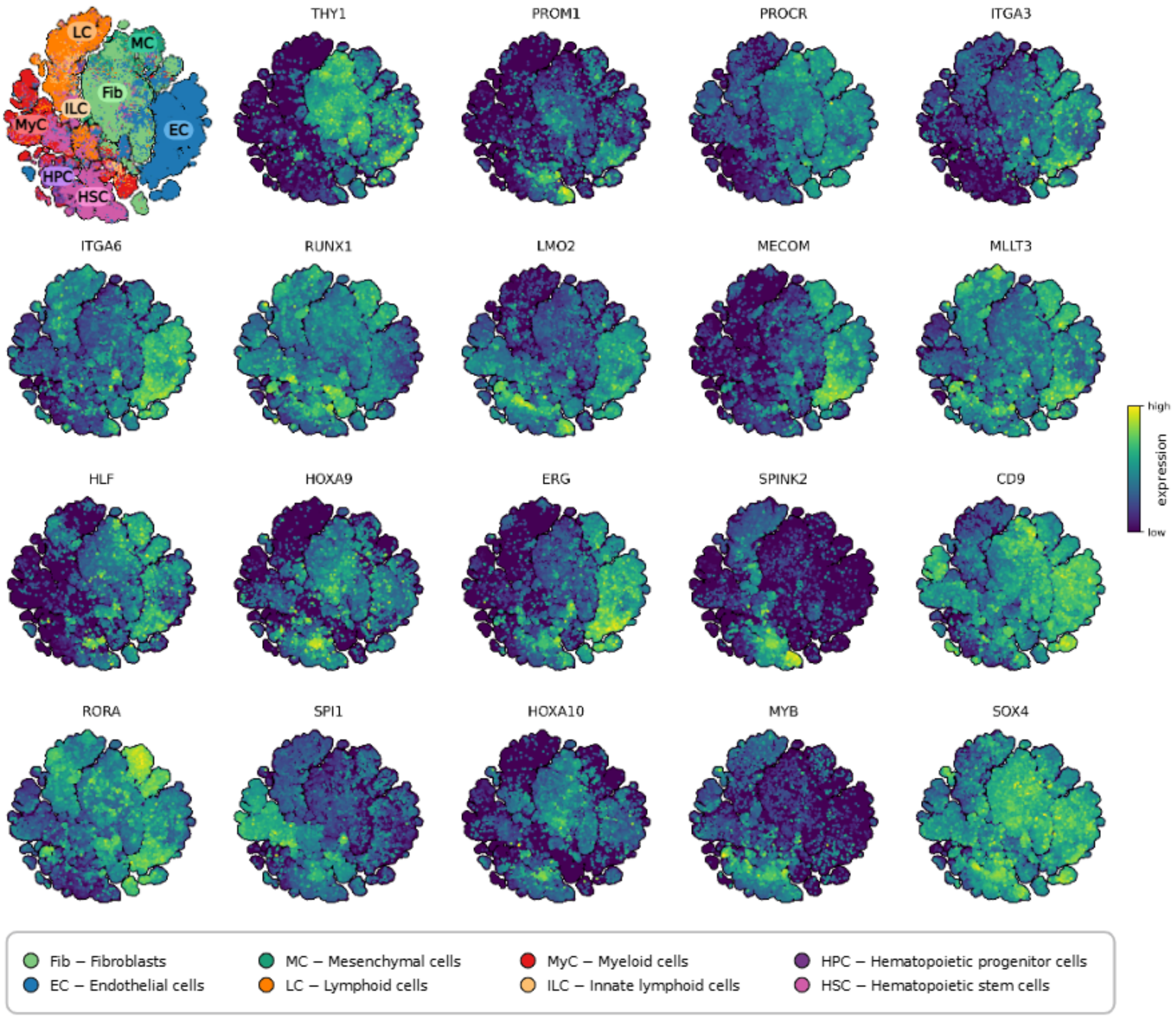


Figure S2: **Gene expression across reference atlas.** Expression patterns of selected genes across all cell types, visualized in a shared low-dimensional (t-SNE) embedding. Each panel shows log-normalized expression of a single gene.

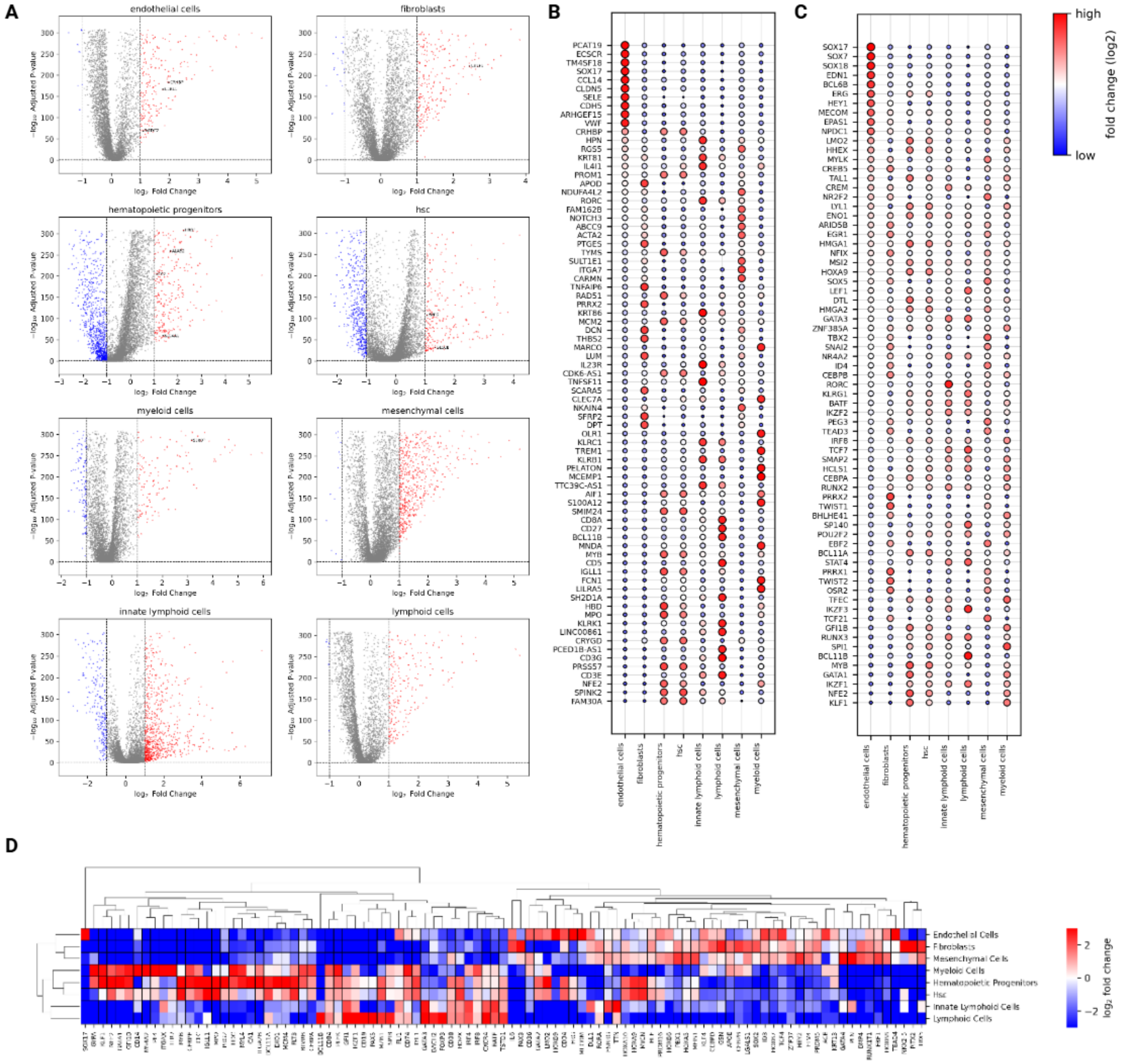


Figure S3: **Differential expression of major cellular compartments.** **(A)** Volcano plots for DEGs for each group (Wilcoxon rank-sum test). Genes with adjusted $p$-value $< 0.05$ and absolute $\log_2$ fold change $> 1$ were considered significant (red: upregulated, blue: downregulated). Dashed lines indicate the significance thresholds. Marker genes derived from Tabula Sapiens[1]. **(B)** The top 10 DEGs per group, filtered to include only genes expressed in at least 15% of cells, with color and size indicating $\log_2$ fold change. **(C)** The top 10 differentially expressed TFs per group, filtered to include only genes expressed in at least 15% of cells, with color and size indicating $\log_2$ fold change. **(D)** Differential expression of literature-curated genes across cell types. Genes filtered significant differential expression (adjusted $p$-value $<$ 0.05) and high variability (standard deviation $>$ 1.5) in $\log_2$ fold change. The heatmap shows average expression changes across annotated cell types, clustered by gene and cell identity.

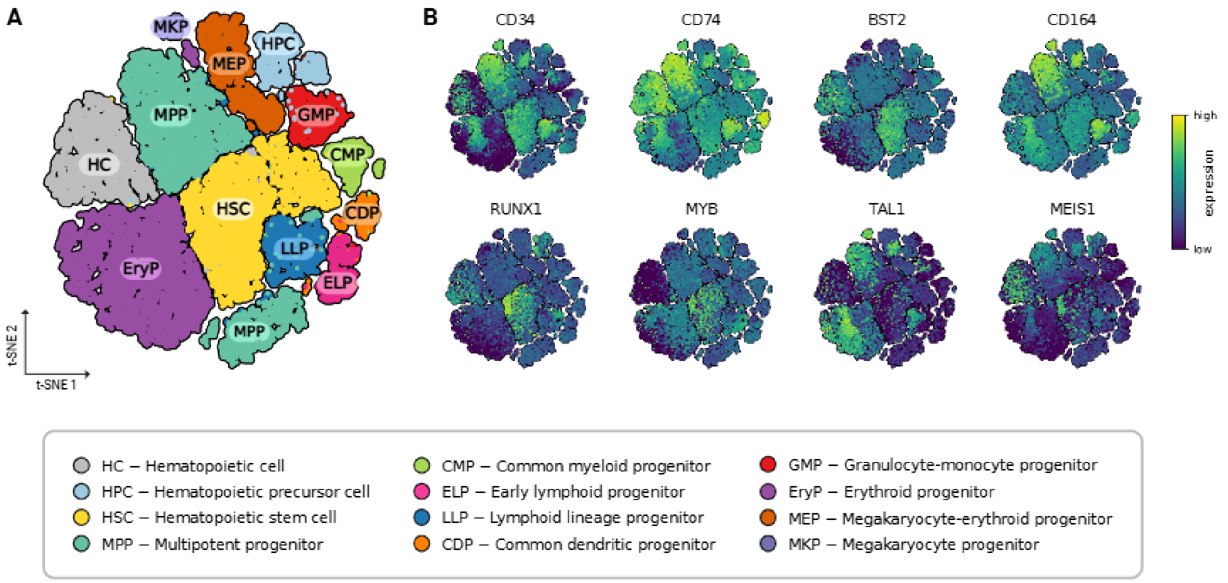


Figure S4: **Marker gene expression in hematopoietic cells from the reference atlas. (A)** t-SNE of major hematopoietic cell types. **(B)** Expression of marker genes on coordinates from (A).

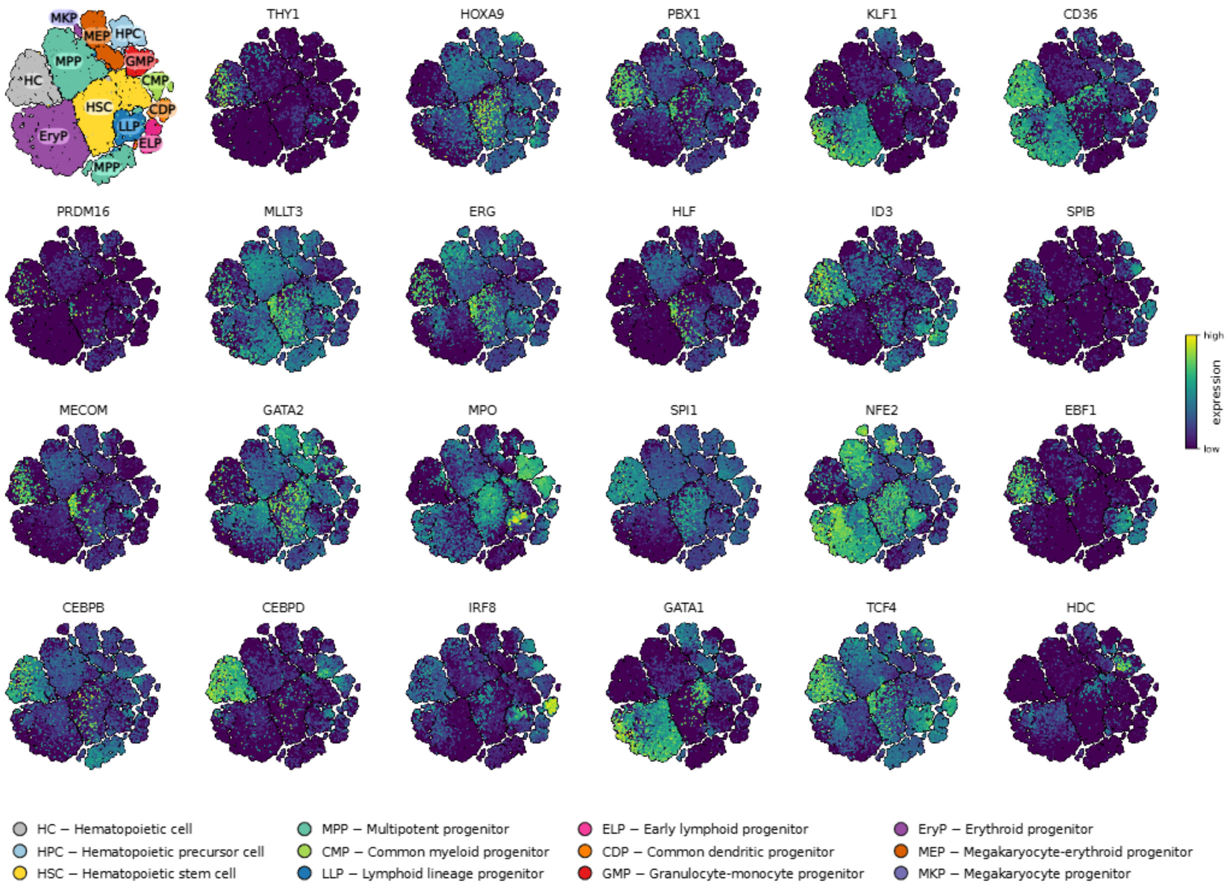


Figure S5: **Gene expression across hematopoietic populations in the reference atlas.** Expression patterns of selected genes across hematopoietic cell types, visualized in a shared low-dimensional (t-SNE) embedding. The top left t-SNE plot is colored by cell type. All other t-SNE plots show log-normalized expression of a single gene.

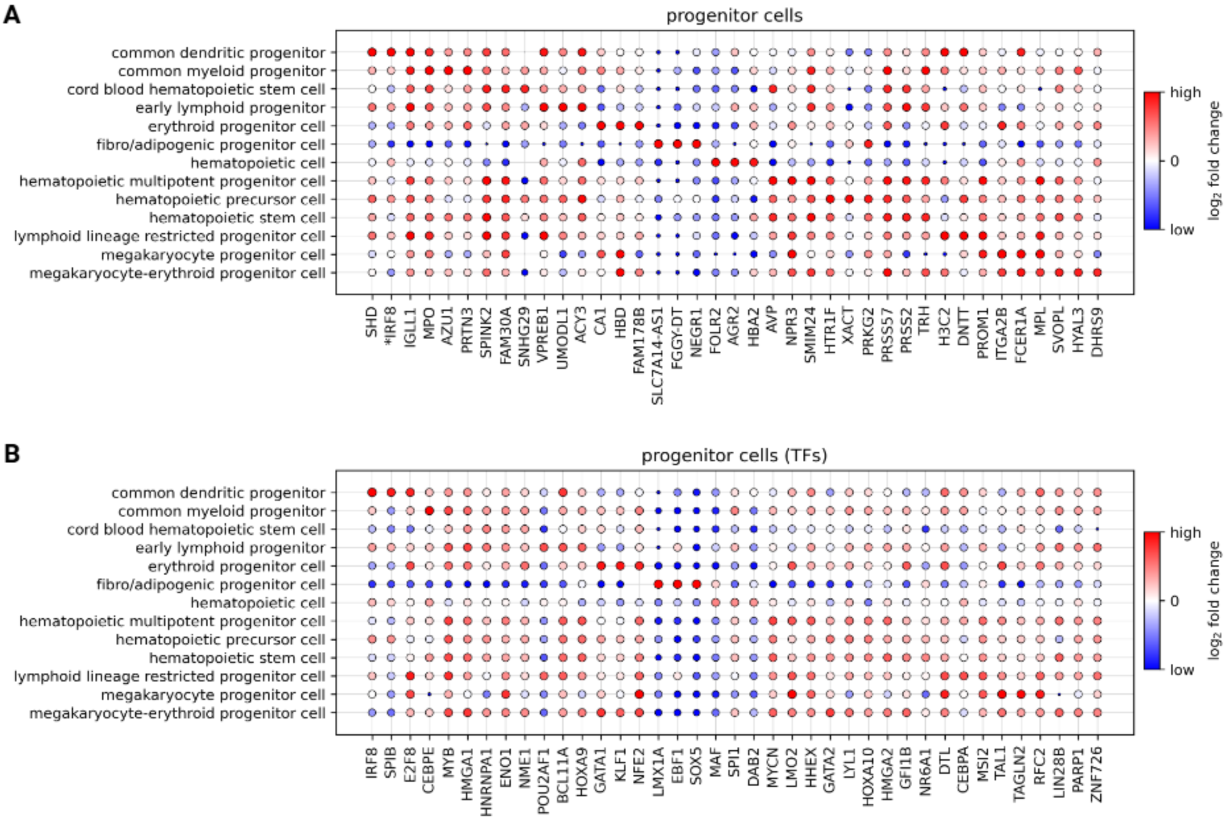


Figure S6: **Differentially expressed genes across hematopoietic progenitor subtypes.** The top 3 unique **(A)** DEGs and **(B)** differentially expressed TFs for each cell type. The size and color of each dot represents $\log_2$ fold change. Asterisks in (A) denote TFs.

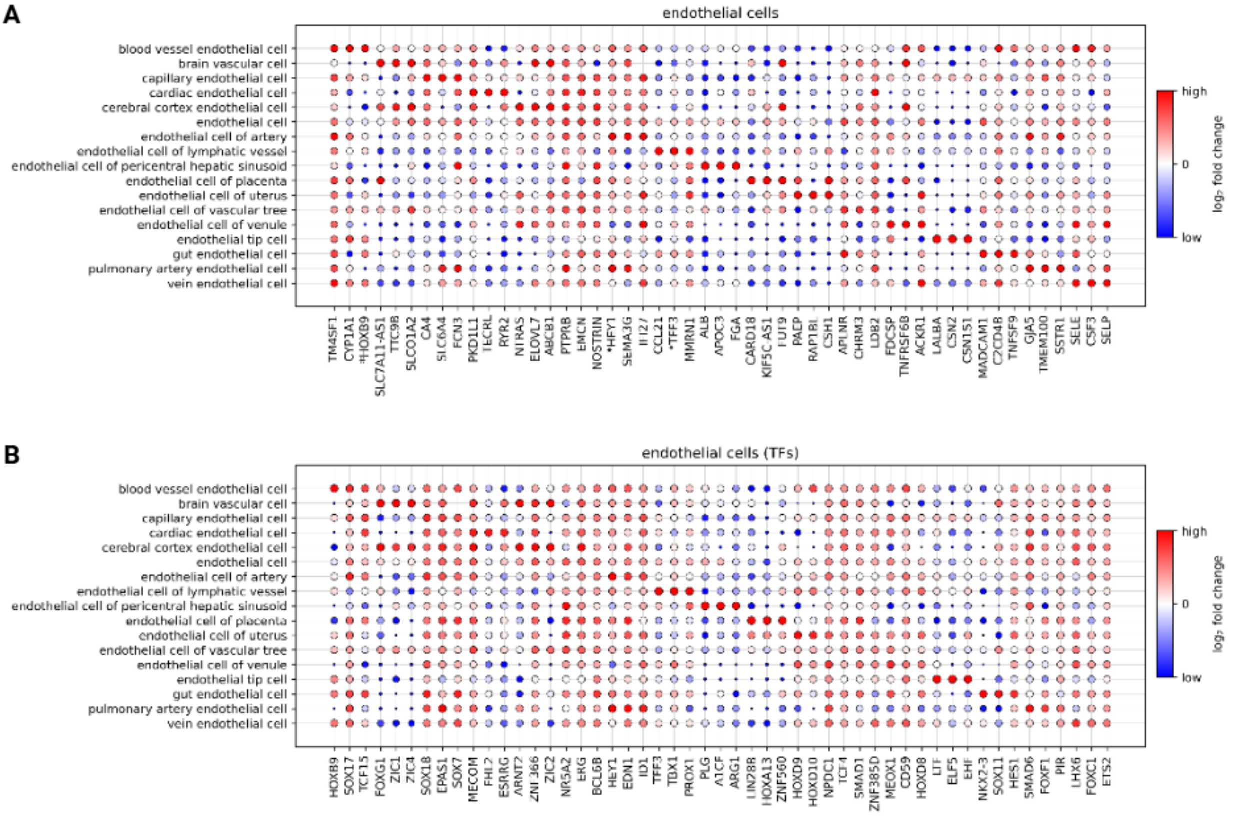


Figure S7: **Differentially expressed genes across endothelial cell subtypes.** The top 3 unique **(A)** DEGs and **(B)** differentially expressed TFs for each cell type. The size and color of each dot represents $\log_2$ fold change. Asterisks in (A) denote TFs.

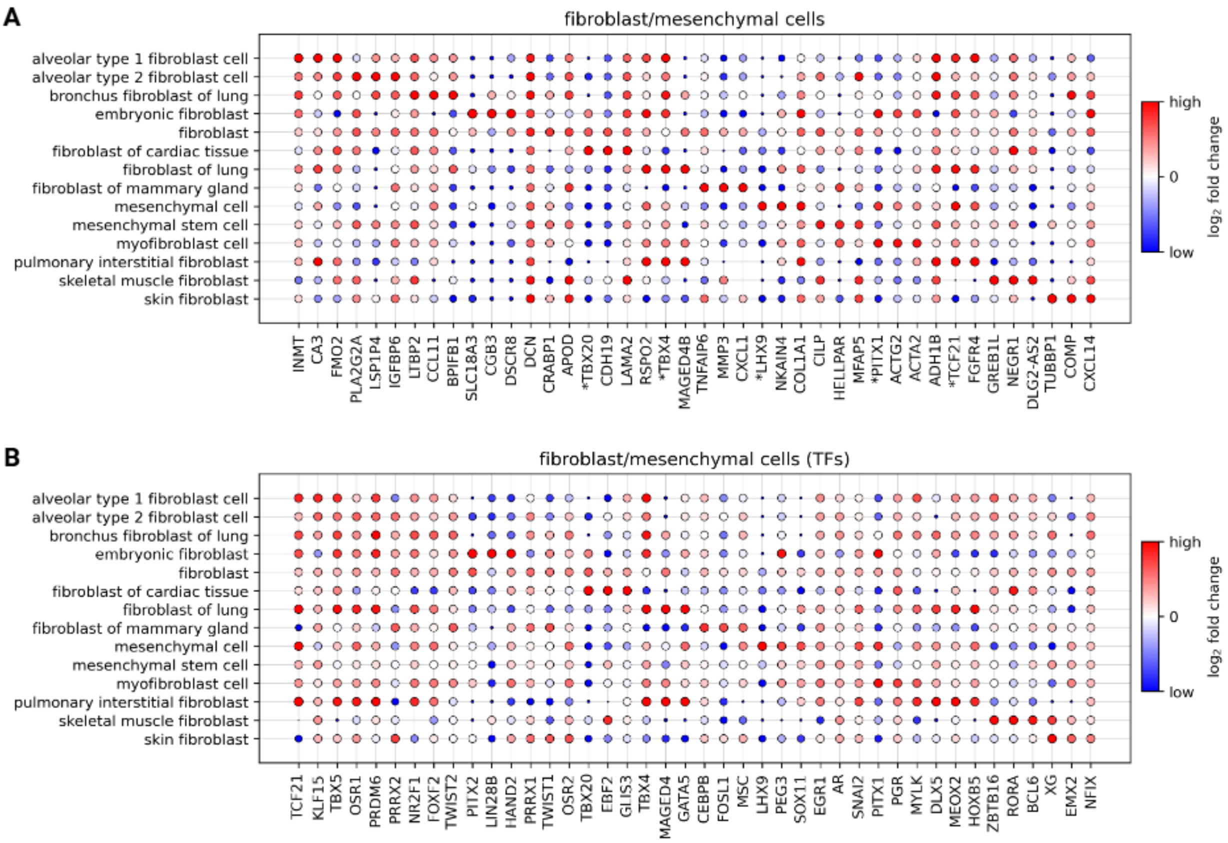


Figure S8: **Differentially expressed genes across fibroblast and mesenchymal subtypes.** The top 3 unique **(A)** DEGs and **(B)** differentially expressed TFs for each cell type. The size and color of each dot represents $\log_2$ fold change. Asterisks in (A) denote TFs.

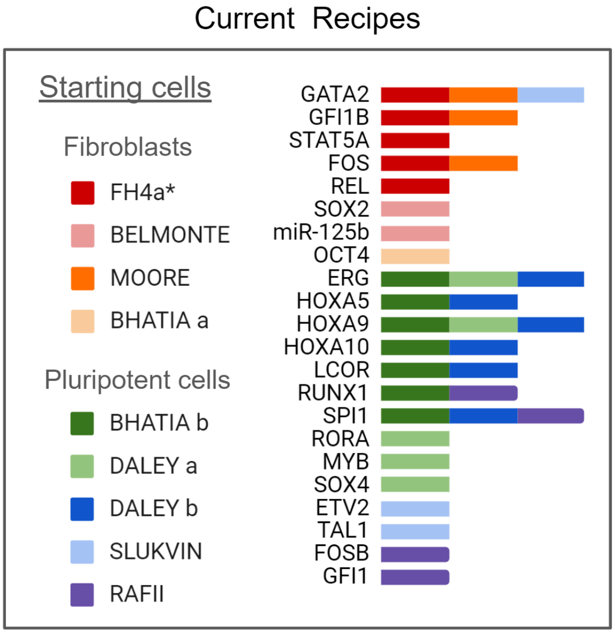


Figure S9: **Transcription factor recipes used in reprogramming human cells into HSCs**. Left: Recipes named by group and divided by the starting cell type. Right: TFs used in each recipe. *FH4a (and FH6) describe the 5TF recipe used throughout this manuscript.

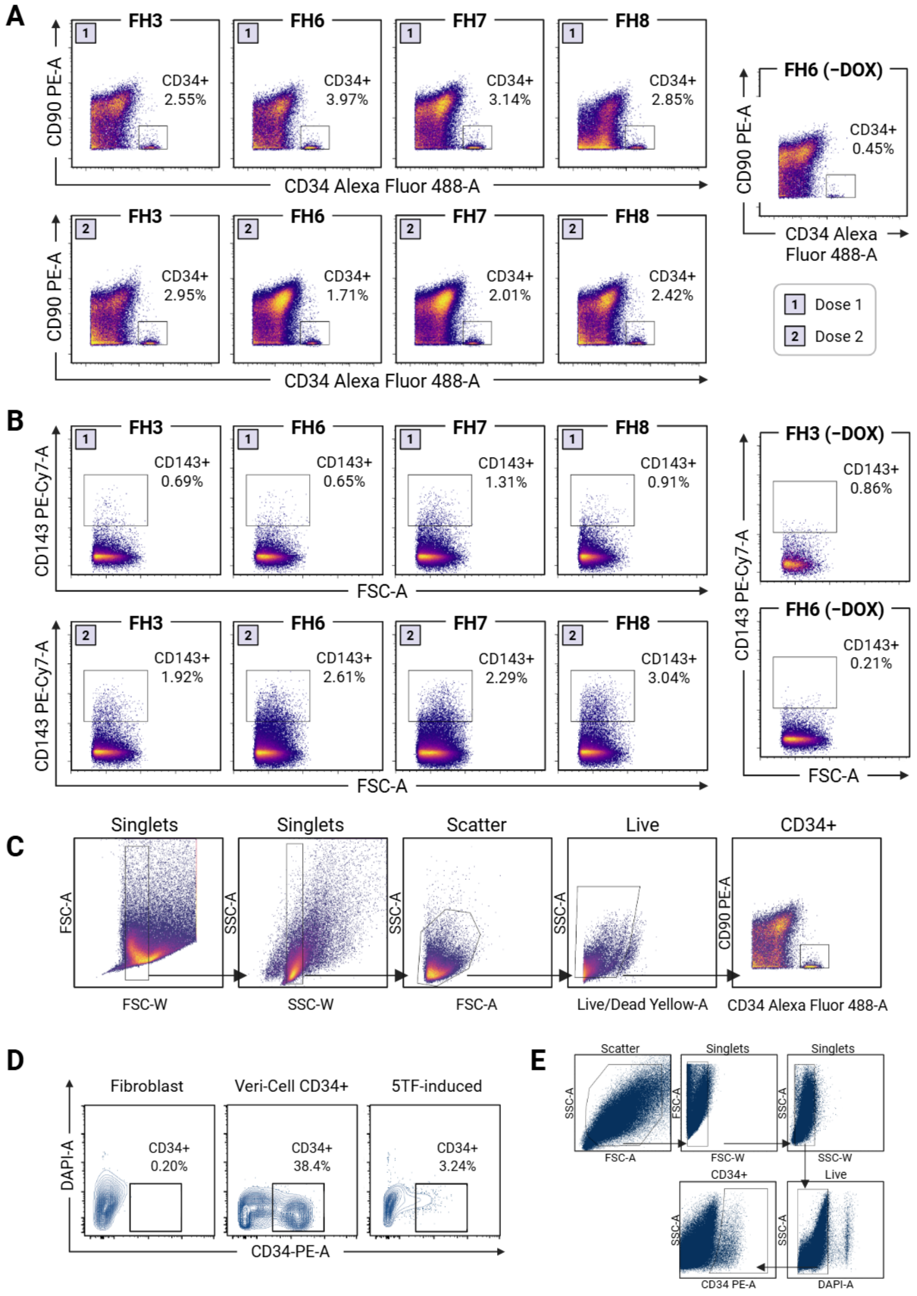


Figure S10: **Flow cytometry analysis of multiple TF recipes.** Different combinations of transcription factors in the 5TF recipe were tested for their ability to generate CD34$^+$ (A) and CD143$^+$ (B) cells. **(A)** Identification of CD34$^+$ cells with TF recipes (Table S7) delivered at low and high concentrations, corresponding to Dose 1 and Dose 2, respectively. FH6-transduced fibroblasts without Dox treatment served as negative controls. FH7 (GATA2, GFI1B, FOS, STAT5A) and FH8 (GATA2, GFI1B, FOS, REL) were prepared according to Method  as described for FH3/FH6. Recipe combinations were made by mixing equal volumes of the corresponding lentiviral preparations. High and low dose amounts were based on identically prepared control fluorescent vectors. **(B)** Identification of CD143$^+$ cells. Right: FH6- and FH3-transduced fibroblasts without Dox treatment served as negative controls. **(C)** Gating scheme for hematopoietic markers, with the final gate shown for CD34. **(D)** Flow cytometry plots generated during flow sorting of the reprogramming cultures used for scRNA-seq. Left: BJ fibroblast negative control. Middle: Veri-Cell CD34$^+$ PBMCs (Biolegend) as a positive/gating control. Right: reprogramming cultures 48 days after initial Dox treatment, showing CD34$^+$ iHSCs. **(E)** Gating scheme for cell sorting.

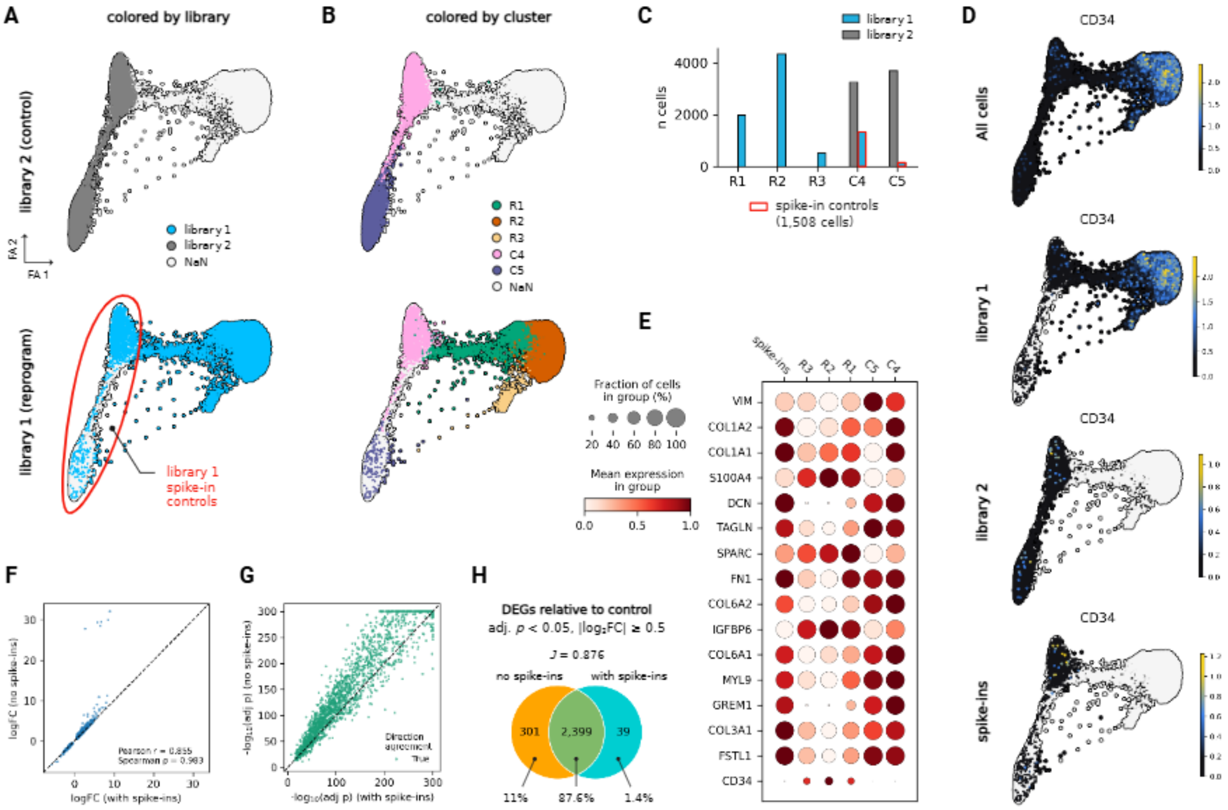


Figure S11: **Identification of spike-in CD34$^-$ controls from library 1 (reprogram). (A)** Force-directed graph layout from Figure 2 colored by library and **(B)** by Leiden cluster for control (top) and reprogrammed (bottom) cells. **(C)** The number of cells in each cluster colored by library, with red boxes indicating the spike-in controls. **(D)** Log-normalized *CD34* expression on the graph from (A). Top to bottom: all cells from both libraries, only cells from library 1 (reprogram), only cells from library 2 (control), spike-in control cells (library 1 cells overlapping with control clusters). **(E)** Mean log-normalized expression of *CD34* and the top 15 fibroblast marker genes expressed in controls by cluster (spike-ins plotted separately). **(F)** Comparison of $\log_2$ fold change (logFC) values for significant DEGs between reprogrammed and control cells when spike-in controls were excluded versus included in the control population (Wilcoxon Rank-sum test, corrected $\alpha = 0.05$, $|\log_2$ fold change$| > 0.5$, expressed in $> 30\%$ of cells where $\log_2$ fold change $> 0$). Each point represents a shared DEG; the dashed line denotes identity. Pearson ($r$) and Spearman ($\rho$) correlation coefficients are shown. **(G)** Comparison of statistical significance (-$\log_{10}$(adjusted $p$-value)) for DEGs from (F). Points are colored by agreement in direction of differential expression between analyses; the dashed line denotes identity. **(H)** Overlap of DEGs from (F). Percentages indicate the fraction of total unique DEGs across both analyses; Jaccard index ($J = 0.876$).

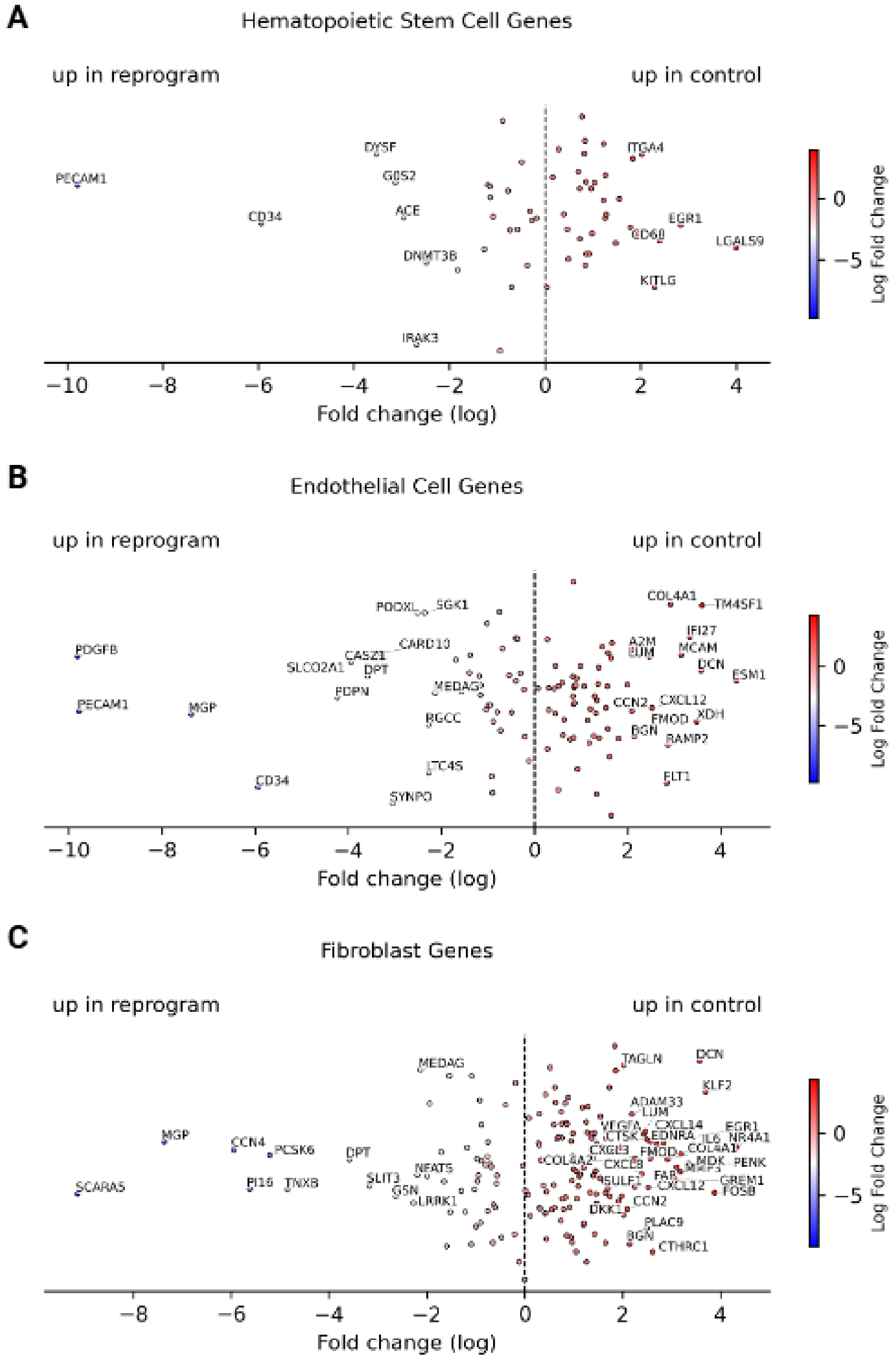


Figure S12: **Differentially expressed marker gene sets.** Differential expression results for marker gene sets from PanglaoDB[2] between reprogrammed cells and controls (Wilcoxon rank sum, corrected $\alpha = 0.05$). All plots highlight genes with absolute $\log_2$ fold change $> 2$. **(A)** HSC genes. **(B)** Endothelial genes. **(C)** Fibroblast genes.

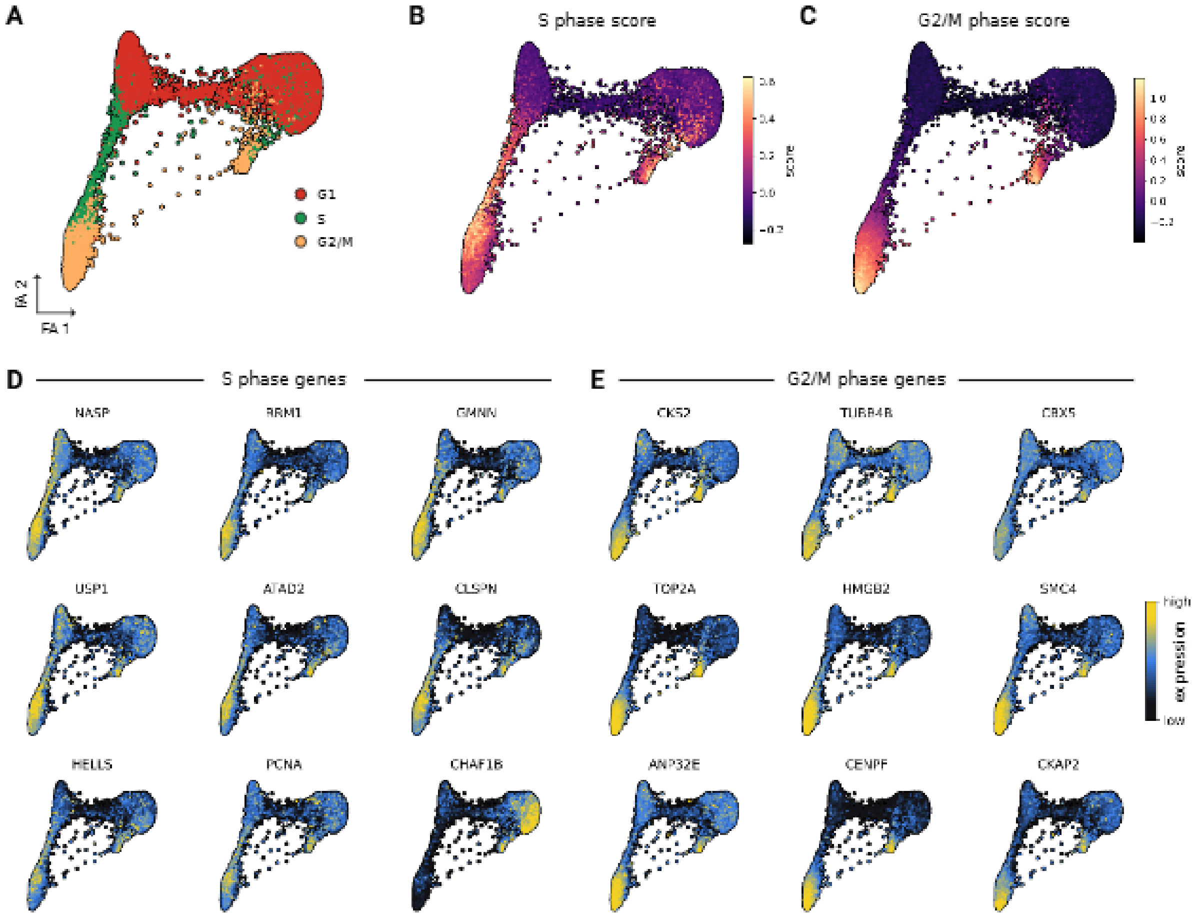


Figure S13: **Cell cycle phase analysis for control and reprogrammed cells.** **(A)** Predicted cell cycle phase for all cells. **(B)** S-phase scores and **(C)** G2/M-phase scores. Scores were computed using the `scanpy.tl.score_genes_cell_cycle` function with gene sets from[3]. **(D)** Log-normalized expression of top expressed S-phase genes. **(E)** Log-normalized expression of top expressed G2/M-phase genes.

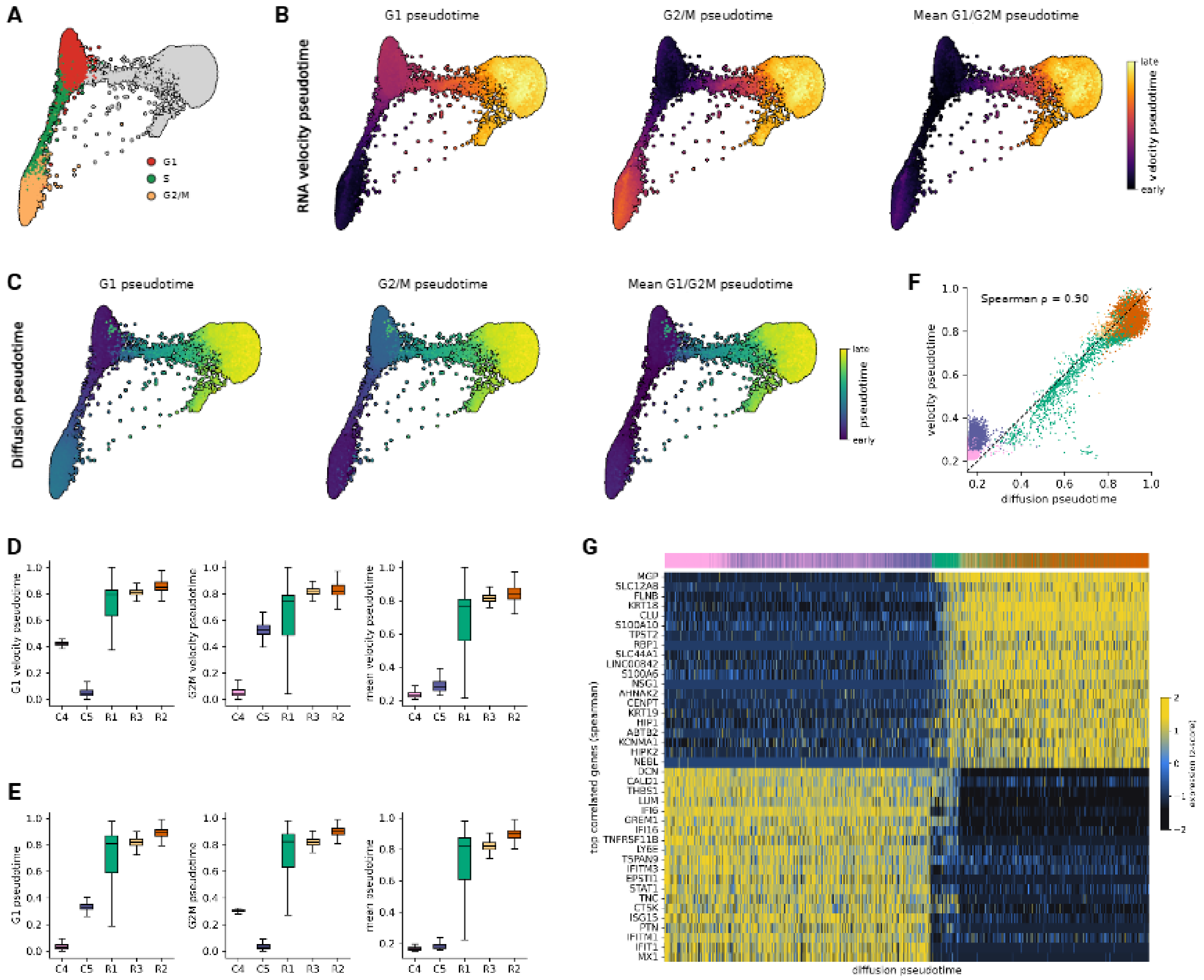


Figure S14: **Comparison between diffusion-based pseudotime and RNA velocity pseudotime.** **(A)** Predicted cell cycle phase for cells from Library 2 (control) used for selecting root cells in pseudotemporal orderings. **(B)** Cells colored by RNA velocity pseudotime with root cells in G1, G2/M, and averaged G1 and G2/M (left to right). **(C)** Cells colored by diffusion-based pseudotime with root cells in G1, G2/M, and averaged G1 and G2/M (left to right). **(D)** Per cluster distributions of velocity-based pseudotemporal orderings and **(E)** diffusion-based pseudotemporal orderings (Data are represented as median and IQR, with whiskers extending to $1.5 \times$ IQR). **(F)** Correlation between RNA velocity pseudotime and diffusion-based pseudotime (Spearman) colored by cell cluster (Spearman $\rho = 0.90$). Dashed line indicates identity. **(G)** Z-scored expression of top positively and negatively correlated genes with diffusion-based pseudotime. Cells (columns) are ordered by diffusion pseudotime and annotated by cluster (color bar, top). Correlations were computed using Spearman's $\rho$, and significance was assessed by Benjamini-Hochberg FDR correction ($\alpha = 0.05$). Only genes with adjusted $p < 0.05$ were considered.

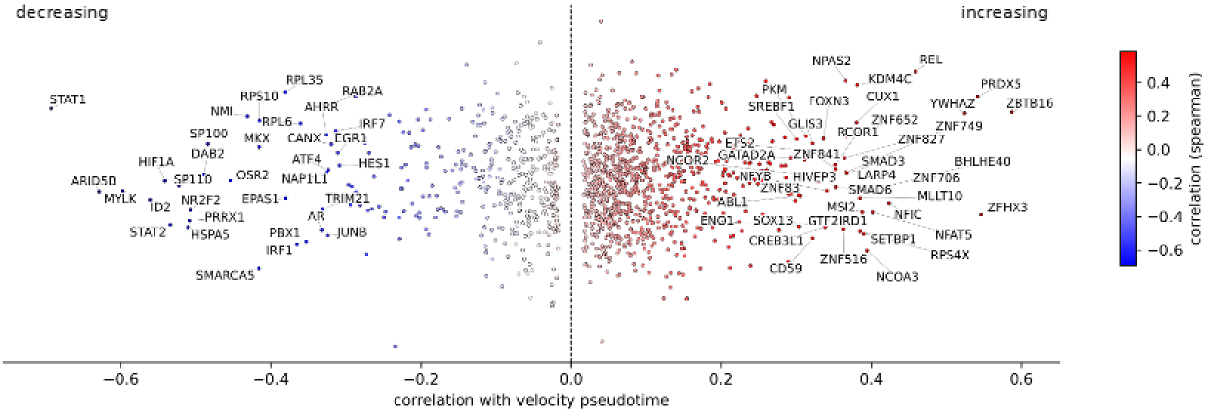


Figure S15: **Correlation of transcription factor expression with RNA velocity pseudotime.** Each point represents a TF, colored by its Spearman correlation with velocity pseudotime. Only TFs with Benjamini-Hochberg FDR-adjusted $p < 0.05$ are shown. Jitter is added along the $y$-axis for visual clarity. TFs with $|\rho| > 0.3$ are labeled.

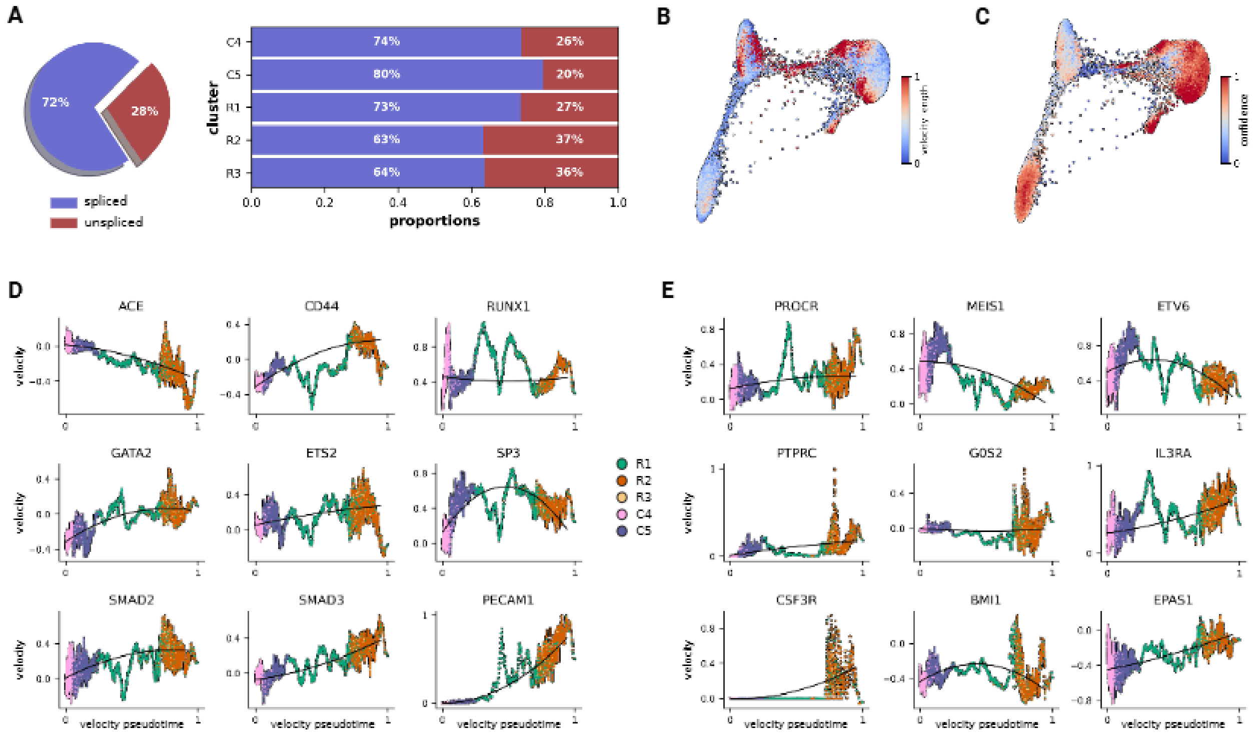


Figure S16: **Extended RNA velocity analysis.** **(A)** Spliced and unspliced transcript proportions by cluster. **(B)** Length of velocity vectors indicative of the rate of differentiation. **(C)** Confidence of velocities indicative of where vector direction is un-/determined. **(D)** Velocity shown over velocity pseudotime for endothelial/EHT-related genes and **(E)** HSC genes colored by Leiden cluster. Velocity values represent the inferred rate of change of gene expression for each gene in each cell, with positive values indicating increasing expression.

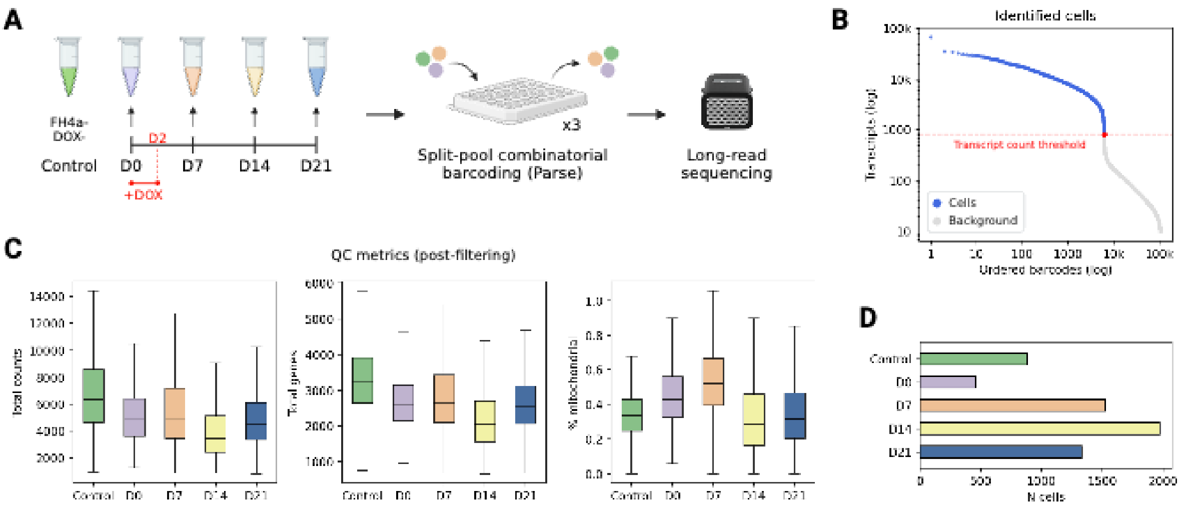


Figure S17: **Time-series reprogramming dataset collection and quality control.** **(A)** 5TF cells were collected on days 0, 7, 14, and 21 of reprogramming. A sample of non-transduced Dox- fibroblasts served as an additional control. Single cells were barcoded via split-pool combinatorial barcoding and sequencing on the ONT platform. **(B)** Barcode rank plot showing the transcript count threshold used to distinguish cells from background. **(C)** Quality control metrics after removal of low quality cells for each time point (Data are represented as median and IQR, with whiskers extending to $1.5 \times$ IQR) **(D)** The total number of cells per time point post-filtering.

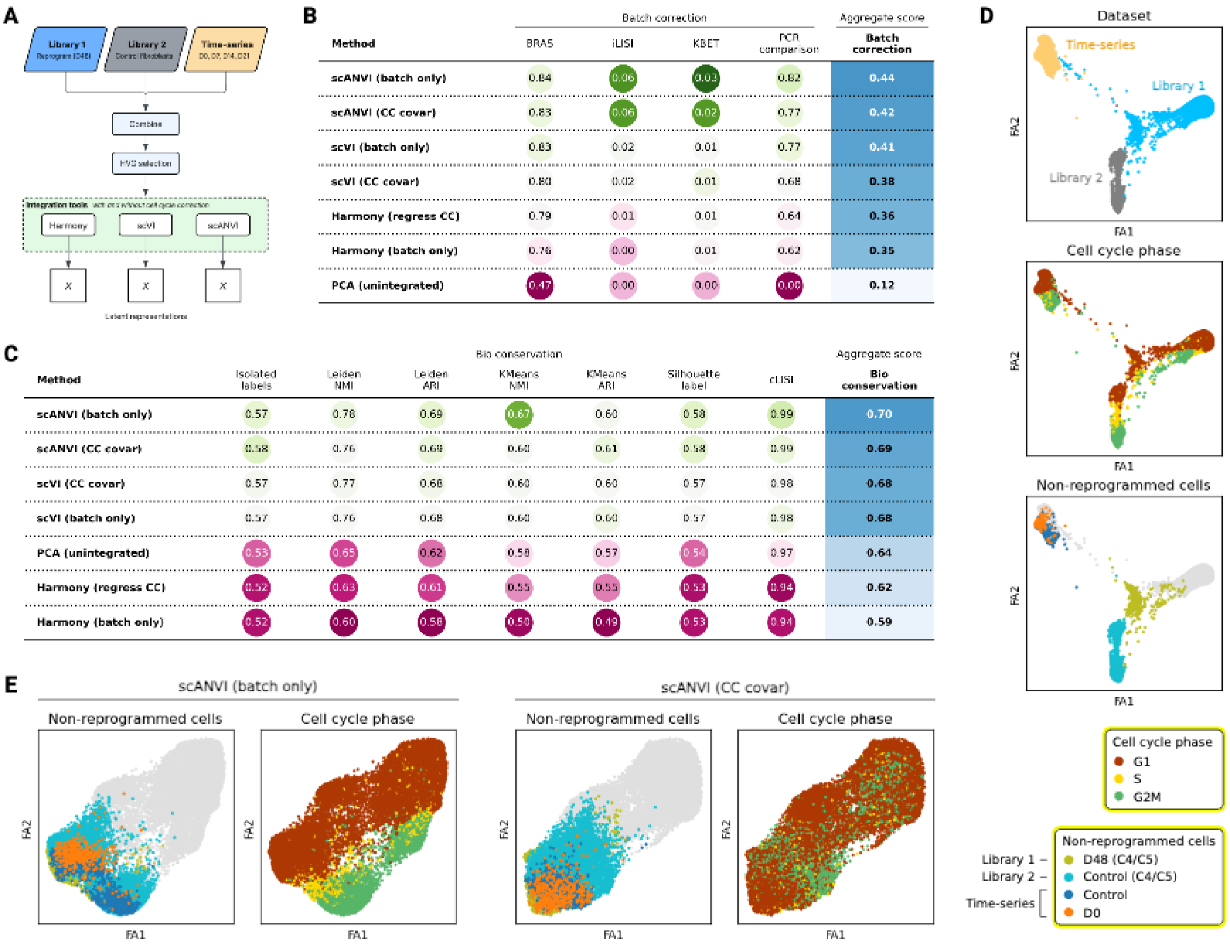


B

| Method | BRAS | iLISI | KBET | PCR comparison | Batch correction |
|---|---|---|---|---|---|
| | Batch correction | | | | Aggregate score |
| scANVI (batch only) | 0.84 | 0.06 | 0.03 | 0.82 | 0.44 |
| scANVI (CC covar) | 0.83 | 0.06 | 0.02 | 0.77 | 0.42 |
| scVI (batch only) | 0.83 | 0.02 | 0.01 | 0.77 | 0.41 |
| scVI (CC covar) | 0.80 | 0.02 | 0.01 | 0.68 | 0.38 |
| Harmony (regress CC) | 0.79 | 0.01 | 0.01 | 0.64 | 0.36 |
| Harmony (batch only) | 0.76 | 0.00 | 0.01 | 0.62 | 0.35 |
| PCA (unintegrated) | 0.47 | 0.00 | 0.00 | 0.00 | 0.12 |

C

| Method | Isolated labels | Leiden NMI | Leiden ARI | KMeans NMI | KMeans ARI | Silhouette label | cLISI | Bio conservation |
|---|---|---|---|---|---|---|---|---|
| | Bio conservation | | | | | | | Aggregate score |
| scANVI (batch only) | 0.57 | 0.78 | 0.69 | 0.67 | 0.60 | 0.58 | 0.99 | 0.70 |
| scANVI (CC covar) | 0.58 | 0.76 | 0.68 | 0.60 | 0.61 | 0.58 | 0.99 | 0.69 |
| scVI (CC covar) | 0.57 | 0.77 | 0.68 | 0.60 | 0.60 | 0.57 | 0.98 | 0.68 |
| scVI (batch only) | 0.57 | 0.76 | 0.68 | 0.60 | 0.60 | 0.57 | 0.98 | 0.68 |
| PCA (unintegrated) | 0.53 | 0.65 | 0.62 | 0.58 | 0.57 | 0.54 | 0.97 | 0.64 |
| Harmony (regress CC) | 0.52 | 0.63 | 0.61 | 0.55 | 0.55 | 0.53 | 0.94 | 0.62 |
| Harmony (batch only) | 0.52 | 0.60 | 0.58 | 0.50 | 0.49 | 0.53 | 0.94 | 0.59 |

Figure S18: **Time-series dataset integration and benchmarking. (A)** Schematic of the integration process. Shared low-dimensional representations were generated using three integration tools (Harmony[4], scVI[5], scANVI[6]) with and without cell cycle correction. **(B)** Batch correction benchmarking results performed on non-reprogrammed cells from each of the three datasets. **(C)** Biological conservation benchmarking results performed on cells from the time-series dataset. **(D)** Force-directed graph layouts generated from the unintegrated PCA representation, colored by dataset, cell cycle phase, and non-reprogrammed cells (top to bottom). **(E)** Force-directed graph layouts generated from the scANVI representation colored by non-reprogrammed cells and cell cycle phase. Left: Batch (dataset) was the only covariate; Right: Cell cycle phase scores were continuous covariates in addition to batch.

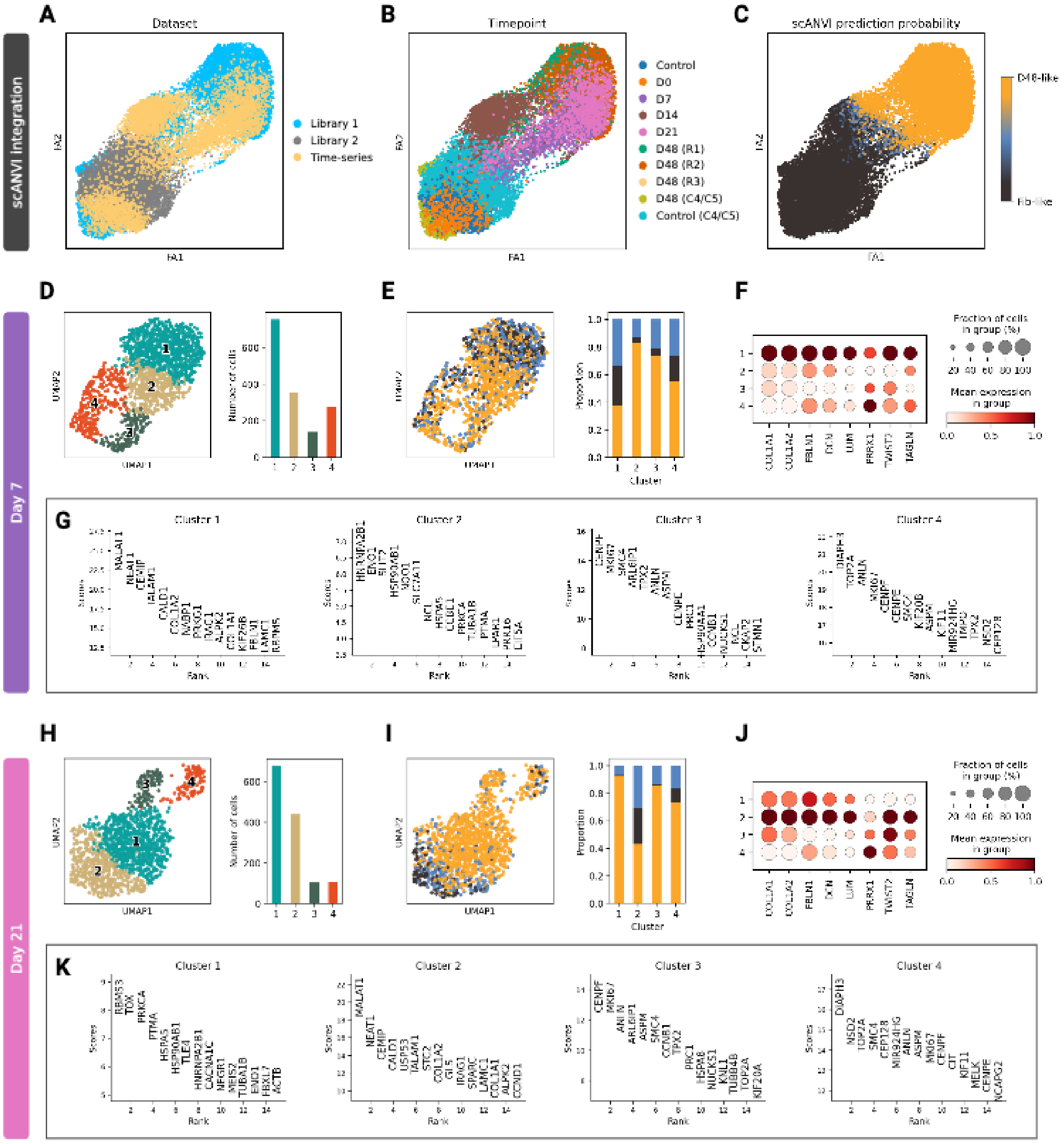


Figure S19: **scANVI predictions and clustering of time-series cells.** **(A)** Force-directed graph layout colored by dataset generated from the scANVI latent representation with cell cycle covariates. **(B)** Graph from (A) colored by timepoint. **(C)** Graph from (A) colored by scANVI prediction probability for fibroblast-like or day 48-like cell classes. **(D)** Left: UMAP plot of day 7 cells colored by Leiden cluster; Right: Number of cells per cluster. **(E)** Left: Day 7 UMAP colored by scANVI-predicted cell class; Right: Distribution of predicted cell classes across clusters. **(F)** Log-normalized expression (scaled from $0-1$) for select fibroblast marker genes across day 7 clusters. **(G)** Top 15 DEGs for each day 7 cluster ranked by z-score (Wilcoxon Rank-sum test, corrected $\alpha = 0.05$). **(H)** Left: UMAP plot of day 21 cells colored by Leiden cluster; Right: Number of cells per cluster. **(I)** Left: Day 21 UMAP colored by scANVI-predicted cell class; Right: Distribution of predicted cell classes across clusters. **(J)** Log-normalized expression (scaled from $0-1$) for select fibroblast marker genes across day 21 clusters. **(K)** Top 15 DEGs for each day 21 cluster ranked by z-score (Wilcoxon Rank-sum test, corrected $\alpha = 0.05$).

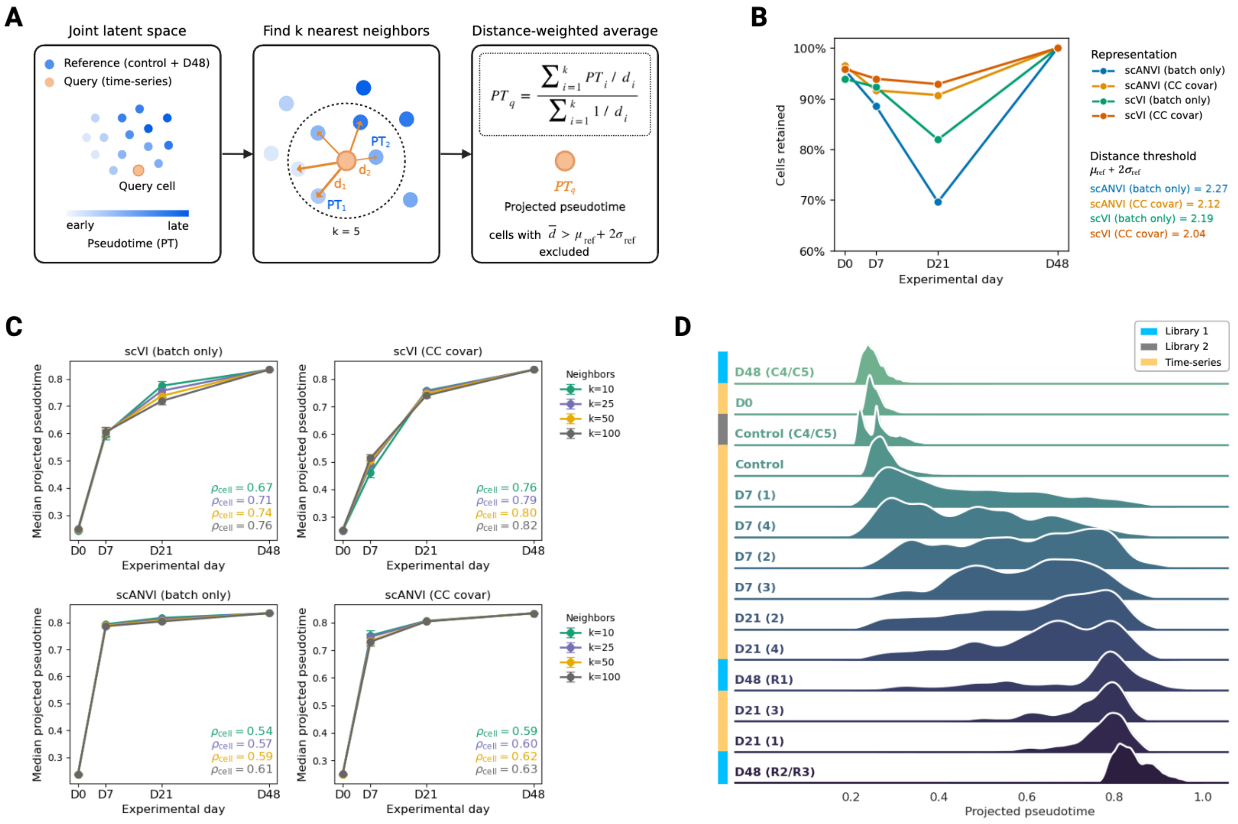


Figure S20: **Pseudotime projection.** **(A)** Schematic of the pseudotime projection method. **(B)** The percentage of cells retained per timepoint after dropping low coverage cells. Lines are colored by latent representation. Query cells (time-series cells) with a mean neighbor distance less than the indicated thresholds were retained. **(C)** Median projected pseudotime values per timepoint for each latent representation. Error bars represent bootstrapped 95% CIs. Lines are colored by the number of nearest neighbors used in the projection. Spearman's correlation ($\rho_{cell}$) was computed on per-cell distributions of projected pseudotime values versus experimental day. **(D)** Density plots of projected pseudotime distributions per cluster-resolved timepoint. The scVI representation with cell cycle covariates was used for projection with $k = 25$ nearest neighbors. Clusters for Libraries 1 and 2 are derived from Fig. 2F. Clusters for time-series cells are derived from Supplementary Fig. S19D and H. Spearman's $\rho$ values were computed on per-cell distributions of projected pseudotime values (with scVI and scANVI representations) versus experimental day across the clusters indicated by the red dots.

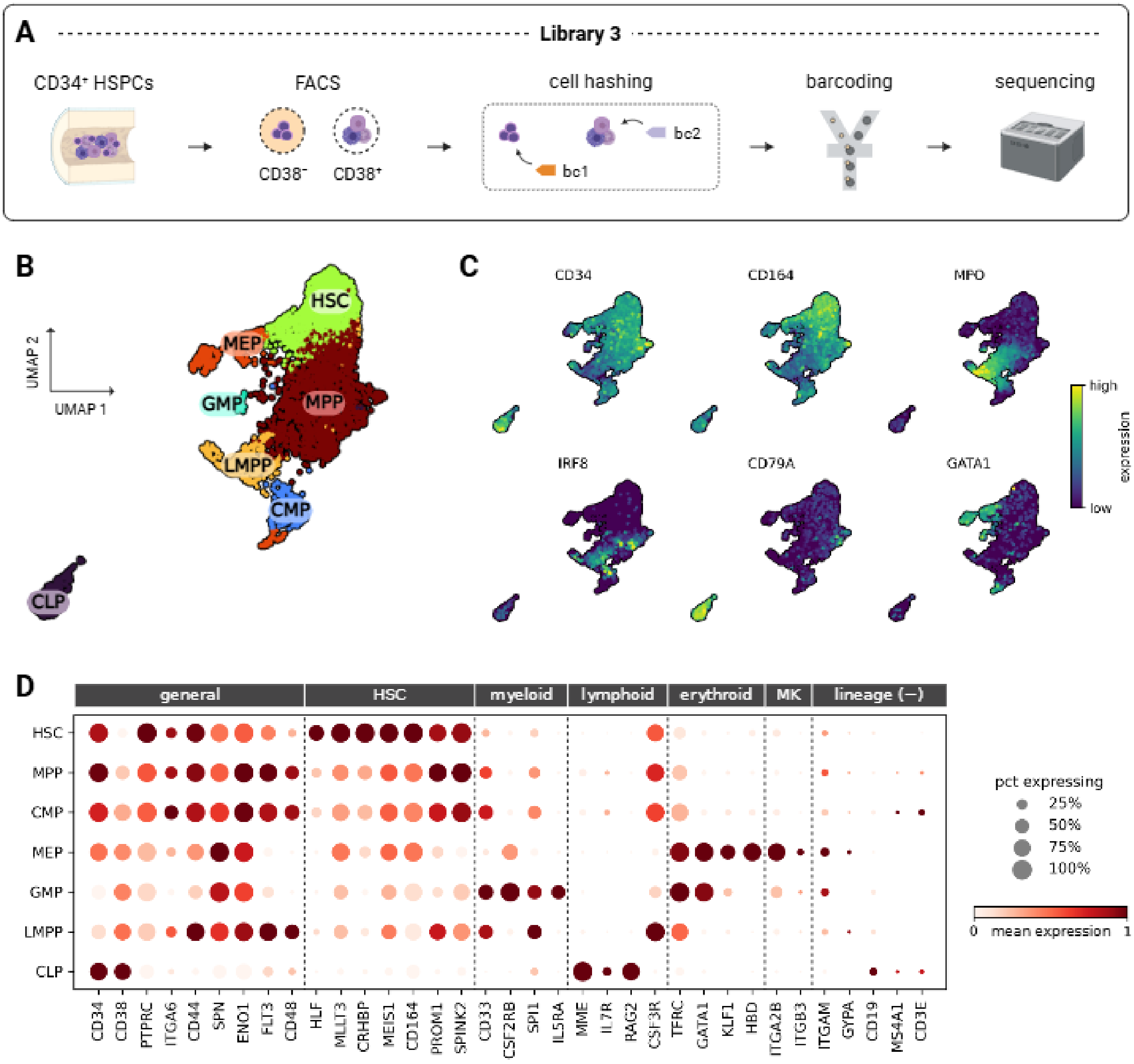


Figure S21: **Characterization of CD34+ human bone marrow scRNA-seq. (A)** Experimental workflow for scRNA-seq of human bone marrow cells (Library 3), including cell hashing of FACS-sorted CD38+ and CD38- populations. **(B)** UMAP of cells colored by Leiden cluster and labeled by cell type annotations. **(C)** Normalized expression of key hematopoietic transcripts[7] on UMAP coordinates from (B). **(D)** Dot plot showing mean log-normalized expression of lineage-specific marker genes per cell type. Dot size represents the percentage of cells expressing a given gene.

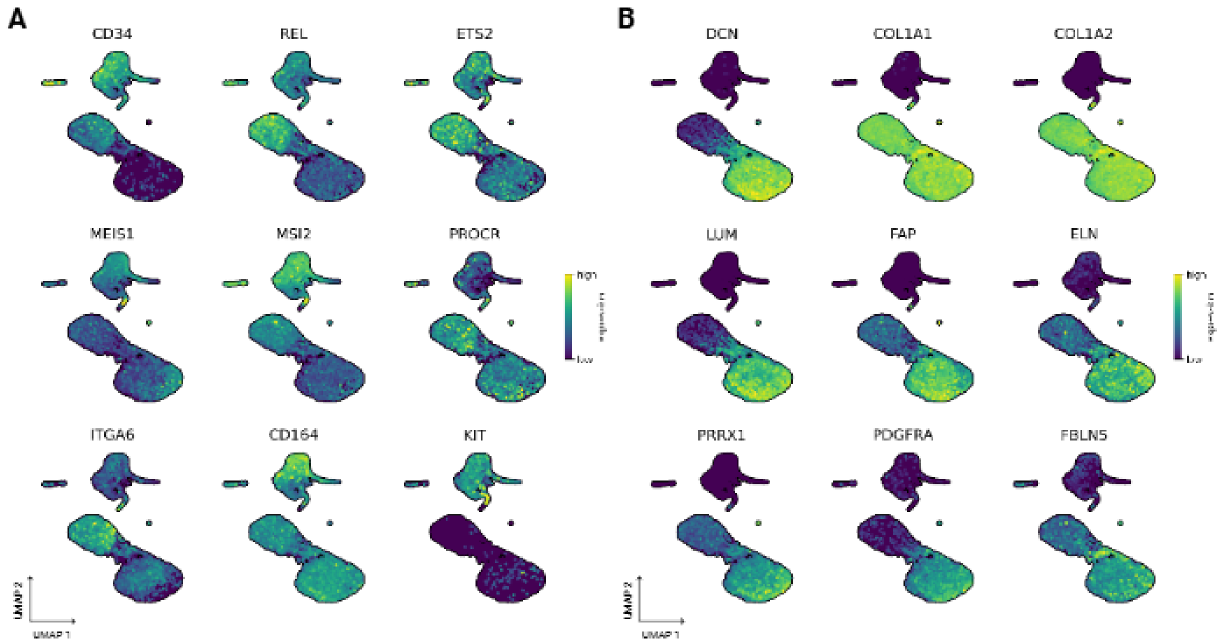


Figure S22: **Marker gene expression in initial, reprogrammed, and target cells.** **(A)** Log-normalized expression of hematopoietic genes and **(B)** fibroblast genes on the UMAP coordinates from Fig. 4B.

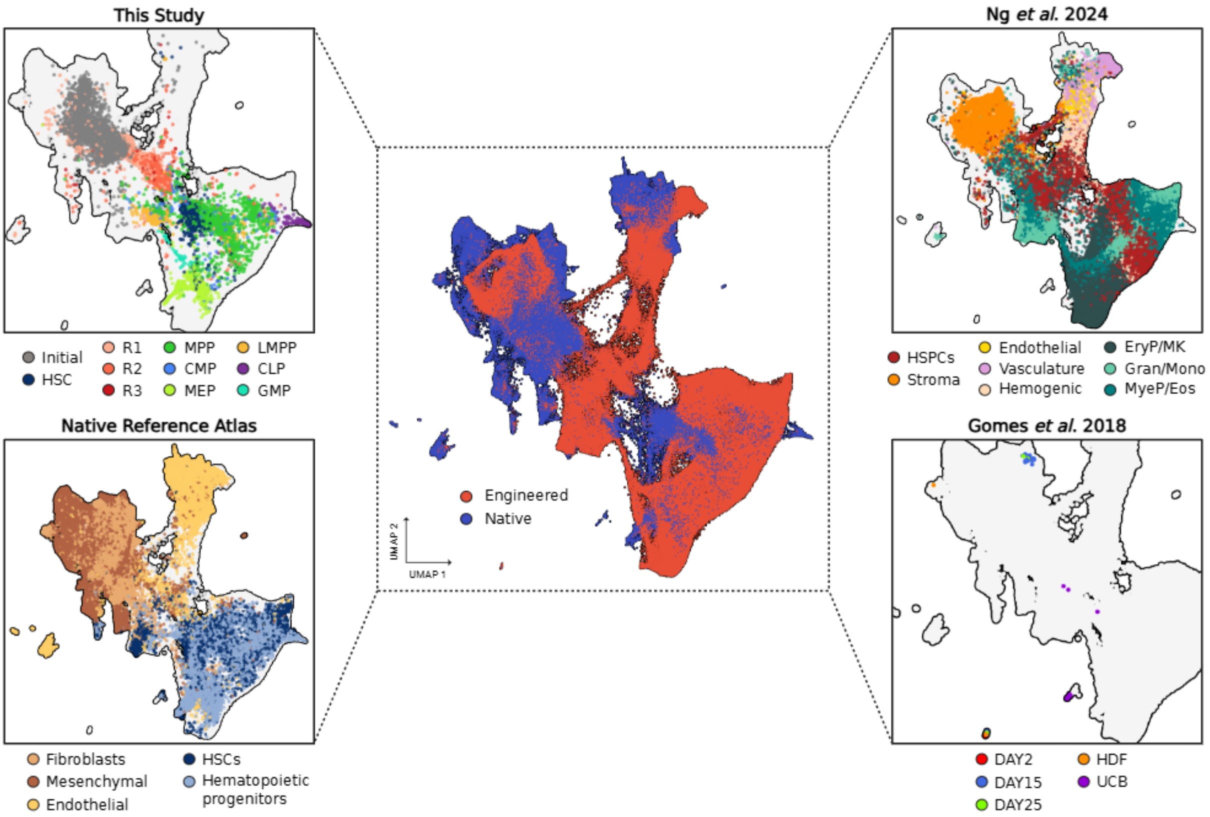


Figure S23: **Overlap between data sources in the integrated atlas.** **Center:** UMAP of 420,108 cells from the native reference atlas, this study, and selected sources based on scVI latent space, colored by engineered or native cell types. **Top left:** Same UMAP showing the 20,620 cells from this study (reprogrammed, control, CD34+ human bone marrow) colored by their cell type annotations. HSC: hematopoietic stem cell, MPP: multipotent progenitor, CMP: common myeloid progenitor, MEP: megakaryocyte-erythroid progenitor, LMPP: lymphoid-primed multipotent progenitor, CLP: common lymphoid progenitor, GMP: granulocyte-monocyte progenitor; **Bottom left:** Same UMAP showing 167,133 cells from the native reference atlas colored by cell annotation; **Top right:** Same UMAP showing 232,072 engineered cells from[8]. These cells underwent in vitro differentiation into the hematopoietic lineage from induced pluripotent stem cells, and annotations from the source were consolidated into broader annotations. HSPCs: hematopoietic stem and progenitor cells, EryP/MK: erythroid-megakaryocyte progenitor/megakaryocyte, Gran/Mono: granulocyte/monocyte, MyeP/Eos: myeloid progenitor/eosinophil; **Bottom right:** Same UMAP showing 283 control and reprogrammed cells from[9]. Reprogrammed cells were induced with the 3TF recipe (GATA2, GFI1B, FOS). HDF: human dermal fibroblasts, UCB: umbilical cord blood (Lin-CD34+), DAY2: 3TF-induced day 2, DAY15: 3TF-induced CD49f+ day 15, DAY25: 3TF-induced CD49f+ day 25.

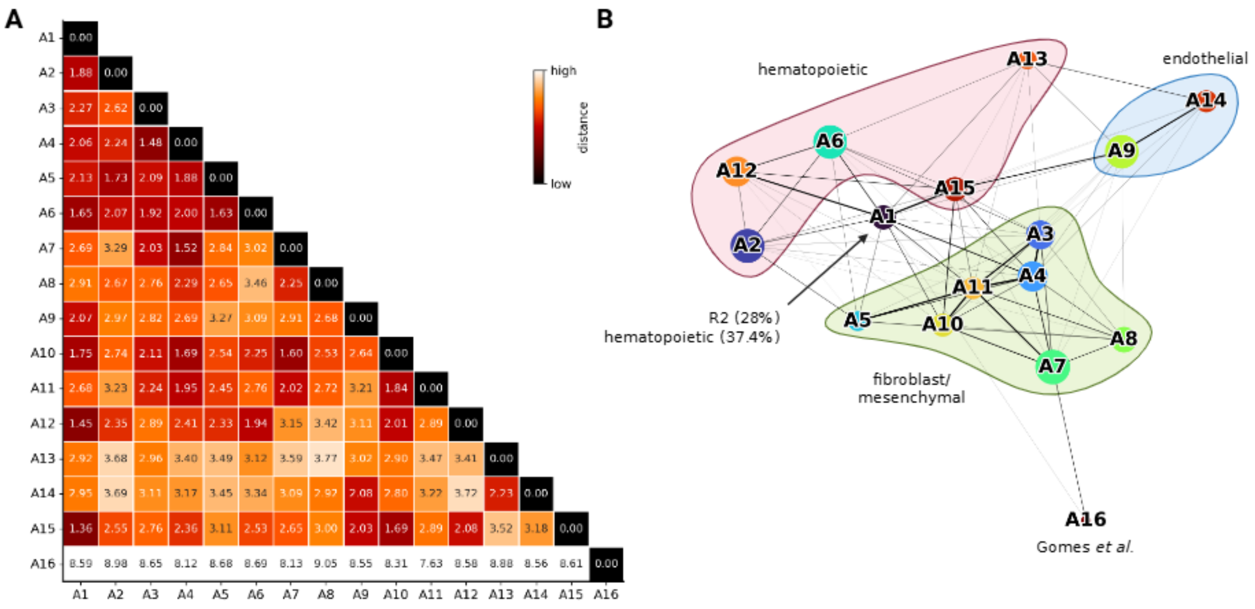


Figure S24: **Pairwise Euclidean distances between clusters in the integrated atlas. (A)** Euclidean distance between mean scVI latent vectors for clusters in the integrated atlas dataset. **(B)** PAGA plot[10] of clusters in the integrated atlas with broad annotations based on most common cell type composition (Table S5).

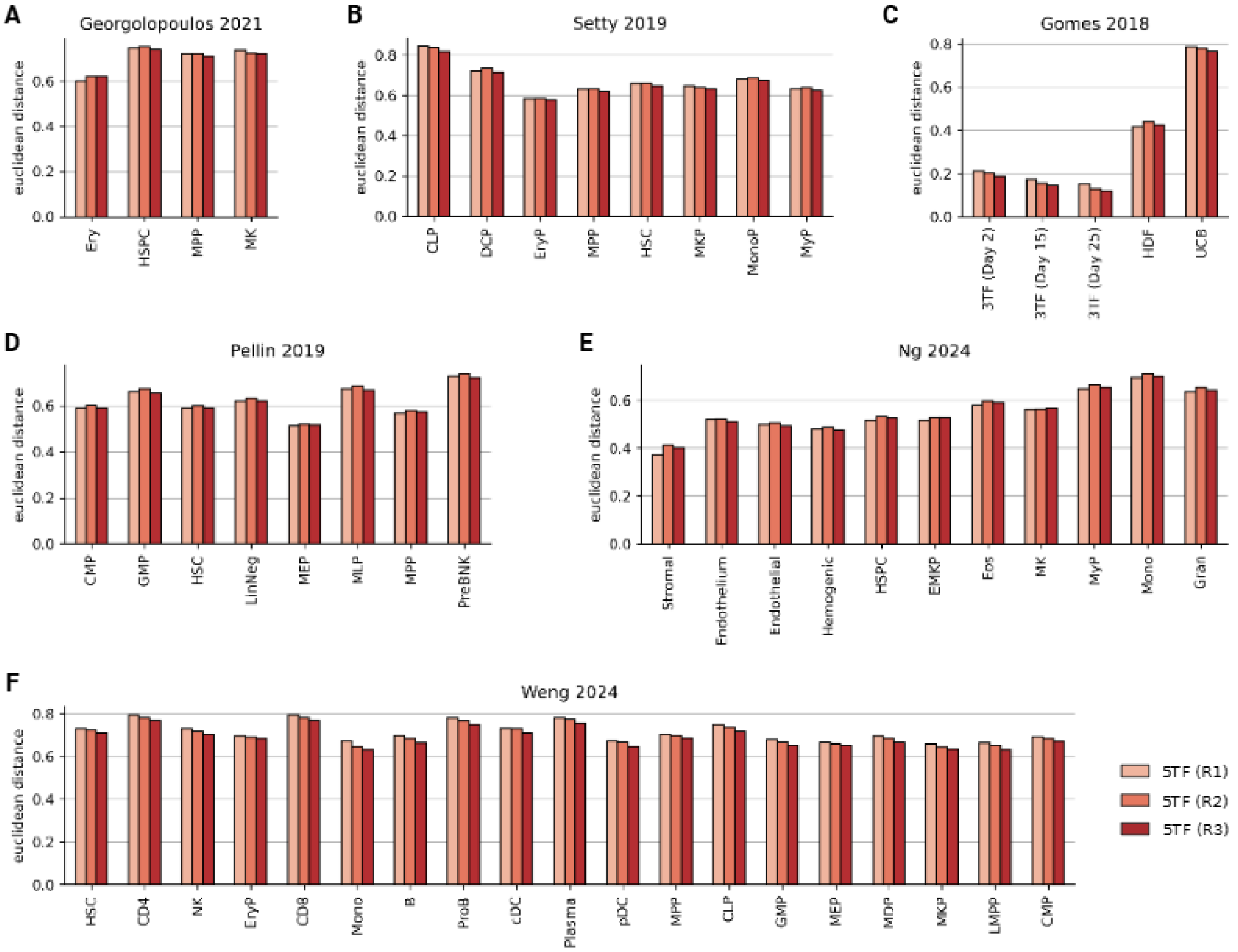


Figure S25: **Pseudobulk distances between reprogrammed cells and datasets from literature. (A)** Distances to cell types in[11]. **(B)** Distances to cell types in[7]. **(C)** Distances to cell types in[9]. **(D)** Distances to cell types in[12]. **(E)** Distances to cell types in[8]. **(F)** Distances to cell types in[13]. Bars indicate distances (min-max normalized) computed independently for each reprogramming cluster (R1–R3).

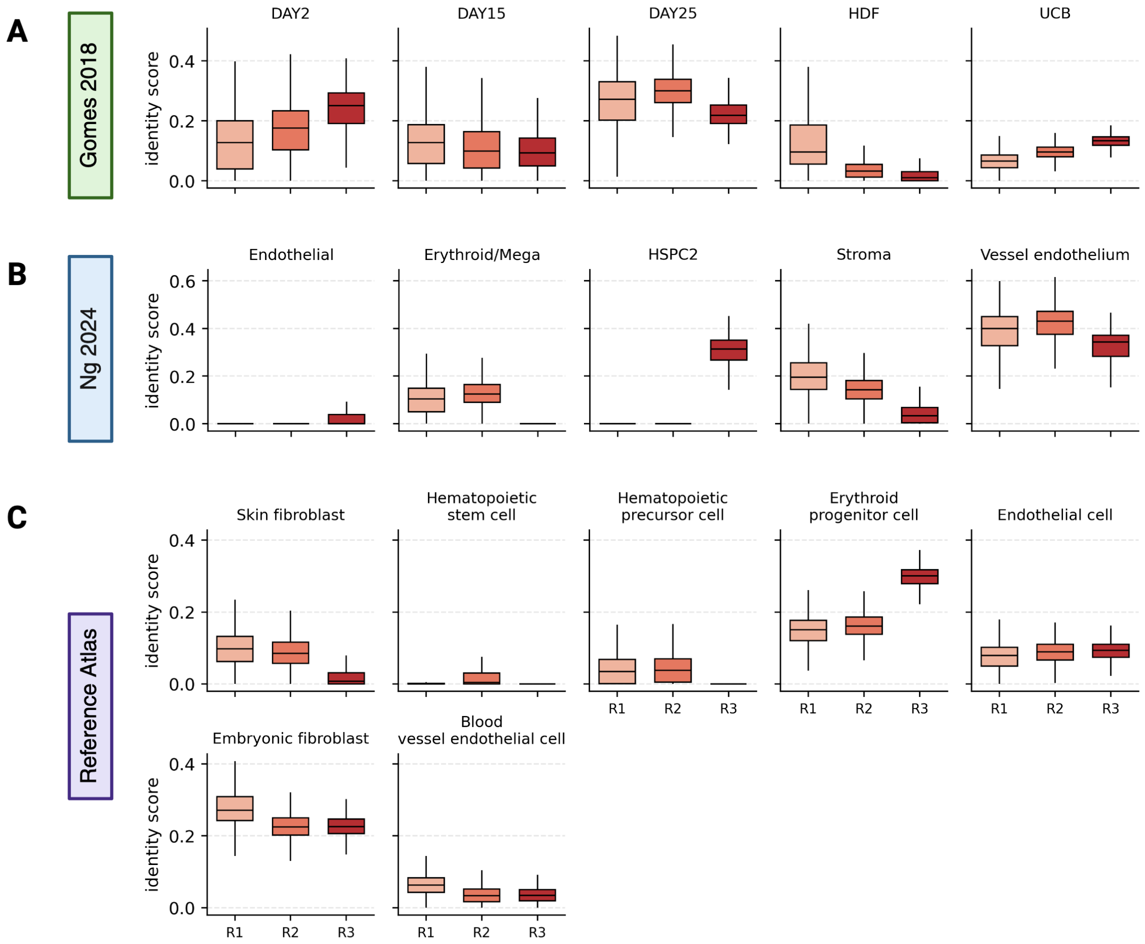


Figure S26: **Capybara identity scores on cell subtypes. (A)** Capybara identity scores of reprogrammed clusters to cell types in[9]. **(B)** Capybara identity scores of reprogrammed clusters to selected cell types in[8]. **(C)** Capybara identity scores of reprogrammed clusters to selected cell types in our native reference atlas. All data are represented as median and IQR, with whiskers extending to $1.5 \times$ IQR.

#### 1.2 Supplementary Tables

| Cell Type | Gene | logFC | Pct in Group | Pct in Ref |
|---|---|---|---|---|
| vascular endothelial cell | TM4SF1 | 5.77 | 0.90 | 0.18 |
| vascular endothelial cell | MT2A | 4.17 | 0.91 | 0.44 |
| vascular endothelial cell | CD59 | 3.56 | 0.84 | 0.34 |
| vascular endothelial cell | SOD2 | 3.95 | 0.83 | 0.35 |
| vascular endothelial cell | UBC | 2.14 | 0.94 | 0.74 |
| vascular endothelial cell | EIF1 | 1.73 | 0.96 | 0.80 |
| vascular endothelial cell | NFKBIA | 2.83 | 0.88 | 0.53 |
| vascular endothelial cell | IFI27 | 3.43 | 0.75 | 0.16 |
| vascular endothelial cell | JUND | 2.67 | 0.88 | 0.49 |
| vascular endothelial cell | ADAMTS9 | 3.87 | 0.69 | 0.13 |
| cord blood HSC | SNHG29 | 4.41 | 0.71 | 0.22 |
| cord blood HSC | HNRNPA1 | 2.48 | 0.82 | 0.69 |
| cord blood HSC | LDHB | 3.34 | 0.76 | 0.51 |
| cord blood HSC | EEF1B2 | 2.45 | 0.81 | 0.64 |
| cord blood HSC | GAS5 | 4.20 | 0.68 | 0.22 |
| cord blood HSC | BTF3 | 1.96 | 0.78 | 0.71 |
| cord blood HSC | NPM1 | 1.98 | 0.77 | 0.69 |
| cord blood HSC | AIF1 | 3.62 | 0.62 | 0.23 |
| cord blood HSC | RPSA | 1.53 | 0.78 | 0.73 |
| cord blood HSC | RPL4 | 1.49 | 0.75 | 0.72 |
| embryonic fibroblast | FN1 | 5.05 | 0.97 | 0.24 |
| embryonic fibroblast | GPC3 | 5.55 | 0.95 | 0.14 |
| embryonic fibroblast | COL3A1 | 5.71 | 0.97 | 0.24 |
| embryonic fibroblast | EGFL6 | 7.62 | 0.87 | 0.03 |
| embryonic fibroblast | DLK1 | 6.83 | 0.89 | 0.09 |
| embryonic fibroblast | PLAGL1 | 4.71 | 0.91 | 0.20 |
| embryonic fibroblast | SERPINE2 | 5.93 | 0.88 | 0.15 |
| embryonic fibroblast | HGF | 5.90 | 0.87 | 0.07 |
| embryonic fibroblast | COL6A2 | 4.30 | 0.94 | 0.31 |
| embryonic fibroblast | CDH11 | 4.71 | 0.89 | 0.14 |
| endothelial cell | EMCN | 3.94 | 0.63 | 0.10 |
| endothelial cell | EGFL7 | 3.34 | 0.68 | 0.18 |
| endothelial cell | EPAS1 | 3.12 | 0.68 | 0.21 |
| endothelial cell | LDB2 | 3.51 | 0.65 | 0.16 |
| endothelial cell | A2M | 3.04 | 0.69 | 0.20 |
| endothelial cell | FLT1 | 3.72 | 0.61 | 0.10 |
| endothelial cell | PECAM1 | 3.20 | 0.64 | 0.19 |
| endothelial cell | PTPRB | 4.02 | 0.57 | 0.07 |
| endothelial cell | ADGRL4 | 3.65 | 0.54 | 0.08 |
| endothelial cell | TM4SF1 | 2.80 | 0.64 | 0.18 |
| erythroid progenitor | H2AZ1 | 3.27 | 0.88 | 0.51 |
| erythroid progenitor | HMGB1 | 2.58 | 0.88 | 0.74 |
| erythroid progenitor | NPM1 | 2.60 | 0.89 | 0.69 |

Continued on next page

Table S1: **Top 10 DEGs in fibroblast, endothelial, and hematopoietic populations.**

| Cell Type | Gene | logFC | Pct in Group | Pct in Ref |
|---|---|---|---|---|
| erythroid progenitor | STMN1 | 3.63 | 0.83 | 0.34 |
| erythroid progenitor | ATP5IF1 | 3.54 | 0.81 | 0.38 |
| erythroid progenitor | PRDX2 | 3.82 | 0.80 | 0.38 |
| erythroid progenitor | BLVRB | 4.16 | 0.76 | 0.26 |
| erythroid progenitor | HINT1 | 2.43 | 0.85 | 0.64 |
| erythroid progenitor | TUBA1B | 2.89 | 0.88 | 0.59 |
| erythroid progenitor | TYMS | 4.43 | 0.65 | 0.09 |
| fibroblast | DCN | 4.44 | 0.85 | 0.23 |
| fibroblast | COL6A2 | 3.05 | 0.79 | 0.25 |
| fibroblast | COL1A2 | 3.55 | 0.77 | 0.22 |
| fibroblast | COL6A1 | 2.98 | 0.72 | 0.20 |
| fibroblast | LUM | 3.88 | 0.66 | 0.13 |
| fibroblast | SERPINF1 | 3.65 | 0.63 | 0.12 |
| fibroblast | COL6A3 | 3.12 | 0.65 | 0.15 |
| fibroblast | FBLN1 | 3.16 | 0.65 | 0.15 |
| fibroblast | MEG3 | 2.94 | 0.67 | 0.21 |
| fibroblast | FSTL1 | 2.74 | 0.68 | 0.21 |
| multipotent progenitor (MPP) | SPINK2 | 6.52 | 0.92 | 0.04 |
| multipotent progenitor (MPP) | HINT1 | 2.95 | 0.99 | 0.64 |
| multipotent progenitor (MPP) | RACK1 | 2.85 | 1.00 | 0.73 |
| multipotent progenitor (MPP) | AIF1 | 3.98 | 0.98 | 0.22 |
| multipotent progenitor (MPP) | NPM1 | 2.75 | 1.00 | 0.69 |
| multipotent progenitor (MPP) | RPLP1 | 2.52 | 1.00 | 0.83 |
| multipotent progenitor (MPP) | HSP90AB1 | 2.69 | 1.00 | 0.70 |
| multipotent progenitor (MPP) | MACROH2A1 | 3.01 | 0.93 | 0.37 |
| multipotent progenitor (MPP) | ENO1 | 2.55 | 0.97 | 0.54 |
| multipotent progenitor (MPP) | HOPX | 4.01 | 0.79 | 0.09 |
| hematopoietic precursor | MACROH2A1 | 2.50 | 0.88 | 0.37 |
| hematopoietic precursor | RPS17 | 3.31 | 0.85 | 0.47 |
| hematopoietic precursor | PRSS57 | 4.23 | 0.68 | 0.06 |
| hematopoietic precursor | HNRNPA1 | 2.13 | 0.98 | 0.69 |
| hematopoietic precursor | SEPTIN6 | 2.59 | 0.83 | 0.24 |
| hematopoietic precursor | SERPINB1 | 2.70 | 0.82 | 0.29 |
| hematopoietic precursor | GIHCG | 3.64 | 0.66 | 0.10 |
| hematopoietic precursor | RPS3 | 1.62 | 0.95 | 0.79 |
| hematopoietic precursor | RPL3 | 1.72 | 0.98 | 0.78 |
| hematopoietic precursor | RPS4X | 1.65 | 0.95 | 0.78 |
| hematopoietic stem cell (HSC) | SPINK2 | 4.88 | 0.74 | 0.05 |
| hematopoietic stem cell (HSC) | SMIM24 | 4.90 | 0.70 | 0.04 |
| hematopoietic stem cell (HSC) | PRSS57 | 4.50 | 0.71 | 0.05 |
| hematopoietic stem cell (HSC) | RPS3 | 1.85 | 0.89 | 0.79 |
| hematopoietic stem cell (HSC) | CYTL1 | 4.06 | 0.69 | 0.07 |
| hematopoietic stem cell (HSC) | HMGA1 | 3.04 | 0.79 | 0.26 |
| hematopoietic stem cell (HSC) | GIHCG | 3.54 | 0.70 | 0.10 |

Continued on next page

Table S1: **Top 10 DEGs in fibroblast, endothelial, and hematopoietic populations.**

| Cell Type | Gene | logFC | Pct in Group | Pct in Ref |
|---|---|---|---|---|
| hematopoietic stem cell (HSC) | EEF1B2 | 2.07 | 0.85 | 0.64 |
| hematopoietic stem cell (HSC) | RPS24 | 1.48 | 0.86 | 0.78 |
| hematopoietic stem cell (HSC) | RPS23 | 1.54 | 0.87 | 0.78 |
| megakaryocyte progenitor | H3-3A | 4.87 | 1.00 | 0.67 |
| megakaryocyte progenitor | PTMA | 3.66 | 1.00 | 0.82 |
| megakaryocyte progenitor | TAGLN2 | 4.89 | 1.00 | 0.55 |
| megakaryocyte progenitor | PLEK | 7.53 | 0.95 | 0.15 |
| megakaryocyte progenitor | CD164 | 5.48 | 0.98 | 0.44 |
| megakaryocyte progenitor | CDK6 | 5.66 | 0.95 | 0.23 |
| megakaryocyte progenitor | HMGB1 | 3.90 | 1.00 | 0.74 |
| megakaryocyte progenitor | YBX1 | 3.89 | 0.99 | 0.69 |
| megakaryocyte progenitor | ITGA2B | 9.24 | 0.91 | 0.03 |
| megakaryocyte progenitor | TALDO1 | 5.13 | 0.98 | 0.40 |
| dermal fibroblast | DCN | 5.84 | 0.97 | 0.29 |
| dermal fibroblast | CXCL14 | 6.22 | 0.82 | 0.08 |
| dermal fibroblast | COL6A2 | 4.05 | 0.90 | 0.30 |
| dermal fibroblast | S100A6 | 3.50 | 0.98 | 0.64 |
| dermal fibroblast | COL1A2 | 4.54 | 0.90 | 0.26 |
| dermal fibroblast | C1S | 4.25 | 0.83 | 0.18 |
| dermal fibroblast | CTSK | 4.89 | 0.78 | 0.11 |
| dermal fibroblast | GSN | 4.05 | 0.90 | 0.40 |
| dermal fibroblast | SOD3 | 4.83 | 0.78 | 0.11 |
| dermal fibroblast | COL6A1 | 3.92 | 0.84 | 0.24 |
| venous endothelial cell | TM4SF1 | 5.34 | 0.90 | 0.19 |
| venous endothelial cell | CD59 | 3.34 | 0.87 | 0.34 |
| venous endothelial cell | HLA-E | 2.43 | 0.94 | 0.61 |
| venous endothelial cell | PLVAP | 4.38 | 0.69 | 0.07 |
| venous endothelial cell | IFI27 | 3.88 | 0.74 | 0.16 |
| venous endothelial cell | GNG11 | 3.01 | 0.80 | 0.26 |
| venous endothelial cell | IGFBP7 | 2.98 | 0.90 | 0.40 |
| venous endothelial cell | ACKR1 | 6.27 | 0.62 | 0.04 |
| venous endothelial cell | SPARCL1 | 3.34 | 0.81 | 0.32 |
| venous endothelial cell | CAV1 | 3.06 | 0.79 | 0.25 |

Table S1: **Top 10 DEGs in fibroblast, endothelial, and hematopoietic populations.**

| Cell Type | Gene | logFC | Pct in Group | Pct in Ref |
|---|---|---|---|---|
| vascular endothelial cell | CD59 | 3.56 | 0.84 | 0.34 |
| vascular endothelial cell | JUND | 2.67 | 0.88 | 0.49 |
| vascular endothelial cell | ETS2 | 2.78 | 0.71 | 0.27 |
| vascular endothelial cell | MAFF | 2.91 | 0.65 | 0.19 |
| vascular endothelial cell | TCF4 | 2.07 | 0.79 | 0.40 |

Continued on next page

Table S2: **Top 10 differentially expressed TFs in fibroblast, endothelial, and hematopoietic populations.**

| Cell Type | Gene | logFC | Pct in Group | Pct in Ref |
|---|---|---|---|---|
| vascular endothelial cell | ATF3 | 2.70 | 0.62 | 0.21 |
| vascular endothelial cell | KLF6 | 1.87 | 0.80 | 0.53 |
| vascular endothelial cell | ETS1 | 2.10 | 0.64 | 0.27 |
| vascular endothelial cell | SOX17 | 4.13 | 0.45 | 0.04 |
| vascular endothelial cell | YWHAE | 1.46 | 0.76 | 0.49 |
| cord blood HSC | HNRNPA1 | 2.48 | 0.82 | 0.69 |
| cord blood HSC | UBB | 1.67 | 0.69 | 0.68 |
| cord blood HSC | ENO1 | 1.98 | 0.58 | 0.55 |
| cord blood HSC | RPS10 | 1.36 | 0.55 | 0.58 |
| cord blood HSC | HMGA1 | 2.14 | 0.37 | 0.26 |
| cord blood HSC | RAN | 1.37 | 0.47 | 0.54 |
| cord blood HSC | SOX4 | 1.13 | 0.31 | 0.34 |
| cord blood HSC | NME1 | 1.74 | 0.22 | 0.21 |
| cord blood HSC | TMSB4XP8 | 2.53 | 0.02 | 0.01 |
| cord blood HSC | XRCC1 | 1.30 | 0.07 | 0.10 |
| embryonic fibroblast | PLAGL1 | 4.71 | 0.91 | 0.20 |
| embryonic fibroblast | PITX2 | 7.80 | 0.79 | 0.01 |
| embryonic fibroblast | TCF21 | 3.53 | 0.62 | 0.08 |
| embryonic fibroblast | PEG3 | 4.66 | 0.54 | 0.05 |
| embryonic fibroblast | PDLIM5 | 2.85 | 0.69 | 0.23 |
| embryonic fibroblast | HAND2 | 4.95 | 0.51 | 0.03 |
| embryonic fibroblast | LIN28B | 5.97 | 0.49 | 0.02 |
| embryonic fibroblast | EZR | 2.14 | 0.71 | 0.28 |
| embryonic fibroblast | EPAS1 | 2.03 | 0.72 | 0.22 |
| embryonic fibroblast | CREB3L1 | 3.64 | 0.53 | 0.06 |
| endothelial cell | EPAS1 | 3.12 | 0.68 | 0.21 |
| endothelial cell | ERG | 3.26 | 0.47 | 0.09 |
| endothelial cell | TCF4 | 1.82 | 0.69 | 0.40 |
| endothelial cell | NFIB | 1.78 | 0.57 | 0.27 |
| endothelial cell | MECOM | 3.29 | 0.39 | 0.09 |
| endothelial cell | NPDC1 | 2.36 | 0.44 | 0.15 |
| endothelial cell | MEF2C | 1.66 | 0.56 | 0.27 |
| endothelial cell | ID1 | 2.01 | 0.50 | 0.22 |
| endothelial cell | ELK3 | 2.03 | 0.46 | 0.18 |
| endothelial cell | ETS2 | 1.79 | 0.51 | 0.27 |
| erythroid progenitor | HMGB1 | 2.58 | 0.88 | 0.74 |
| erythroid progenitor | NFE2 | 4.40 | 0.55 | 0.06 |
| erythroid progenitor | HMGA1 | 2.85 | 0.60 | 0.26 |
| erythroid progenitor | RAN | 1.89 | 0.69 | 0.53 |
| erythroid progenitor | LMO2 | 3.44 | 0.52 | 0.14 |
| erythroid progenitor | NME1 | 2.89 | 0.55 | 0.20 |
| erythroid progenitor | KLF1 | 4.66 | 0.42 | 0.03 |
| erythroid progenitor | GATA1 | 4.67 | 0.41 | 0.03 |
| erythroid progenitor | ENO1 | 1.55 | 0.71 | 0.54 |
| erythroid progenitor | HMGB2 | 1.96 | 0.57 | 0.35 |

Continued on next page

Table S2: **Top 10 differentially expressed TFs in fibroblast, endothelial, and hematopoietic populations.**

| Cell Type | Gene | logFC | Pct in Group | Pct in Ref |
|---|---|---|---|---|
| fibroblast | PRRX1 | 2.93 | 0.51 | 0.10 |
| fibroblast | ARID5B | 1.88 | 0.62 | 0.31 |
| fibroblast | NFIA | 1.64 | 0.61 | 0.29 |
| fibroblast | EGR1 | 1.78 | 0.55 | 0.27 |
| fibroblast | ZBTB20 | 1.40 | 0.59 | 0.34 |
| fibroblast | TWIST2 | 2.61 | 0.32 | 0.06 |
| fibroblast | CEBPD | 1.39 | 0.57 | 0.33 |
| fibroblast | TCF4 | 1.10 | 0.63 | 0.39 |
| fibroblast | PLAGL1 | 1.79 | 0.41 | 0.18 |
| fibroblast | RBFOX2 | 1.41 | 0.42 | 0.17 |
| multipotent progenitor (MPP) | ENO1 | 2.55 | 0.97 | 0.54 |
| multipotent progenitor (MPP) | HMGB1 | 2.10 | 0.99 | 0.74 |
| multipotent progenitor (MPP) | MSI2 | 2.95 | 0.79 | 0.20 |
| multipotent progenitor (MPP) | HMGA1 | 2.89 | 0.79 | 0.25 |
| multipotent progenitor (MPP) | LMO2 | 3.35 | 0.72 | 0.13 |
| multipotent progenitor (MPP) | YBX1 | 1.83 | 0.98 | 0.68 |
| multipotent progenitor (MPP) | HHEX | 3.25 | 0.64 | 0.10 |
| multipotent progenitor (MPP) | BCL11A | 3.27 | 0.61 | 0.08 |
| multipotent progenitor (MPP) | SOX4 | 2.04 | 0.85 | 0.33 |
| multipotent progenitor (MPP) | APEX1 | 1.92 | 0.76 | 0.34 |
| hematopoietic precursor | HNRNPA1 | 2.13 | 0.98 | 0.69 |
| hematopoietic precursor | RPS4X | 1.65 | 0.95 | 0.78 |
| hematopoietic precursor | HMGA1 | 2.29 | 0.80 | 0.26 |
| hematopoietic precursor | SOX4 | 2.09 | 0.85 | 0.34 |
| hematopoietic precursor | RPL35 | 1.18 | 0.87 | 0.76 |
| hematopoietic precursor | APEX1 | 1.74 | 0.75 | 0.34 |
| hematopoietic precursor | NAP1L1 | 1.41 | 0.92 | 0.61 |
| hematopoietic precursor | ENO1 | 1.56 | 0.88 | 0.55 |
| hematopoietic precursor | LYL1 | 2.68 | 0.52 | 0.10 |
| hematopoietic precursor | RPL6 | 1.21 | 0.93 | 0.76 |
| hematopoietic stem cell (HSC) | HMGA1 | 3.04 | 0.79 | 0.26 |
| hematopoietic stem cell (HSC) | RPS4X | 1.58 | 0.87 | 0.78 |
| hematopoietic stem cell (HSC) | APEX1 | 2.14 | 0.79 | 0.34 |
| hematopoietic stem cell (HSC) | HNRNPA1 | 1.79 | 0.87 | 0.69 |
| hematopoietic stem cell (HSC) | PARP1 | 2.07 | 0.71 | 0.23 |
| hematopoietic stem cell (HSC) | ENO1 | 1.73 | 0.83 | 0.54 |
| hematopoietic stem cell (HSC) | BCL11A | 2.94 | 0.55 | 0.09 |
| hematopoietic stem cell (HSC) | MYB | 3.62 | 0.50 | 0.05 |
| hematopoietic stem cell (HSC) | MSI2 | 2.18 | 0.67 | 0.20 |
| hematopoietic stem cell (HSC) | NFE2 | 2.52 | 0.50 | 0.07 |
| megakaryocyte progenitor | TAGLN2 | 4.89 | 1.00 | 0.55 |
| megakaryocyte progenitor | HMGB1 | 3.90 | 1.00 | 0.74 |
| megakaryocyte progenitor | YBX1 | 3.89 | 0.99 | 0.69 |
| megakaryocyte progenitor | LMO2 | 6.06 | 0.94 | 0.15 |
| megakaryocyte progenitor | ENO1 | 4.09 | 0.99 | 0.55 |

Continued on next page

Table S2: **Top 10 differentially expressed TFs in fibroblast, endothelial, and hematopoietic populations.**

| Cell Type | Gene | logFC | Pct in Group | Pct in Ref |
|---|---|---|---|---|
| megakaryocyte progenitor | NFE2 | 6.15 | 0.82 | 0.07 |
| megakaryocyte progenitor | TAL1 | 5.94 | 0.79 | 0.06 |
| megakaryocyte progenitor | XBP1 | 3.44 | 0.89 | 0.39 |
| megakaryocyte progenitor | ETV6 | 3.70 | 0.82 | 0.22 |
| megakaryocyte progenitor | MSI2 | 3.37 | 0.72 | 0.21 |
| dermal fibroblast | ANXA1 | 2.30 | 0.81 | 0.41 |
| dermal fibroblast | PRRX1 | 2.84 | 0.56 | 0.14 |
| dermal fibroblast | TWIST1 | 3.23 | 0.40 | 0.07 |
| dermal fibroblast | PRNP | 1.93 | 0.53 | 0.25 |
| dermal fibroblast | PRRX2 | 3.80 | 0.35 | 0.04 |
| dermal fibroblast | OSR2 | 3.44 | 0.36 | 0.05 |
| dermal fibroblast | NFIX | 2.51 | 0.40 | 0.12 |
| dermal fibroblast | RBM3 | 1.16 | 0.69 | 0.53 |
| dermal fibroblast | XG | 5.06 | 0.31 | 0.01 |
| dermal fibroblast | KLF4 | 1.69 | 0.49 | 0.24 |
| venous endothelial cell | CD59 | 3.34 | 0.87 | 0.34 |
| venous endothelial cell | ETS2 | 2.66 | 0.73 | 0.27 |
| venous endothelial cell | NPDC1 | 3.11 | 0.64 | 0.15 |
| venous endothelial cell | ZNF385D | 3.13 | 0.57 | 0.10 |
| venous endothelial cell | EPAS1 | 2.44 | 0.66 | 0.22 |
| venous endothelial cell | TCF4 | 1.85 | 0.79 | 0.40 |
| venous endothelial cell | ELK3 | 2.28 | 0.57 | 0.19 |
| venous endothelial cell | ID1 | 2.43 | 0.59 | 0.22 |
| venous endothelial cell | SOX7 | 3.62 | 0.42 | 0.05 |
| venous endothelial cell | TAGLN2 | 1.32 | 0.82 | 0.55 |

Table S2: **Top 10 differentially expressed TFs in fibroblast, endothelial, and hematopoietic populations.**

| Gene | $t$ | df | $\bar{x}_A \pm SD$ | $\bar{x}_B \pm SD$ | Mean diff | 95% CI | Effect size | P value |
|---|---|---|---|---|---|---|---|---|
| GATA2 | 25.37 | 2.46 | -9.87 ± 0.76 | 2.42 ± 0.26 | 11.69 | 10.02–13.35 | $\eta^2 = 0.996$ | 0.0005 |
| GFI1B | 9.98 | 2.39 | -10.64 ± 1.00 | -4.59 ± 0.32 | 6.06 | 3.82–8.29 | $\eta^2 = 0.977$ | 0.0053 |
| FOS | 61.33 | 3.53 | -13.69 ± 0.39 | 3.04 ± 0.27 | 16.73 | 15.93–17.53 | $\eta^2 = 0.999$ | $< 0.0001$ |
| STAT5A | 25.27 | 2.19 | -10.08 ± 0.94 | 4.21 ± 0.21 | 14.28 | 12.07–16.49 | $\eta^2 = 0.997$ | 0.0009 |
| REL | 17.27 | 3.98 | -10.16 ± 0.57 | -1.90 ± 0.61 | 8.26 | 6.93–9.59 | $\eta^2 = 0.987$ | $< 0.0001$ |

Table S3: **Statistical details for qRT-PCR analyses.** Group comparisons were performed using unpaired two-tailed $t$-tests with Welch's correction. Effect size is reported as $\eta^2$ (eta squared). Sample means and standard deviations are reported as $\bar{x} \pm SD$ for control (A) and treatment (B). Mean differences are reported as $B - A$ with $95\%$ confidence intervals.

| **Barcoded Phase** | **Predicted G1** | **Predicted G2M** | **Predicted S** |
|---|---|---|---|
| G1 | 2708 (47.5%) | 1008 (17.7%) | 1151 (20.2%) |
| G2M | 87 (6.9%) | 910 (72.1%) | 246 (19.5%) |
| S | 331 (26.3%) | 478 (38.0%) | 563 (44.8%) |

Table S4: **Confusion matrix of predicted cell cycle phases vs. feature barcodes.** Raw counts and row-normalized percentages are shown.

| Cluster | Annotation | Data Key | Count | Proportion |
|---|---|---|---|---|
| A1 | R2 | This Study | 3924 | 0.281000 |
| A1 | hematopoietic multipotent progenitor cell | Reference | 2736 | 0.196000 |
| A1 | HSC | This Study | 1343 | 0.096000 |
| A1 | hematopoietic stem cell | Reference | 1142 | 0.082000 |
| A1 | R1 | This Study | 1004 | 0.072000 |
| A10 | mesothelial cell | Reference | 4419 | 0.402000 |
| A10 | mesenchymal cell | Reference | 2810 | 0.256000 |
| A10 | hematopoietic multipotent progenitor cell | Reference | 1108 | 0.101000 |
| A10 | fibroblast of lung | Reference | 746 | 0.068000 |
| A10 | pulmonary interstitial fibroblast | Reference | 404 | 0.037000 |
| A11 | fibroblast | Reference | 2843 | 0.312000 |
| A11 | mesenchymal cell | Reference | 1059 | 0.116000 |
| A11 | fibroblast of cardiac tissue | Reference | 979 | 0.107000 |
| A11 | initial | This Study | 704 | 0.077000 |
| A11 | fibro/adipogenic progenitor cell | Reference | 659 | 0.072000 |
| A12 | HSPC1 | Ng 2024 | 12965 | 0.369000 |
| A12 | EMK Pro1 | Ng 2024 | 3502 | 0.100000 |
| A12 | HSPC2 | Ng 2024 | 3394 | 0.097000 |
| A12 | Mye Pro2 | Ng 2024 | 3366 | 0.096000 |
| A12 | EMK Pro2 | Ng 2024 | 3289 | 0.094000 |
| A13 | endothelial cell of lymphatic vessel | Reference | 3237 | 0.734000 |
| A13 | CLP | This Study | 453 | 0.103000 |
| A13 | lymphoid lineage restricted progenitor cell | Reference | 353 | 0.080000 |
| A13 | endothelial cell | Reference | 241 | 0.055000 |
| A13 | early lymphoid progenitor | Reference | 17 | 0.004000 |
| A14 | Venous | Ng 2024 | 7742 | 0.878000 |
| A14 | endothelial cell | Reference | 378 | 0.043000 |
| A14 | capillary endothelial cell | Reference | 71 | 0.008000 |
| A14 | Endo | Ng 2024 | 53 | 0.006000 |
| A14 | Hemogenic | Ng 2024 | 48 | 0.005000 |
| A15 | HSPC1 | Ng 2024 | 4610 | 0.336000 |
| A15 | Hemogenic | Ng 2024 | 4413 | 0.322000 |
| A15 | HSPC2 | Ng 2024 | 1708 | 0.124000 |
| A15 | EMK Pro1 | Ng 2024 | 1061 | 0.077000 |
| A15 | EMK Pro2 | Ng 2024 | 549 | 0.040000 |
| A16 | DAY25 | Gomes 2018 | 91 | 0.453000 |
| A16 | DAY2 | Gomes 2018 | 38 | 0.189000 |
| A16 | HDF | Gomes 2018 | 37 | 0.184000 |
| A16 | DAY15 | Gomes 2018 | 35 | 0.174000 |
| A2 | EMK Pro1 | Ng 2024 | 12141 | 0.206000 |
| A2 | Mega | Ng 2024 | 11511 | 0.195000 |
| A2 | EMK Pro2 | Ng 2024 | 8835 | 0.150000 |
| A2 | Eos | Ng 2024 | 8556 | 0.145000 |
| A2 | Gran | Ng 2024 | 5133 | 0.087000 |

Continued on next page

Table S5: **Top 5 annotation-data key pairs by cluster.**

| Cluster | Annotation | Data Key | Count | Proportion |
|---|---|---|---|---|
| A3 | fibroblast | Reference | 6224 | 0.213000 |
| A3 | Mono | Ng 2024 | 4597 | 0.157000 |
| A3 | Gran | Ng 2024 | 4430 | 0.151000 |
| A3 | Mye Pro1 | Ng 2024 | 4148 | 0.142000 |
| A3 | fibroblast of mammary gland | Reference | 3671 | 0.125000 |
| A4 | mesenchymal cell | Reference | 7741 | 0.218000 |
| A4 | initial | This Study | 6564 | 0.185000 |
| A4 | fibroblast of mammary gland | Reference | 4874 | 0.137000 |
| A4 | fibroblast | Reference | 3204 | 0.090000 |
| A4 | HSPC1 | Ng 2024 | 2792 | 0.079000 |
| A5 | adipocyte | Reference | 4000 | 0.716000 |
| A5 | endothelial tip cell | Reference | 1060 | 0.190000 |
| A5 | fibroblast of mammary gland | Reference | 109 | 0.020000 |
| A5 | capillary endothelial cell | Reference | 104 | 0.019000 |
| A5 | HSPC3 | Ng 2024 | 46 | 0.008000 |
| A6 | Mye Pro1 | Ng 2024 | 13693 | 0.244000 |
| A6 | HSPC1 | Ng 2024 | 12154 | 0.217000 |
| A6 | Mye Pro2 | Ng 2024 | 10104 | 0.180000 |
| A6 | Mono | Ng 2024 | 5162 | 0.092000 |
| A6 | HSPC2 | Ng 2024 | 3704 | 0.066000 |
| A7 | fibroblast | Reference | 13892 | 0.215000 |
| A7 | Strom2 | Ng 2024 | 13569 | 0.210000 |
| A7 | Strom3 | Ng 2024 | 12507 | 0.194000 |
| A7 | mesenchymal cell | Reference | 9202 | 0.143000 |
| A7 | Strom1 | Ng 2024 | 2854 | 0.044000 |
| A8 | pericyte | Reference | 7808 | 0.442000 |
| A8 | vascular associated smooth muscle cell | Reference | 4877 | 0.276000 |
| A8 | fibroblast | Reference | 1937 | 0.110000 |
| A8 | mural cell | Reference | 1420 | 0.080000 |
| A8 | lung pericyte | Reference | 294 | 0.017000 |
| A9 | Art1 | Ng 2024 | 8402 | 0.150000 |
| A9 | endothelial cell | Reference | 7738 | 0.138000 |
| A9 | blood vessel endothelial cell | Reference | 7446 | 0.133000 |
| A9 | Endo | Ng 2024 | 6468 | 0.115000 |
| A9 | vein endothelial cell | Reference | 5100 | 0.091000 |

Table S5: **Top 5 annotation-data key pairs by cluster.**

| Dataset | Cells | Genes | Cell types | Accession | Citation |
|---|---|---|---|---|---|
| Gomes 2018 | 286 | 13,492 | 5 | GSE51025 | Gomes et al.[9] |
| Pellin 2019 | 21,412 | 13,979 | 11 | GSE117498 | Pellin et al.[12] |
| Setty 2019 | 5,780 | 10,438 | 8 | N/A | Setty et al.[7] |
| Georgolopoulos 2021 | 53,511 | 13,384 | 7 | GSE182816 | Georgolopoulos et al.[11] |
| Weng 2024 | 25,252 | 14,179 | 20 | GSE219015 | Weng et al.[13] |
| Ng 2024 | 252,607 | 15,023 | 20 | GSE232710 | Ng et al.[8] |

Table S6: **Summary of Single-Cell Datasets.** Setty 2019 data is available via the Human Cell Atlas portal (https://explore.data.humancellatlas.org/projects/091cf39b-01bc-42e5-9437-f419a66c8a45).

| FH3 | FH4a | FH6 |
|---|---|---|
| GATA2<br>GFI1B<br>FOS | GATA2<br>GFI1B<br>FOS<br>REL<br>STAT5A | GATA2<br>GFI1B<br>FOS<br>REL<br>STAT5A |
| **Vector preparation** | | |
| individual lenti per TF | single lenti | individual lenti per TF |

Table S7: **TF recipes and lentiviral vector preparation methods.**

| Primers | |
|---|---|
| GATA2-V5 Fwd | TTCGAGGAGCTGTCAAAGTG |
| GFI1B-V5 Fwd | CCAGAAGTCCGACATGAAGAAG |
| FOS-V5 Fwd | GGACCTATCTGGGTCCTTCTA |
| STAT5A-V5 Fwd | CAAGTGGTCCCTGAGTTTGT |
| REL-V5 Fwd | TCCATGCCATCAGCAGATTTA |
| Common-V5 Rev | CTAGGAGTGGGTTTGGGATTG |
| LDLR Fwd | ACCGACTCTGTCCTGGGCACTG |
| LDLR Rev | AGTTCCCCAGTCAGTCCAGTA |

Table S8: **Primers used for qRT-PCR validation of 5TF expression.**

| Dataset | Reprogram | Control fibroblasts | Bone marrow HSPCs | Reprogram time-series |
|---|---|---|---|---|
| **Name** | Library 1 | Library 2 | Library 3 | Time-series |
| **Barcoding technology** | 10x Genomics | 10x Genomics | 10x Genomics | Split-pool combinatorial barcoding |
| **Multiplexing method** | N/A | TotalSeqB hashtag antibodies | TotalSeqB hashtag antibodies | Split-pool combinatorial barcoding |
| **Cell populations** | CD34+ D48<br>CD34- D48 | Fibroblasts (G1)<br>Fibroblasts (S)<br>Fibroblasts (G2M) | CD34+/CD38-<br>CD34+/CD38+ | Control fibroblasts; D0, D7, D14, D21 |
| **Total cells** | 8,357 | 6,960 | 5,608 | 6,174 |
| **Mean counts/cell** | 12,143 | 14,571 | 21,013 | 5,416 |
| **Median counts/cell** | 11,091 | 14,160 | 18,366 | 4,511 |
| **Mean genes/cell** | 3,606 | 3,694 | 4,493 | 2,684 |
| **Median genes/cell** | 3,576 | 3,653 | 4,417 | 2,521 |

Table S9: **Overview of scRNA-seq libraries generated in this study.** Total cells and summary statistics are post-quality control.

| | | |
|---|---|---|
| **10x Genomics** | | |
| Next GEM Single Cell 3' Kit (V3.1,V4) | | |
| **Parse Biosciences** | | |
| Parse Cell Fixation Kit v3 | | |
| Evercode WT Mini v3 | | |
| **ONT** | | |
| **Sequencer** | **3' Protocol Version** | **Adapter Kit** |
| GridION | SST_V9148_V111_REVB_12JAN2022 | cDNA-PCR Sequencing Kit (SQK-PCS111) |
| PromethION P2 Solo | SST_V9198_V114_REVB_06DEC2023 | Ligation Sequencing Kit V14 (SQK-LSK114) |
| | **Parse Protocol Version** | **Adapter Kit** |
| PromethION P2 Solo | SST_9198_V114_REVK_29JAN2025 | Ligation Sequencing Kit V14 (SQK-LSK114) |

| | | |
|---|---|---|
| **Primers** | | |
| *Transcript libraries* | | |
| Fwd: 5' - [Biotin] - | CAGCACTTGCCTGTCGCTCTATCTTCCTACACGACGCTCTTCCGATCT | - 3' |
| Rev: 5' - | CAGCTTTCTGTTGGTGCTGATATTGCAAGCAGTGGTATCAACGCAGAG | - 3' |
| *Hashing library* | | |
| Fwd: 5' - [Biotin] - | GCAGCGTCAGATGTGTATAAGAGACAG | - 3' |
| Fwd: 5' - | GTGACTGGAGTTCAGACGT | - 3' |

Table S10: **Protocols and primers used for single cell barcoding and ONT sequencing.**

| Method | Variant | Description |
|---|---|---|
| PCA | - | Unintegrated baseline |
| Harmony | Batch only | Correct batch effects |
| Harmony | Batch + cell cycle | Regress cell cycle, then correct batch |
| scVI | Batch only | Batch as covariate |
| scVI | Batch + cell cycle | Batch and cell cycle as covariates |
| scANVI | Semi-supervised | Adds cell type label guidance |

Table S11: **Summary of data integration strategies evaluated for pseudotime projection.**

| Code | Description |
|---|---|
| SE | Skipping exon |
| A5 | Alternative 5′ splice site |
| A3 | Alternative 3′ splice site |
| MX | Mutually exclusive exon |
| RI | Retained intron |
| AF | Alternative first exon |
| AL | Alternative last exon |

Table S12: **Alternative splicing event types.**
